# Personalized transcriptome signatures in a cardiomyopathy stem cell biobank

**DOI:** 10.1101/2024.05.10.593618

**Authors:** Emma Monte, Takaaki Furihata, Guangwen Wang, Isaac Perea-Gil, Eric Wei, Hassan Chaib, Ramesh Nair, Julio Vicente Guevara, Rene Mares, Xun Cheng, Yan Zhuge, Katelyn Black, Ricardo Serrano, Orit Dagan-Rosenfeld, Peter Maguire, Mark Mercola, Ioannis Karakikes, Joseph C. Wu, Michael P. Snyder

## Abstract

**BACKGROUND:** There is growing evidence that pathogenic mutations do not fully explain hypertrophic (HCM) or dilated (DCM) cardiomyopathy phenotypes. We hypothesized that if a patient’s genetic background was influencing cardiomyopathy this should be detectable as signatures in gene expression. We built a cardiomyopathy biobank resource for interrogating personalized genotype phenotype relationships in human cell lines.

**METHODS:** We recruited 308 diseased and control patients for our cardiomyopathy stem cell biobank. We successfully reprogrammed PBMCs (peripheral blood mononuclear cells) into induced pluripotent stem cells (iPSCs) for 300 donors. These iPSCs underwent whole genome sequencing and were differentiated into cardiomyocytes for RNA-seq. In addition to annotating pathogenic variants, mutation burden in a panel of cardiomyopathy genes was assessed for correlation with echocardiogram measurements. Line-specific co-expression networks were inferred to evaluate transcriptomic subtypes. Drug treatment targeted the sarcomere, either by activation with omecamtiv mecarbil or inhibition with mavacamten, to alter contractility.

**RESULTS:** We generated an iPSC biobank from 300 donors, which included 101 individuals with HCM and 88 with DCM. Whole genome sequencing of 299 iPSC lines identified 78 unique pathogenic or likely pathogenic mutations in the diseased lines. Notably, only DCM lines lacking a known pathogenic or likely pathogenic mutation replicated a finding in the literature for greater nonsynonymous SNV mutation burden in 102 cardiomyopathy genes to correlate with lower left ventricular ejection fraction in DCM. We analyzed RNA-sequencing data from iPSC-derived cardiomyocytes for 102 donors. Inferred personalized co-expression networks revealed two transcriptional subtypes of HCM. The first subtype exhibited concerted activation of the co-expression network, with the degree of activation reflective of the disease severity of the donor. In contrast, the second HCM subtype and the entire DCM cohort exhibited partial activation of the respective disease network, with the strength of specific gene by gene relationships dependent on the iPSC-derived cardiomyocyte line*. ADCY5* was the largest hubnode in both the HCM and DCM networks and partially corrected in response to drug treatment.

**CONCLUSIONS:** We have a established a stem cell biobank for studying cardiomyopathy. Our analysis supports the hypothesis the genetic background influences pathologic gene expression programs and support a role for *ADCY5* in cardiomyopathy.

## Introduction

Hypertrophic cardiomyopathy (HCM) occurs in 1 in 500 individuals, and patient phenotypes range from asymptomatic to serious adverse outcomes such as heart failure or sudden cardiac death.[1] HCM is marked by an enlarged left ventricular muscular wall, with left ventricular ejection fraction typically preserved or increased,[1] whereas dilated cardiomyopathy (DCM) is characterized by reduced ejection fraction.[2] DCM is estimated as the cause of heart failure in ∼12.5 percent of patients and has been estimated to affect 1 in 250 individuals,[2] with familial DCM representing a fraction of those cases.[3] Until recently, the accepted inheritance mechanism for HCM and familial DCM was a single or few dominant, rare mutations, most commonly in genes encoding sarcomere proteins (HCM, especially MYH7 and MYBPC3) [1, 3] or across at least nine key cardiac structures and components (DCM, including sarcomere [TTN] and nuclear envelope [LMNA]).[4] However, the full list of genes proposed to harbor HCM or DCM pathogenic variants exceeds 100 and is continually being refined based on increased genome sequencing data and molecular validation studies (Table S4).[5, 6] For sarcomeric genes known to harbor pathogenic mutations for both HCM and DCM, the opposing effect of the specific mutation on tension generation during cardiomyocyte contraction is thought to distinguish between the development of HCM and DCM.[7] However, despite their contrasting phenotypes, there are shared disease processes between HCM, DCM, and common forms of heart disease, and a shared need for tools to dissect genotype-phenotype relationships. We built a biobank of patient-derived induced pluripotent stem cells for studying disease mechanisms of cardiomyopathy, focused on recruitment for the two most common cardiomyopathies, HCM and DCM.

Both HCM and DCM develop gradually with age and are marked by pathogenic mutations with incomplete penetrance and variable expressivity, and subsequent variability in disease manifestation, with just over half of unaffected individuals harboring a pathogenic variant for HCM remaining unaffected for 15 years post-genetic identification, while first degree relatives of a patient with familial DCM have only a 19% risk of developing DCM by age 80.[3] Furthermore, previous cardiomyopathy subtyping efforts have shown minimal correlation to the underlying gene carrying the pathogenic mutation[8] including broad segregation by patients with known sarcomeric pathogenic mutations or not (HCM).[9] Finally, with few, rare exceptions, actionable changes to clinical care do not exist to tailor treatment based on the mutated gene[10, 11]. Potential explanations for the disparate phenotypes of individuals with a common mutated gene include differences in environment and physiology, nuances of the specific mutation within the gene, and a role for modifying mutations to influence disease onset, severity, and symptomology.

Furthermore, a pathogenic or likely pathogenic mutation is identified for only 30-60 percent of HCM patients[3] and ∼35 percent of DCM patients[4] who undergo clinical genetic testing. This number has remained recalcitrant to expanded application of whole genome sequencing, and replicated in our study as well (see below). This is likely partially explained by the observation that many mutations are family-specific[12] and therefore lack evidence in the literature to support definitive pathogenicity classification. Increasingly though, there is evidence for a subset of patients to have a different genetic architecture. Oligogenic inheritance, where multiple rare variants drive disease, has been proposed for both DCM[4] and HCM[10], as has polygenic inheritance, and the role of modifying mutations to influence disease manifestation in monogenic cases. Genome-wide association studies for DCM, HCM and cardiac morphological and functional traits have revealed individual loci and polygenic risk scores can partially capture cardiomyopathy inheritance, in both patients with and without known pathogenic mutations and in sporadic cases (non-familial DCM).[4, 13, 14]

We hypothesized that if noncoding variants, or variants outside of traditional cardiomyopathy genes influence cardiomyopathy, that should be detectable as signatures in the gene expression. We sought to use inferred, personalized co-expression networks to test whether transcriptomic subtypes exist for HCM and DCM.

## METHODS

The methods are described in the Supplemental Material. This study is in compliance with the Stanford Human Research Protection Program guidelines and approved by the Stanford Institutional Review Board (IRB #30064). In addition, the procedures are in compliance with the International Society of Stem Cell Research guidelines and approved by the Stanford IRB / Stem Cell Research Oversight panel (SCRO #656). The whole genome sequencing and RNA-sequencing data will be made publicly available via dbGaP, with pertinent metadata available in the Supplemental Materials.

## RESULTS

### We established a cardiomyopathy stem cell biobank

We generated a biobank resource for studying cardiomyopathy. We recruited patients exhibiting either hypertrophic cardiomyopathy (HCM), dilated cardiomyopathy (DCM), or left ventricular noncompaction (LVNC) or serving as controls (Figure 1A and Tables 1 and S1). We successfully reprogrammed peripheral blood mononuclear cells (PBMCs) into induced pluripotent stem cells (iPSCs) for 300 donors (Figures 1A and S1). This represented 101 HCM donors, 88 DCM donors, 14 LVNC donors, 95 control donors and 2 donors with other cardiac diseases (long QT syndrome [LQT] and Fabry disease). For most of the samples, echocardiogram measurements of the donor were available in the electronic medical record (EMR) for left ventricular ejection fraction (LVEF, 195/205 diseased iPSC lines) and interventricular septum thickness, end diastole (IVSd, 196/205 diseased iPSC lines) (Table S1).

**Figure 1.**
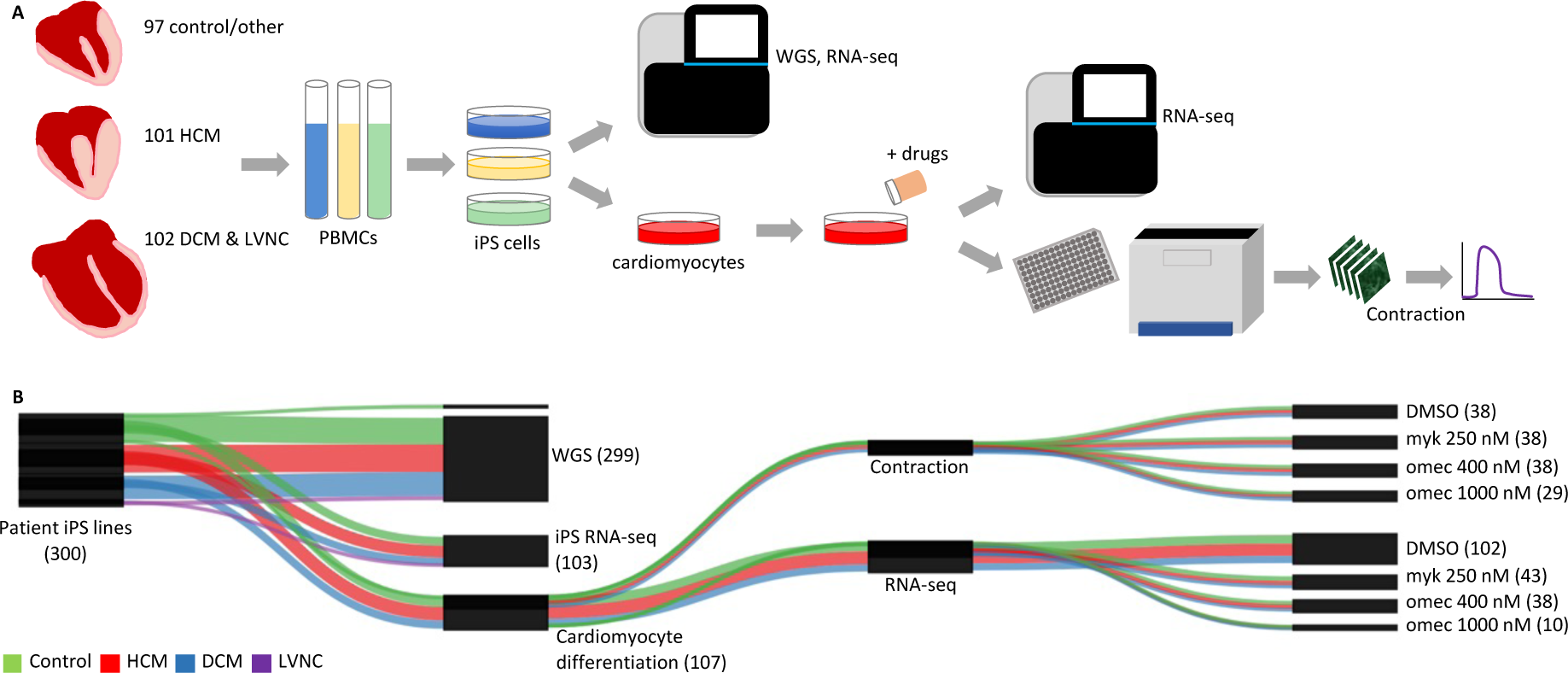
We built a cardiomyopathy stem cell biobank for 300 donors. **A.** We created a biobank of stem cells from 300 donors exhibiting either HCM, DCM, LVNC, or serving as controls. iPS cells were profiled by whole genome sequencing and a subset differentiated into cardiomyocytes for additional profiling via RNA-seq, drug treatment, and microscopy-based contractility assaying. **B.** Plotted are the datasets which passed quality control filtering. Numbers indicate number of donors for each dataset type. The actual number of datasets is higher due to replicates. Abbreviations: hypertrophic cardiomyopathy (HCM), dilated cardiomyopathy (DCM), left ventricular noncompaction (LVNC), induced pluripotent stem cells (iPS cells), whole genome sequencing (WGS), RNA sequencing (RNA-seq), dimethyl sulfoxide (DMSO), mavacamten (myk), omecamtiv mecarbil (omec).

**Table 1.** Demographic and echocardiography metadata. Data are presented as mean ± standard deviation (age/LVEF/IVSd) or as percentage (sex/race/ethnicity). Control includes Healthy Control and Other. Echocardiography data provided where available in electronic medical records. Abbreviations: hypertrophic cardiomyopathy (HCM), dilated cardiomyopathy (DCM), left ventricular noncompaction (LVNC), left ventricular ejection fraction (LVEF), interventricular septum thickness end diastole (IVSd).

|  | Control (n=97) | HCM (n=101) | DCM (n=88) | LVNC (n=14) |
| --- | --- | --- | --- | --- |
| Age | 52.4 ± 18.2 years | 54.4 ± 16.3 years | 50.1 ± 14.6 years | 43.8 ± 17.5 years |
| Male | 54.6 % | 61.4 % | 50 % | 57.1 % |
| Race |  |  |  |  |
| White | 52.6 % | 71.3 % | 63.6 % | 78.6 % |
| Asian | 27.8 % | 10.9 % | 10.2 % | 7.1 % |
| African American | 7.2 % | 5 % | 10.2 % | 7.1 % |
| Other/Unknown | 12.4 % | 12.9 % | 15.9 % | 7.1 % |
| Hispanic | 11.3 % | 8.9 % | 9.1 % | 0 % |
| LVEF (%) |  | 61.6 ± 10.7 (n=93) | 40.0 ± 15.4 (n=87) | 51.7 ± 12.5 (n=14) |
| IVSd (cm) |  | 1.66 ± 0.49 (n=96) | 0.94 ± 0.18 (n=87) | 0.87 ± 0.20 (n=13) |

We performed whole genome sequencing (WGS) on 299 iPSC lines. We differentiated a subset of iPSC lines into cardiomyocytes and profiled them by RNA-seq. Cardiomyocyte transcriptomic data was generated for 102 lines at baseline after quality control filtering (Figure 1B). This represented 44 HCM, 26 DCM, 31 control, and 1 LQT donors for iPSC-derived cardiomyocyte RNA-seq data. We also performed RNA-seq on 103 iPSC lines as a control dataset. A portion of cardiomyocytes were subjected to cardiac drug treatment followed by RNA-seq and contractility measurements using kinetic image cytometry. This resulted in 11-18 drug-treated lines for each disease condition and each drug (mavacamten and omecamtiv mecarbil) after QC.

### Pathogenic variants were annotated via whole genome sequencing

We first sought to identify the pathogenic variants for each iPSC line. Given the open questions around the diversity of the genetic architecture of HCM and DCM, this allowed us to evaluate whether any transcriptomic patterns were evident in both lines with and without known pathogenic mutations. Annotating iPSC lines by their pathogenic variant also enhanced the utility of the biobank as a resource for others.

We performed whole genome sequencing on 299 of the iPSC lines (Tables S2 and S3), and called single nucleotide variants (SNVs) and insertions and deletions (indels). Given the known challenges in identifying pathogenic variants in HCM and DCM our approach was to first filter variants with several different less-stringent criteria and then pool the variants from these different strategies (Figure S2), followed by manual application of the more stringent American Medical College of Genetics (ACMG) guidelines for determining pathogenicity.[15] Identifying pathogenic mutations is dependent on evidence in the literature to support pathogenicity as well as our current understanding of the inheritance model to inform mutation filtering. (We assumed one or several rare dominant mutations in a set of known or potential cardiomyopathy genes (referred to as our “panel genes” [Table S4]) could be pathogenic in any individual). Since both of these may change with time, we provide the specific criteria that we used in Figure S2 and Table S5. Our panel of 235 potential cardiomyopathy genes was purposely broad, encompassing genes from six clinical panels as well as authoritative resources (Table S4, and Supplemental Methods). After applying our initial filters for candidate mutations (pool 1 variants in Figure S2, filtered for frequency <0.001), on average, each iPSC line had only 4-6 rare candidate missense or splicing mutations across the 235 panel genes, and there was no difference between control, HCM, and DCM iPSC lines (Figure 2A), highlighting the challenge of variant classification. Rare candidate truncating, frameshift, or startgain mutations (i.e. mutations potentially altering protein length) were slightly more common in DCM than HCM (0.56 vs 0.33 such mutations per iPSC line).

**Figure 2.**
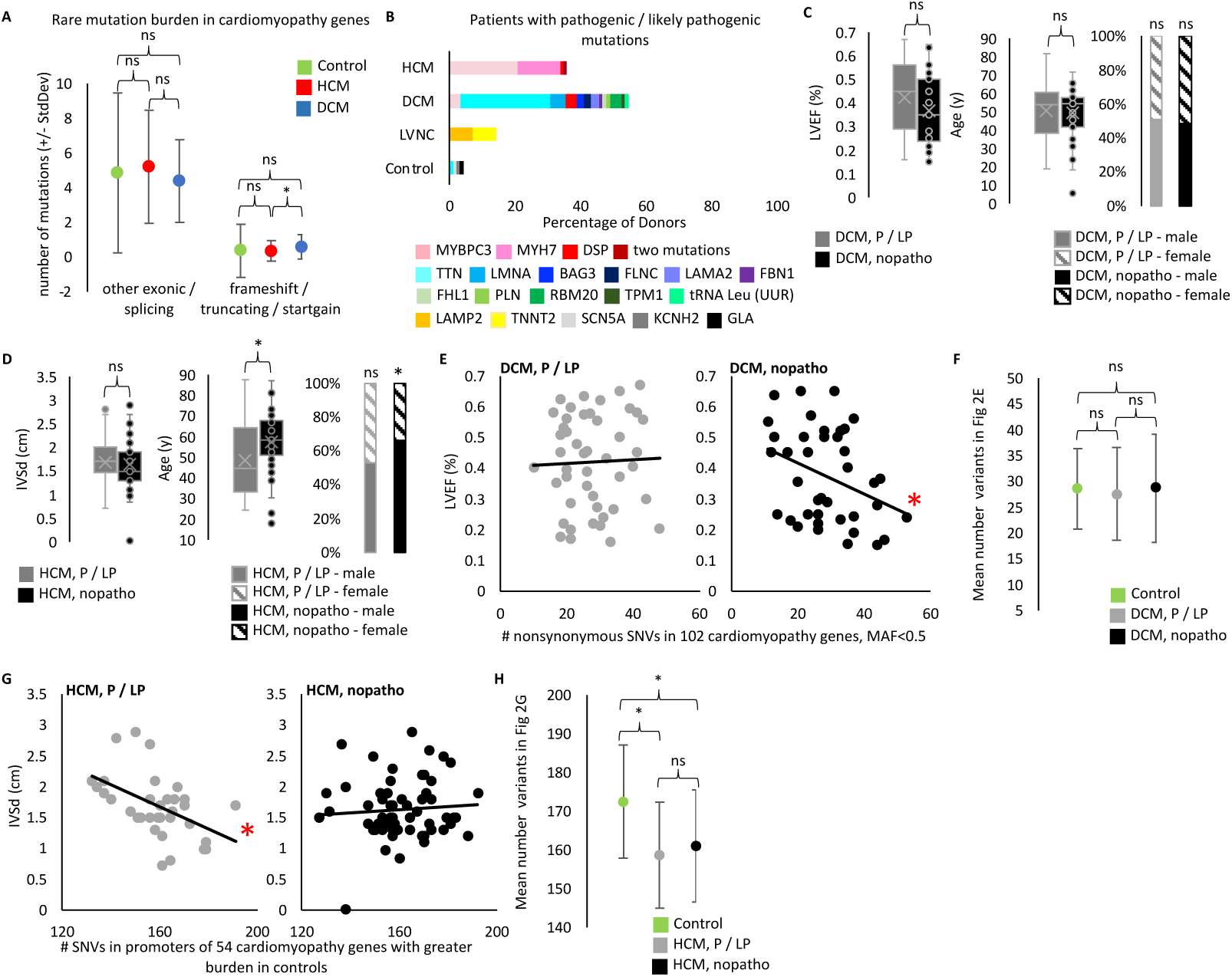
WGS confirms cumulative role of cardiomyopathy variants. **A.** We performed WGS on 299 of the 300 iPS lines and called SNPs and indels. From a starting pool of ∼4.28 million SNPs and indels per donor iPS cell line (mean for biobank, Table S2) we found a mean of only 4-5 missense and splicing mutations per HCM or DCM line and 0.3-0.6 truncating, frameshift, and startgain mutations per HCM or DCM line when we focus on potentially pathogenic variants by filtering for rare variants in 235 cardiomyopathy genes (pool 1 variants in Figure S2, filtered for frequency <0.001). We found control samples show no difference in the number of rare candidate variants in cardiomyopathy genes compared to diseased samples. Rare candidate truncating, frameshift, or startgain mutations (i.e. mutations potentially altering protein length) in cardiomyopathy genes were more common in DCM than HCM. (t-test: DCM vs HCM p-value = 0.0199477. Control vs HCM p-value = 0.9479979. DCM vs Control p-value = 0.2322563). Rare missense and splicing mutations show no difference by disease status. (t-test: Control vs HCM p-value = 0.5390647. Control vs DCM p-value = 0.4026431. HCM vs DCM p-value = 0.05390647). Plotted are the mean number of filtered mutations per line in each disease cohort. Error bars indicated standard deviation. **B.** We identified a pathogenic or likely pathogenic (P/LP) mutation for 86 out 203 diseased iPS lines. Plotted is the percentage of lines with an identified P/LP mutation by gene for each disease category. The HCM line with two mutations has ALPK3 and MYBPC3 mutations. The control line with a GLA mutation was classified as Other due to the donor’s known condition of Fabry disease (an HCM lookalike syndrome). The other two mutations found in control lines are probably not pathogenic in these donors, however we list their finding here as evidence of the background rate of finding pathogenic mutations when applying our filtering and classification workflow to non-cardiomyopathy donors. **C.** We compared echocardiogram and demographic data of the DCM donors with a P/LP mutation identified in the iPS line and those without (referred to as nopatho). Neither LVEF, nor age, nor sex differ between P/LP and nopatho in DCM. (LVEF t-test: p-value = 0.09055781 [p-value 0.08214506 when limit to DCM donors with clinical diagnosis, data not shown]. Age t-test: p-value =0.64370666 [p-value 0.569830699 when limit to DCM donors with clinical diagnosis, data not shown]. Sex chi-square: P/LP male vs female p-value = 0.8864 [p-value 0.6617 when limit to DCM donors with clinical diagnosis, data not shown]. nopatho male vs female p-value = 0.8728 [p-value 0.8694 when limit to DCM donors with clinical diagnosis, data not shown].) **D.** For HCM, P/LP donors are younger (t-test: p-value = 0.00921755), but show no difference in IVSd (t=test: p-value = 0.60639198). Nopatho lines are more commonly male than female, while P/LP lines are equally male and female. (chi-square: nopatho male vs female p-value = 0.0092. P/LP male vs female p-value = 0.7389). **E.** Pucklewartz et al. defined 102 cardiomyopathy genes whose nonsynonymous SNV mutation burden correlated with LVEF in DCM but not with any HCM echocardiogram metrics tested. We found increased cumulative burden of nonsynonymous variants with a minor allele frequency (MAF) <0.5 in the 102 Puckelwartz genes correlated with worse LVEF only in the nopatho samples (right, linear regression: p-value = 0.03141 [p-value = 0.04568 when limit to DCM donors with clinical diagnosis, data not shown; p-value = 0.05929 when remove MAF filter, data not shown.]) but not P/LP samples (left, linear regression: p-value = 0.8093 [p-value = 0.7954 when limit to DCM donors with clinical diagnosis, data not shown. p-value = 0.7514 when remove MAF filter, data not shown]). **F**. However the mean number of nonsynonymous variants in the Puckelwartz et al. genes is not different between the nopatho and P/LP cohorts nor between either DCM cohort and control. (t-test: nopatho vs control p-value = 0.984052497. P/LP vs control p-value = 0.469184129. P/LP vs nopatho p-value = 0.600062881.) Bars represent standard deviation. **G.** We found 54 of the Puckelwartz et al genes had lower mean number of SNVs in the promoter region, in HCM than control. Greater SNVs in this subset of promoters correlated with less enlarged IVSd measurements in the P/LP HCM samples but not the nopatho HCM samples. (Linear regression: P/LP p-value = 0.003773. Nopatho p-value = 0.5669). **H.** As selected for, control samples had greater SNVs in these 54 promoters. There was no difference in the mean SNV count between P/LP and nopatho HCM samples. (t-test: nopatho vs control p-value = 0.00000497. P/LP vs control p-value = 0.00000491. P/LP vs nopatho p-value = 0.439656222.) Bars represent standard deviation.

We identified a pathogenic or likely pathogenic mutation in 36 percent of HCM lines (36 of 101), most commonly in MYH7 or MYBC3 (13 and 22 iPSC lines, respectively), and in 55 percent of DCM lines (48 of 88), most commonly in TTN (24 iPSC lines; Figure 2B and Table S5). Complementary RNA-seq data from iPSC-derived cardiomyocytes was helpful for evaluating potentially truncating variants, but ultimately did not change our annotation of a mutation from a variant of uncertain significance to pathogenic or likely pathogenic (Figure S3A). 145 of the cardiomyopathy donors had clinical genetic testing results in their EMR. We re-evaluated pathogenicity for any variant listed in the EMR. The rate of identifying a pathogenic or likely pathogenic mutation by whole genome sequencing was only slightly higher than by clinical testing (45.5 percent of lines versus 41.4 percent) (Figure S3B and Table S5). In total, the biobank contains diseased iPSC lines for 21 different mutated genes.

Reduced LVEF is a hallmark of symptomatic DCM and an indicator of poor cardiac function. We saw no difference in the LVEF, age, or sex of DCM donors for which we found a pathogenic or likely pathogenic mutation compared to those without a known pathogenic mutation (Figure 2C). Similarly, we saw no difference in IVSd, one measure of cardiac size, between HCM donors with or without pathogenic and likely pathogenic mutations (Figure 2D). However, we found donors lacking pathogenic mutations were older and more commonly male than female, while HCM donors with a known pathogenic mutation were equally male and female (Figure 2D).

### Correlation between mutation burden in cardiomyopathy genes and echocardiogram metrics was distinct between lines with and without known pathogenic mutations

Pucklewartz et al.[16] had previously evaluated mutation burden in 102 cardiomyopathy genes (101 of which are in our “panel genes” list) focusing on nonsynonymous SNVs of any allele frequency and found greater nonsynonymous SNVs correlated with decreased LVEF in DCM but not HCM patients. The authors proposed a role for oligogenic inheritance to contribute to DCM phenotype. By contrast, we found neither our HCM nor DCM cohort displayed this relationship (Figure S4A). However, Pucklewartz et al. specifically enriched for donors without known pathogenic or likely pathogenic mutations in building their cohort. When we distinguished between iPSC lines with a pathogenic or likely pathogenic mutation and lines without, and limited the analysis to variants with an alternate allele frequency <0.5, we found the linear relationship between mutation burden and LVEF is specific to our DCM cohort lacking pathogenic mutations (Figure 2E). Importantly, there was a large range in the mutation burden of nonsynonymous SNVs in the 102 cardiomyopathy genes from 10 to 43 variants across the samples (DCM and control), with no difference between control, DCM with a pathogenic or likely pathogenic mutation, and DCM without a pathogenic or likely pathogenic mutation (Figure 2F). The significance was specifically related to the correlation between the mutation burden and LVEF. This supported the hypothesis in the literature for two different mechanisms of DCM inheritance; the DCM samples with pathogenic mutations exhibiting monogenic inheritance and the DCM samples without known pathogenic mutations potentially exhibiting oligogenic inheritance. Importantly, when we restricted the analysis to a subset of 20 core DCM genes with stronger evidence for pathogenicity (see Supplemental Methods), we no longer saw a correlation between mutation burden and LVEF, further supporting the hypothesis of the Puckelwartz et al. authors for oligogenic inheritance (linear regression: nopatho p-value = 0.06238. P/LP p-value = 0.8751; data not shown). Here we applied the Puckelwartz et al. analysis from 2021 to our cohort, but additional DCM GWAS datasets and machine learning approaches should enable improved selection of loci for which mutation burden could contribute to DCM. Importantly, this analysis does not suggest these particular mutations (coding mutations in 102 cardiomyopathy genes) act as modifier mutations, in that they do not correlate with LVEF in the donors with a known pathogenic mutation.

Because the Puckelwartz et al. study examined multiple echocardiogram measurements for correlation to mutation burden in HCM and did not find any, we did not pursue this line of investigation, except to confirm the lack of correlation to LVEF as a complement to our DCM analysis. Instead, we performed only one complimentary analysis, which was to compare the IVSd data we had for HCM to mutation burden in the promoters of the Puckelwartz et al. genes. Our hypothesis was that the same genes which may contribute to oligogenic inheritance in DCM could act as modifiers of disease severity in HCM, in which case, non-coding mutations may be sufficient to influence phenotype. While the majority of pathogenic cardiomyopathy mutations influencing protein sequence are harmful, we could not assume whether individual non-coding mutations would be harmful or protective, nor were we powered to determine this statistically. Rather, we distinguished promoters by those with greater mutation burden (SNVs of any allele frequency) in control samples and those with greater mutation burden in HCM samples. Of the 102 promoters, 54 had greater average mutations in control versus HCM lines. Total mutation burden in these 54 promoters was significantly associated with lower IVSd in HCM lines with pathogenic mutations (Figure 2G). The trend remained significant when only analyzing data from white donors (p-value = 0.01004, data not shown). We did not see this relationship in the HCM samples without a pathogenic mutation (Figure 2G). The difference between HCM samples with and without a pathogenic variant was not due to a difference in total mutation burden between these two groups (Figure 2H), but the specific relationship between mutation burden and IVSd. Our limited sample size for selecting and analyzing variants meant these results were not sufficient for making conclusions about the role of these promoters in HCM. Rather, this provided preliminary evidence for our HCM cohort to encompass diverse genetic architecture mechanisms. We thus sought to investigate transcriptomic signatures as a molecular phenotype of cardiomyopathy that may be sensitive to distinct genetic backgrounds. Importantly, we were not attempting to replicate cardiac expression quantitative trait loci (eQTL) studies.[17] Rather, we hypothesized that leveraging a diverse human dataset could uncover important cardiomyopathy disease mechanisms by specifically interrogating shared and personalized transcriptomic features across patients of differing genetic architectures.

### Disease co-expression networks identified important cardiomyopathy genes, with *ADCY5* as the largest node for both HCM and DCM

iPSC lines were differentiated into cardiomyocytes and profiled via RNA-seq (Table S6) resulting in RNA-seq data from iPSC-derived cardiomyocytes for 102 subjects after QC. We performed traditional differential gene expression analysis using DESeq2 and identified 236 and 62 genes up an down-regulated respectively in HCM and 8 and 21 genes up and downregulated in DCM with gene ontology analysis resulting in few disease pathways (Supplemental Methods, data not shown), likely due to the diverse genetic etiology of the cohort and our limited ability to perform multiple cardiomyocyte differentiations for each of the over 100 iPSC lines. We also expected that for some samples the presence of a pathogenic mutation would not guarantee that the iPSC-derived cardiomyocytes would be mature enough, nor the model stressed enough, to bring out a phenotype for that specific mutation, and we thus lacked a true positive set of samples for building a definition of diseased expression. Nor did we want to build a unique model of cardiomyopathy expression for each sample to accommodate the heterogeneity of symptoms from different pathogenic mutations. Instead, we sought to identify common transcriptional signatures of cardiomyopathy and evaluate each sample by the manifestation of the shared signatures. We focused on gene co-expression relationships based on the supposition that the influence of genetic architecture and noncoding variants may be better captured.

We calculated patient-specific gene co-expression networks using lionessR, an algorithm for linear interpolation to obtain network estimates for single samples.[18, 19] First, for HCM and DCM separately, we built a co-expression network with the 200 most differential gene-gene co-expression relationships calculated between the control and diseased cohort (Figure 3A). A red edge in the HCM network indicated two genes were highly co-expressed (large r^2^, pearson) with a positive correlation (positive r, both genes up or down expressed similarly across samples) in the HCM cohort compared to the control cohort. By contrast green edges indicated strong, positive co-expression in the control cohort. Separately, we built a DCM network comparing DCM co-expression with control samples. We then used lioness to remove one sample from the cohort, and recalculate the co-expression correlations. The change in the level of co-expression upon sample removal was used to infer the individual contribution of that sample to the network. We thus generated inferred networks for each sample individually. We then asked how the sum of all edges surrounding a gene (node strength) varied across patients (Figure 3B).

**Figure 3.**
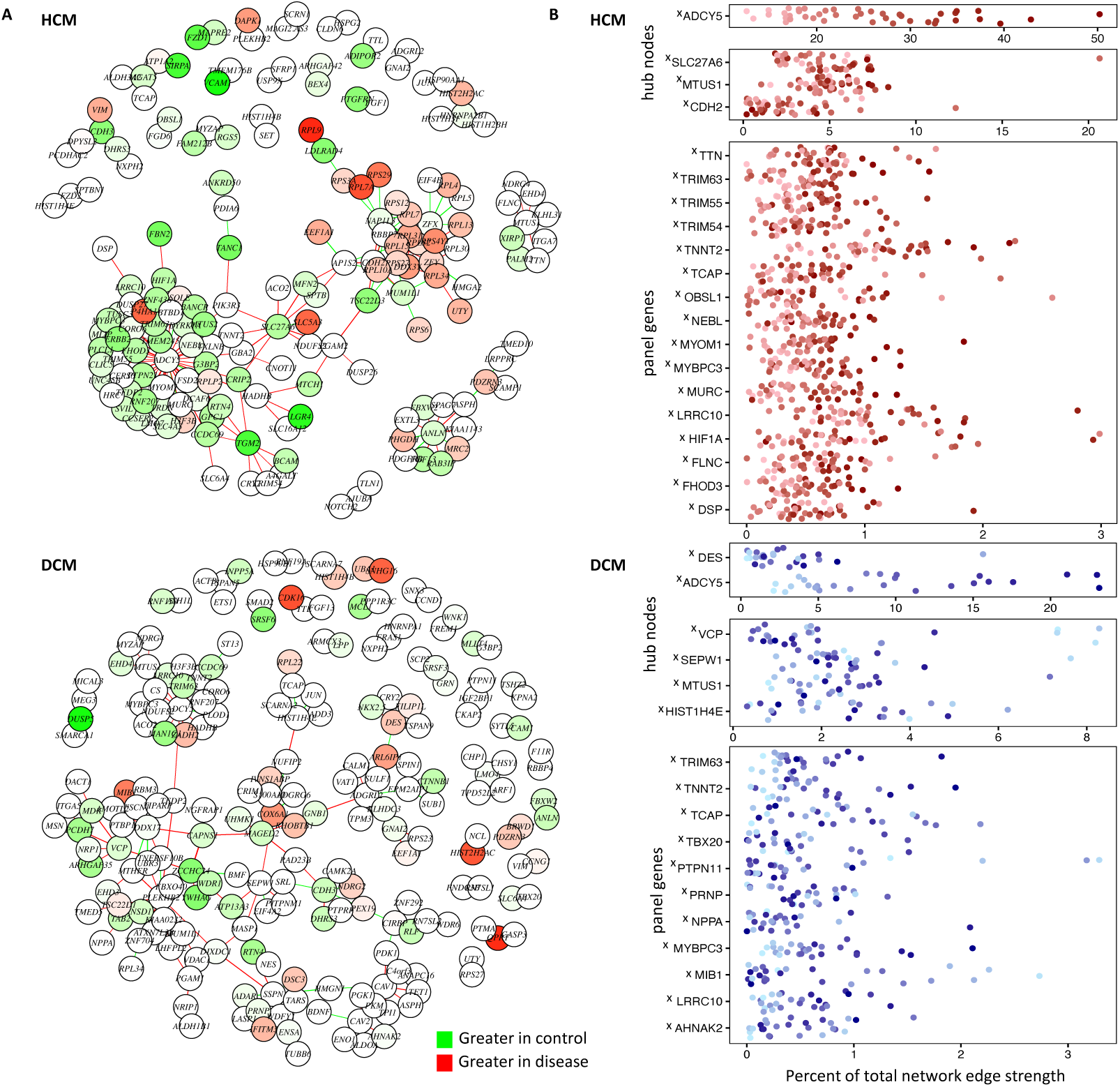
Personalized co-expression networks capture otherwise undetectable genes contributing to the disease signature and reveals line-specific differences in network activation. **A.** We calculated an HCM and DCM co-expression network using lionessR, an algorithm for Linear Interpolation to Obtain Network Estimates for Single Samples. (Genes/nodes are represented as circles. Edges are represented as lines. Green indicates stronger edges or greater expression [nodes] in control, while red indicates stronger edges/greater expression in disease.) **B.** We the inferred personalized co-expression networks for individual lines. For select genes we highlight their contribution to the network (sum of their edge strengths as a percentage of the total edge strengths of the network), plotted for each sample. Samples are colored by their *ADCY5* ranking. X indicates genes which only show up in this lines-specific co-expression analysis but are not flagged as significant in traditional DESeq2 analyses for differential expression between control and disease. Panel genes refers to cardiomyopathy gene list used to annotate pathogenic variants.

Of our “panel genes” that we screened for pathogenic variants, 16 were in the HCM network (plus *CDH2*, which had been shown to be mutated in arrhythmogenic right ventricular cardiomyopathy[20], but not HCM or DCM) and 12 were in the DCM network. Despite not showing up in our traditional differential gene expression analysis as exhibiting a conserved difference in gene expression across the disease cohort, we saw they exhibit disease-specific co-expression. Other genes of interest in the HCM network included *SLC27A6* which encodes fatty acid transport protein 6 (FATP6), the primary FATP in the heart.[21] FATPs enable cellular uptake of fatty acid, with fatty acid oxidation being the dominant source of ATP in healthy adult hearts, while classic pathologic transcription remodeling via the “fetal gene program” entails a switch to other substrates.[21] *SLC27A6* was previously identified in an exome-wide association study for association with left ventricular internal diastolic dimension in the Hypertension Genetic Epidemiology Network of paired siblings with and without hypertension.[21] *MTUS1* was in both the HCM and DCM network. *Mtus1A*, a MTUS1 splice variant, was shown to be upregulated in a murine model of pressure overload with corresponding increase in cardiac hypertrophy, while overexpression attenuated hypertrophy in response to pressure overload and catecholaminergic stimulation.[22] *JUN*, also found in both the HCM and DCM networks encodes a transcription factor with a known role in regulating sarcomere gene expression and attenuating cardiac hypertrophy.[23]

In DCM, we found additional examples of genes previously implicated in cardiomyopathy. *VCP* is a molecular chaperone with roles in mitochondrial maintenance and protein homeostasis whose overexpression or disrupted function in mice can moderate ischemia reperfusion injury and heart failure respectively.[24] *HIST1H4E*, (encoding Histone H4) was previously identified for differential expression in cardiomyopathy and cardiomyopathy risk factors in microarray datasets.[25] We also identified genes previously understudied in cardiomyopathy, such as *SEPW1*. Selenium deficient disruption of selenoprotein function has been implicated in heart failure,[26] but little is known of a specific role for *SEPW1* in DCM. In total, we saw the personalized co-expression analysis allowed for interrogation of individual genes in a sample-specific manner, as well as capturing otherwise undetectable genes contributing to the disease transcriptome.

Network activation is a measure of the total strength of all edges in the network. High network activation in a diseased sample meant the sample exhibited strong disease-specific gene co-expression. Likewise, an activated hubnode represented a gene with strong co-expression relationships in a sample. In both HCM and DCM, the *ADCY5* gene was the largest node (connected to the most other genes) and had the largest contribution to the total strength of the network (Figure 3A). The prominence of *ADCY5* in both the HCM and DCM networks indicated *ADCY5*, despite not being a gene mutated in cardiomyopathy, was co-expressed with multiple cardiomyopathy genes and central to the disease networks. Mouse models have demonstrated the role for *ADCY5* perturbation to influence other forms of heart disease,[27] with our network data suggesting *ADCY5* may also be important to cardiomyopathy. Importantly, the relative contribution of the *ADCY5* hubnode to the level of network activation was highly variable between lines (Figure 3B), prompting us to next examine how differences in network activation related to disease severity.

### Personalized networks illuminated distinct relationships between network activation and disease severity

Having confirmed the utility of co-expression analysis for identifying cardiomyopathy genes of interest, we next tested whether the network itself offered disease insights. We defined a hubnode as a gene with at least three edges and asked whether edges around a shared hub node were further co-modulated, signifying the hub node itself was a unit of network activation (Figure 4A). Put simply, if we found in one of our HCM samples that the inferred co-expression relationship between *ADCY5* and another gene (for example *MYBPC3*) was strong, could we expect *ADCY5*’s co-expression relationship with the other 48 genes it is connected with to also be stronger in that HCM sample as compared to the other HCM samples. This was calculated separately on the two networks (HCM and DCM). Importantly, we calculated this for the control samples separate from the diseased samples, such that we could compare how network activation presented differently for each sample despite having the same disease.

**Figure 4.**
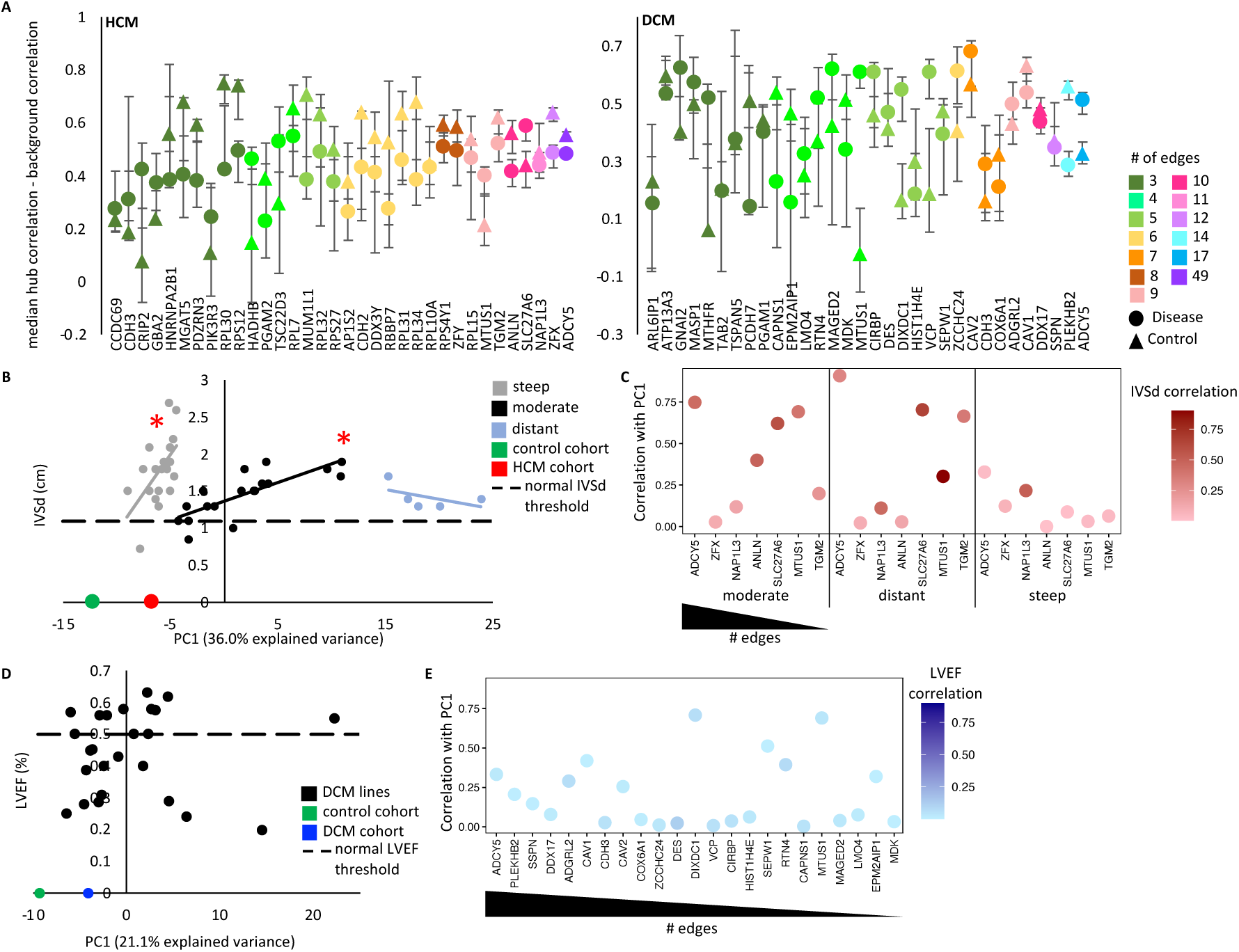
Variable *ADCY5* hub node activation corresponds to clinical disease severity in HCM, with two HCM subgroups exhibiting distinct patterns of network activation. **A.** Co-modulation of edges around a node was evaluated by comparing the difference in correlation between strength of two randomly selected edges around the node with two randomly selected edges that do not share a node. Correlation was evaluated separately on the diseased and control cohorts. Plotted are the mean and 95% confidence intervals after sampling 10,000 times. **B.** Principal component analysis of the HCM network was computed on the control-cohort, HCM-cohort, and individual HCM lines. Principal component 1 (PC1) is plotted against IVSd (intraventricular septal thickness end diastole) for the 42 HCM samples with echocardiogram data. For PC1 values -9.02 to -4.4 (gray dots, “steep” samples) and for PC1 -4.3 to 11 (black dots, “moderate” samples), PC1 correlates with IVSd (linear regression: steep p-value = 0.01167, moderate p-value = 0.0001398). The 5 most distant samples (PC1 >15.2, blue dots) show no relationship to IVSd (p-value = 0.3729). **C.** For hubs with 8 or more edges, we tested how well they served as a proxy for the overall PC1 score. Plotted are the correlation (R2) of the sum of all edge strengths around a hub with the PC1 score for the sample in the HCM cohort (y-axis). Color, indicates the R2 correlation of the sum of edge strengths to IVSd. **D.** There was no significant relationship between PC1 of the network and LVEF in DCM. **E.** Plotted are correlation between node strength and PC1 for nodes with 4 or more edges in DCM.

In the HCM network we saw *ADCY5* was a unit of network regulation, in that for both the control and HCM lines there was greater co-modulation of *ADCY5* edges than background co-modulation of two unconnected edges in the network. Despite many of the individual edges around *ADCY5* being stronger in HCM (red edges in Figure 3A top panel), the co-modulation of the *ADCY5* edges was lower in HCM (HCM node lower than control node in Figure 4A), suggesting *ADCY5* was more activated in HCM, but with individual edges being sporadically activated depending on the sample. In the DCM network, the opposite was true. Like with HCM, *ADCY5* had mostly stronger edges in DCM compared to control, however, the co-modulation was also stronger in DCM than control, suggesting the entire *ADCY5* hub was upregulated in tandem in DCM samples, to varying degrees. Many additional nodes also behaved as significant units of network activation. Notably, DCM had five nodes with greater co-modulation in diseased samples. *MTUS1* showed the largest difference, with co-modulation of the edges around *MTUS1* showing no correlation in the healthy cohort. Conversely, HCM had only one such node (*SLC27A6*), suggesting the level of network activation in HCM samples was not a singular feature, rather the genes being most activated in the network were sample dependent.

For our next analysis of the HCM network, we examined the inferred co-expression values for each HCM sample. We also included the composite values for the HCM cohort as a whole and control cohort as a whole. Principal component analysis was applied and principal component one (PC1) compared to IVSd (Figure 4B). As expected, the PC1 value for the control cohort was the most distant from all the other samples (PC1 = -12.26). Surprisingly, we saw a significant relationship between PC1 and IVSd in the individual HCM samples. For PC1 values closer to control (PC1 -9.02 to -4.4), we saw a linear relationship, where greater distance from control, corresponds to enlarging hearts, with a steep linear trendline. We called these “steep” samples. The linear relationship then reset (PC1 -4.3 to 11) with a moderate linear trendline where greater distance again corresponded to enlargement of the heart. We called these “moderate” samples. The five most distant samples (PC1 >15) showed no relationship to IVSd. This observation provided confidence that the gene expression relationships captured in our network analysis of iPSC-derived cardiomyocytes reflected aspects of the biology of the donor heart and furthermore was measuring critical components of pathologic gene expression remodeling indicative of disease severity. However, while PC1 was useful as a singular indicator value to represent the full disease co-expression network, it was harder to interpret biologically. We next evaluated if individual genes could also be indicators of the network activity.

For hub genes with eight or more edges, we tested how well they served as a proxy for the PC1 value (Figure 4C). We found the moderate and distant groups showed a high correlation between *ADCY5* node strength and PC1, which was expected as it was the gene with the most co-expression pairs (25%, 49 out of 200 edges) and therefore likely drove the largest variability of network strength. Correspondingly, *ADCY5* node strength, like PC1, exhibited significant correlation to IVSd in moderate samples. However, steep samples exhibited a weak correlation between *ADCY5* with either PC1 or IVSd. Notably, no other node showed greater correlation to PC1 in steep samples than *ADCY5* (even when checking all 34 nodes with a minimum of 3 edges [versus nodes with a minimum of eight edges], data not shown).

From the data in Figures 4B and 4C, we drew the following conclusions. Firstly, we identified two distinct HCM groups based on transcriptional behavior. (We focused on the moderate and steep groups as the distant group was only comprised of five samples.) Secondly, both groups encompassed a spectrum of disease severity (range of IVSd values). Thirdly, they shared a common disease co-expression network, such that for both groups network activation levels corresponded to disease severity of the donor (though notably smaller PC1 values were sufficient to indicate high IVSd values for steep samples). Fourth, the groups were distinguished by the manner with which they activated the disease network (even when comparing samples with similarly severe IVSd measurements). Specifically, in moderate samples, *ADCY5* activation was occurring as a unit, such that a moderate HCM sample that exhibited a stronger co-expression relationship between *ADCY5* and one of its paired genes was likely to also have stronger co-expression relationships for all of the gene-gene pairs in the network and to have a correspondingly larger heart (IVSd). By contrast, a steep HCM sample with a large heart (IVSd) was expected to also have a high level of network activation relative to other steep samples, but this would be driven by only specific gene-gene pairs exhibiting strong co-expression, with the specific genes depending on the sample. The observation that network activation correlated to disease severity for both HCM-moderate and HCM-steep groups highlighted the importance of the network genes to HCM. The observation that the manner of network activation differed between the groups, even for samples with similar echocardiogram measurements, suggested the difference between steep and moderate samples was not due to differences in disease severity driven by a single pathogenic mutation but could be due to genetic background.

A similar analysis of the DCM network revealed no significant relationship between PC1 and LVEF (Figure 4D). Like what we found for the HCM steep samples, in the DCM samples no single node served as a good proxy for the whole network (Figure 4E).

We looked for experimental features which could explain the segregation of HCM samples into steep and moderate categories. Returning to the principal component analysis of the network, we found PC3 values partially segregated the steep and moderate HCM samples (Figure S5A). While *ADCY5* edges were the largest contributor to PC1, *ANLN* edges represented the edges with the individual greatest relative contribution to PC3 (Figure S5B). We found that HCM lines with pathogenic mutations were more common in the steep group, especially for female lines (Figure S5C), and that *ANLN* node strength was weaker in steep samples as well as female samples with pathogenic mutations (Figure S5D). *ANLN* had been shown to turn on in mitotic cardiomyocytes,[28] a process not typical of the adult heart, and thus further testing is needed to determine if *ANLN* is a feature in the donor hearts, or only in the iPSC-cardiomyocyte model.

### The RNA network findings and promoter mutation analyses provided mutual validation, supporting characterization of HCM subtypes

Given that moderate samples showed cohesive activation of the HCM network, we wondered if this signified a partially shared genetic background mechanism. We re-examined our analysis of the Puckelwartz et al. genes. We had preliminarily found that the mutation burden in the promoters of 54 of the Puckelwartz et al. genes was correlated with smaller IVSd in samples with a known pathogenic or likely pathogenic mutation, but not in samples without a known mutation. In fact, we now saw that for moderate samples, both samples with and without pathogenic mutations exhibited this correlation (Figure S5E), while steep samples analyzed on their own did not exhibit this correlation, even amongst those with a pathogenic mutation (Figure S5F). Given that our promoter analysis of the Puckelwartz et al genes was underpowered to draw meaningful conclusions and could represent spurious correlations, we applied a published polygenic risk score for HCM.[29] We saw no difference in the average risk of HCM-steep samples versus control samples. However HCM-moderate samples had significantly higher scores than both control samples and HCM-steep samples (Figure S6G). Further, we found that moderate samples, but not steep samples, exhibited the expected phenomenon whereby the donors with a pathogenic mutation were younger than those without (Figure S6H). Taken together, these data supported our hypothesis that moderate HCM samples represented a subgroup of HCM where shared genetic background mechanisms may be influencing both disease severity and the transcriptional phenotype.

### *ADCY5* dysregulation was a shared feature of both HCM and DCM and partially corrected with drug treatment

Further investigation of the importance of *ADCY5* to the HCM network, revealed *ADCY5* node strength explained the vast majority of the network activation in moderate samples and to a lesser extent in distant samples (Figures 5A and 5B), with stronger *ADCY5* co-expression relationships in samples with greater network activation. Whereas, *ADCY5* was only minimally activated in steep samples with minimal variability between samples as well (Figures 5A and 5B). Importantly, *ADCY5* expression showed no difference between the HCM subgroups nor between HCM and control (Figure 5C). This highlighted the value of the network analysis to uncover important pathologic transcriptional remodeling features, but also meant unfortunately investigating future samples could not be done by simply measuring *ADCY5* expression in the absence of co-expression analysis. Despite HCM and DCM hearts exhibiting contrasting phenotypes, we found *ADCY5* was also important in DCM. Increased *ADCY5* node strength compared to control was a shared feature DCM samples, and this was true for both samples coming from donors with normal LVEF (50% or greater) and those with reduced or moderately reduced LVEF (less than 50%) (Figure 5C). Taken together with our previous observation that *ADCY5* node activation did not correlate with total network activation in DCM (Figure 4E), this showed that the *ADCY5* hub node was being universally activated in the DCM samples. *ADCY5* had co-expression relationships with 49 genes in the HCM network and 17 genes in the DCM network. 10 genes were common to both HCM and DCM. Gene ontology analysis of these genes revealed enrichment for the sarcomere (Figure 5D). For HCM specifically, gene ontology analysis also returned 65 significantly enriched transcription factor motifs, the most significant being for MEF2A. MEF2A is a transcription factor with a central role in driving cardiac hypertrophy.[30] We found that in moderate samples but not in steep samples, *MEF2A* expression and *ADCY5* expression is highly correlated (Figure 5E).

**Figure 5.**
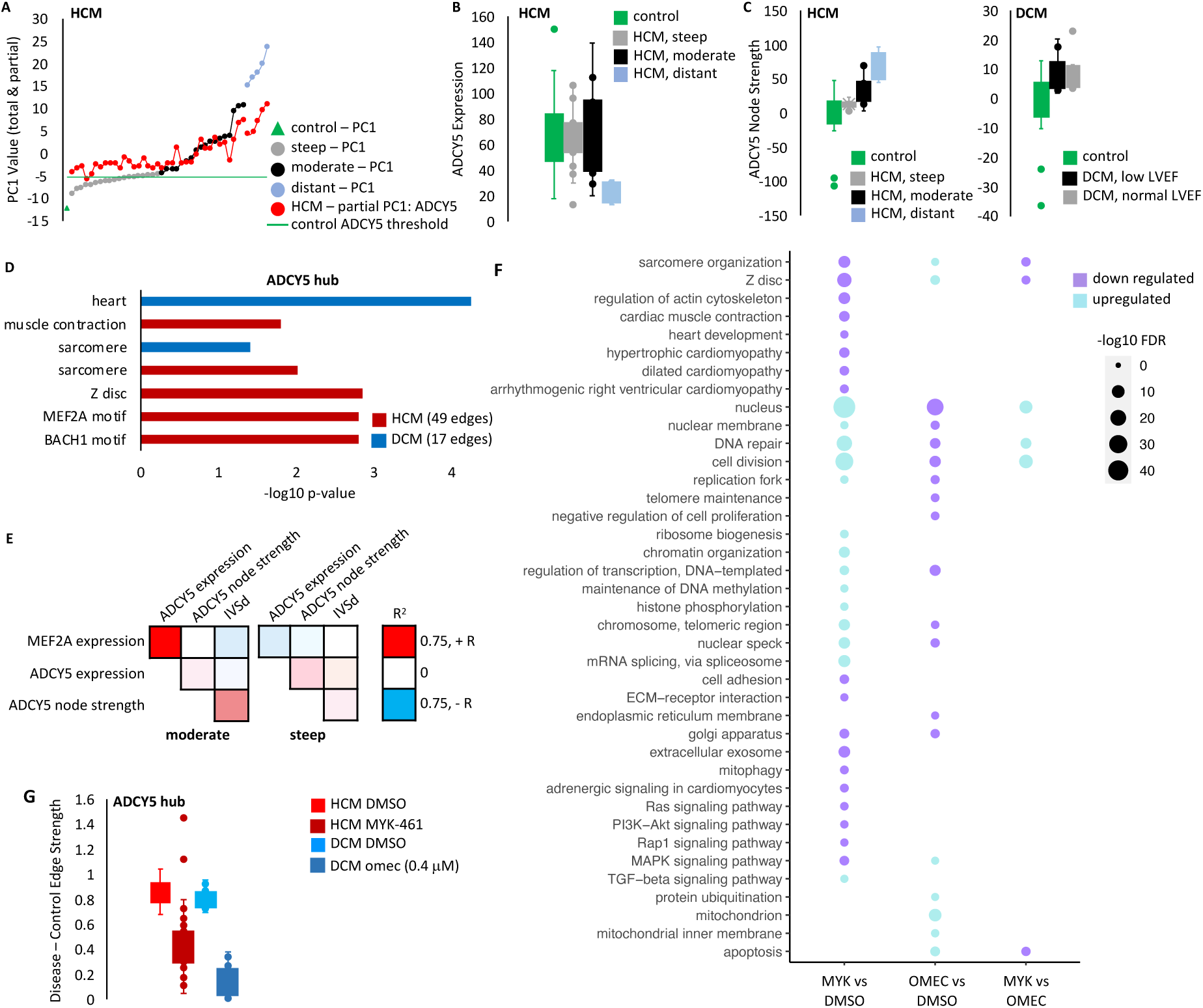
Treatment with MYK-461 or omectamtiv mecarbil partially corrects *ADCY5*. **A.** Plotted is the principal component data from Figure 4B. PC1 for each HCM sample is shown (gray, black, blue, by HCM subgroup). PC1 is a sum of values for each edge. Also plotted is the sum of the scores for all *ADCY5* edges specifically. HCM-moderate samples show *ADCY5* scores increasing with PC1, while ADCY5 scores are similar across HCM-steep samples. **B.** ADCY5 expression is similar between control, moderate and steep samples, but ADCY5 node strength **(C)** is increased in HCM and DCM. **D.** Gene ontology analysis of genes sharing edges with *ADCY5.* **E.** Pearson correlation values between *MEF2A* expression and *ADCY5.* **F.** Cardiomyocytes were treated with the small molecule sarcomere activator (omecamtiv mecarbil) or inhibitor (mavacamten) at 0 hours and 24 hours, with RNA harvested at 48 hours for RNA-seq analysis. Gene ontology analysis revealed many shared drug targets. **G.** For the *ADCY5* node, mean edge strength in the DMSO-treated control cohort was compared with the mean strength in the DMSO-treated or drug-treated disease cohort. Plotted are the difference for each edge around the node in the respective comparisons. In both cases, we see a partial correction (smaller difference vs control) with drug treatment.

Finally, mavacamten (known commercially as Camzyos), is a small molecule inhibitor of MHY7 for treating patients with obstructive HCM that.[31] We treated cardiomyocytes with both mavacamten as well as an MYH7 inhibitor, omecamtiv mecabril[32] for 48 hours and then performed RNA-seq. Kinetic image cytometry was used to visually measure cellular deformation over time and confirm the treatment strategy successfully altered contractility. As expected, mavacamten reduced contractility, while omecamtiv mecarbil increased contractility (Figures S6A and S6B). Gene ontology analysis of RNA-seq after drug treatment revealed many shared drug targets between mavacamten and omecamtiv mecarbil, such as an opposing effect on expression of Z disc components (Figure 5F). For each edge around *ADCY5*, we compared the mean edge strength in the diseased cohort to the control cohort before and after drug treatment and found drug treatment partially corrected the *ADCY5* node for both HCM and DCM (Figure 5G).

## DISCUSSION

We identified *ADCY5* as a central hub node in both the HCM and DCM diseased networks. Adenylyl cyclases catalyze ATP to cAMP conversion, with *ADCY5* and *ADCY6* being the major isoforms in the heart.[27] ADCY5 is sensitive to and able to influence contractile regulation. Beta-adrenergic stimulation and PKC activate ADCY5 which in turn catalyzes cAMP formation, driving PKA signaling. PKA phosphorylation and local calcium levels inhibit ADCY5.[33] Previous studies support a role for *ADCY5* in heart disease. Adenylyl cyclases drive the increased inotropy and lusitropy induced by beta-adrenergic agonist stimulation of the heart by producing cAMP which activates downstream pathways of protein kinase A.[34] In mice, *ADCY5* overexpression increases oxidative stress and worsens cardiomyopathy outcome under chronic stress conditions, while *ADCY5* knockout is protective in chronic stress conditions and a high fat diet model of diabetic cardiomyopathy.[27] Furthermore *ADCY5* knockout mice have increased lifespan, and blunted aging-associated left ventricular hypertrophy and cardiomyopathy.[27] In mice and rabbits, pharmaceutical inhibition of *ADCY5* shortly after coronary artery reperfusion reduced myocardial infarct size.[35] Alternately, G_αq_ overexpression-induced cardiomyopathy mice have decreased *ADCY5*, and further *ADCY5* knockout is not protective.[27] In silico analysis of HCM and DCM identified ADCY5 as a potential drug target for modulating other disease processes.[36] We found *ADCY5* activation was a universal feature of DCM lines, while serving as a biomarker of network activation and donor disease severity for a subgroup of HCM. Importantly, only 10 edges were shared between the *ADCY5* node in HCM (49 edges) and DCM (17 edges). These included contractile genes *MYBPC3*, *TNNT2*, *TRIM63*, and regulators of excitation and excitation-contraction coupling *RFN207* and *LRRC10*[37, 38], with the nodes as a whole enriched for sarcomere constituents (Figure 5D). Here we demonstrated increased *ADCY5* activation in multiple genetic backgrounds from both HCM and DCM, and in the context of disparate pathogenic mutations in a human cell-line model. We further show *ADCY5* node activation is sensitive to contractility modulation through drug treatment and posit it may be sensitive to pathogenic mutations in contractile proteins. In turn, we propose ADCY5 represents a shared molecular phenotype that can influence molecular remodeling downstream of contractile dysfunction, and that targeting ADCY5 may be able to influence contractile dysfunction stemming from multiple etiologies.

Additionally, we confirmed and expanded on the Puckelwartz et al observation of cumulative mutation burden in cardiomyopathy genes to correlate with DCM severity finding in our cohort the relationship is specific to samples without known pathogenic mutations. This supports the hypothesis for distinct DCM inheritance mechanisms and highlights the need for further studies which can properly delineate the risk loci responsible, as it is understood many of the cardiomyopathy gene variants used in this analysis likely do not contribute.

Finally, we characterized individual samples by RNA signatures. For DCM we found individual hub genes represented units of diseased network activation (Figure 4A). However, the relative degree of activation of separate hub genes varied by sample (Figure 4E). Thus the network constituents are important indicators of disease biology and may represent conserved candidates for therapeutic intervention (including *ADCY5*, Figure 5C), but additional RNA signatures are needed to explain disease severity. In HCM, we defined a single diseased transcriptional network with applicability to distinct HCM subgroups, in that for all subgroups, network activation corresponded to more severe echocardiogram measurements of the donor. We interpret the differences in the moderate and steep RNA subtypes as indicative of distinct genetic backgrounds. These data represent preliminary evidence for genetic background to influence molecular phenotype in cardiomyopathy.

## Acknowledgments

We wish to thank the patients who contributed to our cardiomyopathy biobank, making this research possible.

## ARTICLE INFORMATION

### Nonstandard Abbreviations and Acronyms

ADCY5: adenylyl cyclase type 5
DCM: dilated cardiomyopathy
DMSO: dimethyl sulfoxide
EMR: electronic medical record
HCM: hypertrophic cardiomyopathy
indels: insertion and deletions
iPSC: induced pluripotent stem cell
IVSd: interventricular septum thickness, end diastole
lioness: linear interpolation to obtain network estimates for single samples
LVEF: left ventricular ejection fraction
LVNC: left ventricular noncompaction
MAF: minor allele frequency
myk: mavacamten / MYK-461 *\*this is an unconventional abbreviation created for labeling figures*
nopatho: lines without a known pathogenic or likely pathogenic mutation *\*this is an unconventional abbreviation created for labeling figures*
omec: omecamtiv mecarbil *\*this is an unconventional abbreviation created for labeling figures*
P/LP: pathogenic or likely pathogenic mutation PBMC – peripheral blood mononuclear cell
PC: principal component
SNV: single nucleotide variant
WGS: whole genome sequencing

## SUPPLEMENTAL METHODS

### Subject recruitment

Subjects were recruited for participation in our cardiomyopathy biobank. Patients undergoing cardiac procedures, as well as non-cardiac patients with known genetic mutations (as identified by their health care provider) were targeted. In the latter case, we had 4 DCM subjects (two of which exhibit reduced ejection fraction <45%), 2 HCM subjects (with IVSd >=1.8 cm but wall thickness 0.9 and 1.2 cm), and 4 LVNC subjects who lacked a clinical diagnosis. These samples were used for WGS but not the cardiomyocyte differentiation and subsequent RNA-seq with the exception of subject 969 (DCM, reduced LVEF of 23.8%), subject 544 (HCM, IVSd 1.8cm), subject 603 (HCM, IVSd 2.1cm). Their data is included in the WGS figures except where indicated.

Healthy subjects without known genetic mutations and lacking a progressive condition were recruited from our cardiovascular prevention clinic. An additional category of control patients (referred to as “other” in Table S1) represent patients with non-cardiac conditions who were recruited at the clinic and over the phone, with permission of their providers. Two patients with known cardiac conditions other than cardiomyopathy (long QT syndrome and Fabry disease) were also recruited. Echocardiogram assessment of left ventricular ejection fraction (LVEF) and interventricular septum thickness, end diastole (IVSd) from the most recent measurement in the electronic medical record were queried and populated in RedCap when available.

### iPSC reprogramming

Induced pluripotent stem cells were reprogrammed from PBMCs using Sendai virus (CytoTune iPS 2.0 Sendai Reprogramming Kit) as previously described.[39] Three clones were generated per subject, karyotyped (KaryoStat, ThermoFisher Scientific), determined to be mycoplasma-free, and evaluated by immunohistochemistry for expression of pluripotency markers TRA-1-60 (LifeTech MA1023) and SSEA4 (LifeTech MA1021). Cells were maintained under feed-free conditions in mTeSR (STEMCELL Technologies, 5850) or Essential 8 media (Fisher, A1517001) and stored in liquid nitrogen.

To assess pluripotency of our cohort, we compared our RNA-seq data from 102 iPSC lines to 196 iPSC lines from the HipSci project (human induced pluripotent stem cell initiative) of the Wellcome Sanger Institute and EMBL (Expression Atlas ID for dataset: E-MTAB-4748)[40]. The HipSci dataset also contained 5 fibroblast samples and 4 PBMC samples for control. Expression from the HipSci project was publicly available as an expression matrix with expression tabulated as transcripts per million (TPM). To enable equal comparison, we used our raw RNA-seq data to tabulate TPM for our cohort (tabulated using DESeq2). (Note that we used salmon-aligned [ensemble90] RNA-seq data versus STAR, as this initial quality control assessment of the biobank was done prior to designing our subsequent RNA analysis workflow.) The joint HipSci-Stanford TPM dataset was log2 transformed. Stanford iPSC lines were all derived from blood while HipSci lines were derived from either blood or skin tissue, both of which are from the mesoderm lineage. We selected both pluripotency and mesoderm genes for examination based on the iPSCORE resource (genes taken from Figure 2A of the iPSCORE paper).[41] The pheatmap package in R was used to generate a heatmap (samples and genes clustered using Euclidean distance). We confirmed our iPSC cohort exhibited similar expression profiles as the HipSci iPSCs and did not cluster with PBMC samples (Figure S1).

### Cardiomyocyte differentiation and drug treatment

As previously described,[42] iPSCs were plated on Matrigel and cultured in StemMACS iPS-Brew XF (MACS Miltenyi Biotec, 130-104-368) until the final passage in Essential 8 media (Fisher, A1517001). Cardiomyocyte differentiation was induced at 60-80% confluency, with culture in RPMI media (Gibco/LifeTech 11875-119) plus B27 supplement lacking insulin (Gibco/LifeTech A1895601). 6µM of CHIR-99021 (Fisher, NC0976209) was added on day 0 and 6 µM IWR1 (Fisher, NC1319406) was added on day 3. Beginning on day 7, media was changed every other day using RPMI media supplemented with B27 containing insulin (Gibco/LifeTech 17504-044). Upon commencement of beating (around day 15), cells underwent purification via a three-day glucose starvation (RPMI media without glucose [Gibco/LifeTech 11879-020] supplemented with insulin-containing B27), a one-day recovery in glucose-containing media, and subsequent replating (dissociated in TrypLE, Fisher, 50-591-353). Cells were then maintained in RPMI media supplemented with insulin-containing B27 until approximately day 30. After differentiation, drug treatment occurred at 0 hours and 24 hours and samples assayed at 48 hours. Cells were treated with 250nM MYK-461 (Cayman Chemical, 19216-5mg), 400nM or 1uM omecamtiv mecarbil (Selleckchem via Fisher, NC1069600), or DMSO.

Additionally, at approximately day 24, one to three wells of the ongoing differentiation were replated (dissociated with TrypLE) into 96-well plates for immunohistochemistry (two wells, ∼40,000 cells/well) or 384-well plates (Thermo, 142761, ∼20,000 cells / well) for contractility assays and maintained in parallel until the end of differentiation. Cardiomyocytes were analyzed by immunohistochemistry to assess purity as previously described,[42] via staining for cardiac troponin T (Rabbit cTnT, Abcam, ab45932, 1:100). Cells were imaged on the Cytation5 Image Reader (BioTek) running the accompanying software (Gen5 Image+ version 3.03) to screen differentiations for a minimum of 90% cTnT positive cells.

### Whole genome sequencing

Library preparation and sequencing was performed by Macrogene (first 10 samples) and Novogene on genomic DNA we extracted from iPSC cells (Qiagen DNeasy kit). Paired-end 150bp reads were acquired on the Illumina HiSeq X Ten for a minimum of 90 gigabases of data. Reads were processed using Sentieon’s FASTQ to VCF pipeline (Sentieon version 201808.07).[43] This pipeline is a drop-in replacement for a BWA[44] plus GATK best-practices[45] pipeline for germline SNVs and indels, but has been highly tuned for optimal computational efficiency. BWA alignment to hg38 was followed by deduplication, realignment, base quality score recalibration, and variant calling to generate g.vcf files for each sample. Coverage was assessed (GATK version 3.7) (Tables S2 and S3). Individual sample g.vcf files were joined and variant quality score recalibration performed.

### Curation of candidate pathogenic mutations

To manually curate pathogenic and likely pathogenic variants we first created an overly-broad list of potential cardiomyopathy genes (referred to as our “panel genes” in the figures) (Table S4). The rationale was to include genes posited to play a role in cardiomyopathy, even where the data supporting a causal role was sparse to create a more comprehensive list of candidate mutations that we then filtered further. This included genes from six clinical genetic testing panels for HCM and DCM, the American College of Medical Genetics (ACMG) recommended list of genes to test for in HCM or DCM,[46] any gene annotated for HCM, DCM, or LVNC in the Human Genome Mutation Database, and genes evaluated for HCM or DCM pathogenicity in two systematic studies from the literature.[5, 6] We used ANNOVAR[47] to apply various filters, generating different pools of mutations (Figure S2) for manual interpretation.

Others have suggested a maximum minor allele frequency of 1 × 10^−4^ for cardiomyopathy.[5] For pool 1, we set a more inclusive filter for a minor allele frequency less than 0.01, which is the threshold for a rare variant, (frequency in ExAc, version November 2015), and required the variant be an exonic (excluding synonymous SNVs) or splicing mutation or have a CADD phred score greater than or equal to 20. Thus, pool 1 represents rare variants with the potential to alter protein sequence in our “panel genes”. For the sake of thoroughness, we also sought to capture mutations regardless of their likeliness to alter protein sequence if they were rare enough. These were curated separately in pool 0. For pool 0, we filtered for variants with a minor allele frequency less than or equal to 0.001 in ExAC or 1000 Genomes (version August 2015). Pool 0 (15.9 million mutations) and pool 1 (6082 mutations) were too large to examine manually. We thus further filtered for a ClinVar designation of pathogenic or likely pathogenic (for any disease) as curated by ANNOVAR (and thus a reflection of the latest ClinVar information in the ANNOVAR database). We found a large number of rare GATA4 variants in introns (933 mutations) or untranslated region (270 mutations) that had been flagged in ClinVar for congenital heart disease (and not cardiomyopathy). After removing these for lack of relevance to HCM and DCM, we had 159 mutations in pool 5. We call pool 5 “WGS_P” for pathogenic, to demarcate this filtering strategy was dependent on a pathogenic or likely pathogenic ClinVar designation. These represent our first strategy for filtering for candidate variants. We evaluated each of these manually and with CardioClassifer, an online research tool for annotating pathogenicity of cardiomyopathy mutations.[48] However, we then went back and applied additional filtering strategies to overcome some of the technical limitations of this strategy. Below is a brief description. See Figure S2 for the full filtering workflow.

The first complication we addressed was that our variant calling workflow had the potential for a larger indel to be miscategorized as two neighboring smaller indels or SNVs. We thus created pool 10 to merge nearby mutations and evaluate the resulting larger mutation for pathogenicity. This step was performed only for the diseased samples and not the control subjects. We started by flagging any mutation within 40 bp of another mutation in the same subject (365 mutations). We removed individual indels greater than 50 bp since this could have represented a sequencing error. (This was applied before merging neighboring mutations). For SNVs, we merged SNVs if they occurred within 2 bp of each other (ie could be on the same codon, and thus their expected effect on protein sequence would only be properly determined when analyzed together). We also merged SNVs within 5 bp of an indel to expand the indel. We then confirmed that the neighboring mutations had the same zygosity and were on the same allele, thus justifying our analysis of them in tandem. We call pool 10 “WGS_merge’ to indicate it represents manually merging of nearby mutations.

The second complication we addressed is that our first filtering strategy was dependent on ClinVar flags. This could lead to many false negatives due to many variants not being listed in ClinVar. We thus took any of the pool 1 variants (rare variants with the potential to alter protein sequence of “panel genes”) that hadn’t had a ClinVar flag and kept them in the analysis if they met the more stringent allele frequency of less than 0.001 (pool 13). Note that for Pool 13, unlike the previous frequency filters, here we used the maximum frequency in 1000 Genomes and any individual ethnic group in ExAc (to screen out mutations that while rare in genomic datasets as a whole, are more abundant in specific ancestral backgrounds). We needed to further curate pool 13 to a list that was feasible for manual evaluation. We applied two separate additional filters. First, we kept any mutation in pool 13 that was in a gene for which the CardioClassifer tool could be applied, given that this overcame the technical limitation of manual curation and would allow us to first screen mutations via the tool. This created pool 14. (CardioClassifier is an expert-developed tool incorporating cardiomyopathy specific knowledge to apply ACMG guidelines.) The CardioClassifier genes for HCM are MYH7, TNNT2, TPM1, MYBPC3, PRKAG2, TNNI3, MYL3, MYL2, ACTC1, CSRP3, PLN, TNNC1, GLA, FHL1, LAMP2, and GAA. The CardioClassifier genes for DCM are LMNA, TNNT2, SCN5A, TTN, TCAP, MYH7, VCL, TPM1, TNNC1, RBM20, DSP, and BAG3. For LVNC we used the 12 DCM genes. The CardioClassifer genes for long QT syndrome were KCNQ1, KCNH2, SCN5A, and KCNE1. For pool 14, we required that the mutation fall in a CardioClassifer gene associated with the disease of the subject. We call pool 14 “WGS_freq” to indicate these are mutations that lacked a ClinVar flag but were kept in the analysis due to their low frequency.

Given that truncating variants can have an especially dramatic effect on protein sequence, we separately evaluated pool 13 for mutations that may change the length of the protein sequence to create pool 15. For pool 15, we included stop-gain, stop-loss, frameshift insertion, or frameshift deletion mutations. (Note that for stop-loss and frameshift insertions, they could act to increase protein sequence rather than truncate.) We removed indels greater than 50 bp due to the possibility they represent sequencing artifact. There were 95 mutations, but removing those already identified in pool 14 left 46. We call pool 15 “wgs_trunc” for truncation, to indicate they may alter protein length. For variants most likely to be pathogenic truncating variants (heterozygous, stop-gain mutations) we performed additional characterization, using the RNA-seq data from the iPSC-derived cardiomyocytes where available. First we used our combat-corrected processed data (see Supplemental Methods section for RNA-seq) to compare gene expression in the mutation-carrying line to the other cardiomyopathy (HCM or DCM depending on the disease of the mutation-carrying line) or control lines. Second, we re-processed the RNA-seq fastq files to get allelic expression via STAR, setting the waspOutputMode as SAMtag and inputting a vcf file for the line containing the mutation of interest.

Pools 14 and 15 generated candidates with less definitive annotation data. Thus as a control to provide confidence on the likeliness for false positives, we applied the same filters to the control subjects to evaluate the rate of detecting mutations with these filters in a cohort that should have few true pathogenic or likely pathogenic mutations (pool 16). We filtered for CardioClassifier’s “cardiomyopathy” gene list: ACTC1, BAG3, CSRP3, DSP, FHL1, GAA, GLA, KCNE1, KCNH2, KCNQ1, LAMP2, LMNA, MYBPC3, MYH7, MYL2, MYL3, PLN, PRKAG2, RBM20, SCN5A, TCAP, TNNC1, TNNI3, TNNT2, TPM1, TTN, VCL. We call pool 16 “WGS_healthyfreq” and pool 17 “WGS_healthyTrunc” to indicate it is the same filters from WGS_freq and WGS_trunc applied to the control subjects.

We also pulled any variant listed in the electronic medical record (EMR). For many of these we had already assessed pathogenicity as part of our WGS workflow. However, some variants in the EMR had not passed our WGS filters and had not been annotated yet. We collected these into pool 8 for evaluation. Often, pathogenicity classification for the variant was provided in the EMR, however we always classified them ourselves as well in case the original annotation pre-dated new information in the literature. We call pool 8 “Clin_research” to indicate they are variants that came from the clinical genetic testing for which we needed to research their potential pathogenicity.

Our “final pool” represents all the mutations from all of these filtering strategies. For any variant in our final pool that was only found in WGS data and not listed in the EMR (not clinically validated to be present in the subject’s genome), we further examined the mutation in the vcf file for quality metrics to confirm confidence that the mutation was present. The final pool became Table S5. Column K indicates which filtering strategy resulted in identification of the mutation. Note that if a mutation was identified from our first filtering strategy “WGS_P” it will be listed as such in Column K. Even if the variant is truncating or rare, it won’t be listed as “WGS_trunc” or “WGS_freq” because these additional filtering approaches were not necessary to identify the mutation. Thus column K represents the minimal filtering we needed to identify the variant.

### Comparison of mutation burden in cardiomyopathy genes with echocardiogram measurements

We first analyzed the distribution of pool 1 variants (Figure S2) between HCM, DCM, and control lines. We found six samples (control lines 820, 822; HCM lines 543, 598; DCM line 596, 969) were outliers (z score > 3) for having a large number of pool 1 variants. Thus the subsequent analysis of mutation types in the control, HCM, and DCM cohort were done on the full cohorts and after removing these six samples to ensure there were no differences in the results. Starting with the pool 1 variants, we removed mutations with frequency > 0.001 in 1000 genomes or any ExAc ethnicity. (Previous ANNOVAR filter used to generate pool 1 used the mutation frequency in ExAc as a whole, while here we used the maximum frequency in any ethnicity.) We removed indels > 50bp as these could be due to a sequencing error. We removed mutations shared by more than 10 patients. (Only 3 mutations fit this description. They were shared by 171, 46, 31 patients). The next most common mutations were shared by 7 patients. This is also the max frequency for a mutation we annotated as pathogenic or likely pathogenic. Finally, we grouped the mutations into two categories. The first category was mutations that could change protein length (frameshift insertion, frameshift deletion, stop-gain, stop-loss). The second category was all other exonic or splicing mutations. For calculating mean and standard deviation values, the two “other” samples with known cardiac conditions (long QT syndrome and Fabry’s disease) were excluded from the control cohort. P-values for figure 2A are calculated using t-test.

Pucklewartz et al.[16] defined a set of 102 cardiomyopathy genes whose cumulative burden of nonsynonymous SNVs correlates with LVEF in DCM. We replicated this analysis by summing the instances of a nonsynonmous SNV in the 102 genes. This was done by going back to the original Annovar output files for SNVs (we did not include indels) and using R to identify all nonsynonymous SNVs regardless of allele frequency (as opposed to starting from our pooled of filtered rare variants). We set a cutoff of DP (depth of coverage) >=8 and GQ (genotype quality) >=20. Zygosity was not incorporated. The total burden was plotted against LVEF and linear regression computed. We did this for both the HCM and DCM lines. We the repeated the analysis for DCM after setting an additional threshold of maximum allele frequency of 0.5 (using the maximum frequency in 1000 Genomes and any individual ethnic group in ExAc). This was done separately on DCM samples with a known pathogenic or likely pathogenic variant (P/LP) and those without (nopatho). Finally, we applied a further filter for the variants, restricting variants to 20 core DCM genes with greater evidence for pathogenicity (as defined by appearing in at least one of the following: [4] or DCM genes only[5, 15]). The core genes are: ACTC1, ACTN2, BAG3, DES, DSP, FLNC, JPH2, LMNA, MYH7, NEXN, PLN, RBM20, SCN5A, TCAP, TNNC1, TNNI3, TNNT2, TPM1, TTN, VCL.

To assess mutational burden in HCM samples within the promoter regions of the 102 Puckelwartz et al genes, we defined a promoter as 2000 bp upstream and 500 bp downstream of the transcription start site and collected all SNV variants (not indels) regardless of frequency and regardless of mutation type. DP (depth of coverage) >=8 and GQ (genotype quality) >=20 filters were applied. Unlike the analysis of LVEF versus coding variants in DCM, for promoter analysis we did not restrict the variants to nonsynonymous SNVs. For each gene we computed the mean number of variants in the control and HCM cohorts separately. 54 genes had higher mean in control than promoter. These were: A2ML1, ALPK3, BAG3, CACNA1C, CALR3, CASQ2, CAV3, CHRM2, CSRP3, CTNNA3, DES, DOLK, EMD, EYA4, FHL1, FKTN, GATA6, GATAD1, JUP, KRAS, LAMP2, LDB3, LMNA, LRRC10, MAP2K1, MYL2, MYOM1, MYOZ2, MYPN, NEBL, NEXN, NKX2.5, NRAS, PDLIM3, PRDM16, PRKAG2, PTPN11, RAF1, RASA1, RBM20, RRAS, SCN5A, SHOC2, SLC22A5, TAFAZZIN, TCAP, TGFB3, TMEM43, TNNC1, TNNI3, TNNT2, TPM1, TRDN, TXNRD2. We performed regression on the total mutation count in these promoters compared to IVSd for HCM samples with and without known pathogenic mutations and accounting for RNA subgroup (steep or moderate). Finally, we applied the published polygenic risk score[29] to the HCM samples. The dbSNP IDs were used to convert from hg37 to hg38 coordinates and search the Annovar output files for overlapping variants. In instances where a variant was not returned for the loci, we assumed the sample had the reference allele. For each variant we determined presence or absence of the risk allele (ignoring zygosity) and multiplied by the published beta values, summing across all variants to get the final risk score. The score is composed of 36 SNVs.

### RNA-seq library preparation, sequencing, quality control, and expression matrix generation

RNA was extracted from iPSCs or cardiomyocytes (RNeasy, Qiagen). Illumina RNA-seq libraries (TruSeq Stranded Total RNA LP Gold) were prepared on the Bravo (Agilent; 3 samples prepared manually as indicated in Table S6), pooled (Table S6), and sequenced (NovaSeq-6000, paired-end, 100bp). Where possible drug treatment conditions for the same differentiation were kept together in batches, while replicate differentiations for the same iPSC lines were split apart, and HCM, DCM, and control samples were distributed across batches. Reads were aligned to hg38 (STAR). Principal component analysis on cardiomyocyte and iPS samples separately returned no outlier samples (as defined as Zscore of principal component 1 > 3). Library quality control was assessed via fastp, fastQC, STAR, and Picard metrics. Samples were flagged for poor QC by the following metrics: GC content after filtering outside of 20-80% (fastp), duplication rate greater than 40% (fastp), uniquely mapped read pairs (fragments) < 20 million (STAR), mean reads (average of forward and reverse) <20 million (fastQC), ribosomal RNA bases > 20% (Picard), coding plus UTR (untranslated region) < 50% (Picard), uniquely mapping fragments <60% (STAR). Samples with more than one flag were removed. Cardiomyocyte and iPSC samples were subsequently processed separately. Reads were computed as CPM (edgeR) and corrected for library preparation batch (combat-seq) and TMM normalized (edgeR) to generate the final expression matrix. For samples with biological replicates, TMM counts were averaged. Principal component analysis was performed and principal component 1 assessed for spearman correlation with the following metadata: percent GC content (fastp), mean reads (average of forward and reverse) in millions (fastQC), percent ribosomal RNA bases (Picard), uniquely mapped fragments in millions (STAR), duplication rate (fastp), percent coding or UTR (picard), library preparation batch, and sequencing pool. The maximum absolute value for spearman correlation between PC1 and the library metadata was 0.08 for cardiomyocyte samples, indicating good quality control with technical artifacts having minimal influence on the dataset. iPSC samples had higher correlation for three metrics (0.26 with GC content, 0.22 with duplication rate, and 0.11 with percent coding or UTR), with the remaining less than an absolute value of 0.04.

### DESeq2 analysis of differential expression

Raw data was input into DESeq2 (as required for DESeq2) with library preparation batch included in the design (in line with the combat-seq correction strategy we used for generating our final expression matrix). We assessed baseline (without drug) control vs HCM cardiomyocytes and control vs DCM cardiomyocytes separately and determined significance (Benjamin-Hochberg corrected p-value < 0.05). Drug treatment was compared to DMSO using all samples regardless of disease. Geno ontology analysis for differentially expressed genes (or for *ADCY5* connected genes in the network, see below) was performed using DAVID bioinformatics,[49] with enriched ontologies defined as Benjamin-Hochberg corrected p-value < 0.05.

### Personalized co-expression network construction using lionessR

Linear interpolation to obtain network estimates for single samples was performed using lioness[18, 19] implemented in R (lionessR package). This was done separately on the HCM versus control cohort and the DCM versus control cohort. First, a cohort-level network was built using the control and diseased samples. The finalized cardiomyocte expression matrix (TMM normalized, batch-corrected) was input. The dataset was refined to the top 2000 most variable genes (greatest standard deviation between all samples, diseased and control samples combined). For the control and diseased samples separately a co-expression matrix was computed using Pearson correlation for each gene-by-gene comparison. The control matrix was subtracted from the diseased matrix to assess differential co-expression between the control and diseased cohorts, and the network was trimmed to the 200 most differential edges (LIMMA) between control and disease. Doing this for both the HCM and DCM data, we thus built two networks: an HCM network (reflecting differential co-expression between HCM and control samples) and a DCM network (reflecting differential co-expression between DCM and control samples). Personalized co-expression networks were inferred for each sample individually through an iterative process where lionessR removed one sample from the cohort, recalculated the cohort edge strengths, and determined the difference in cohort edge strength with and without the sample, and then applyied a linear model to estimate the edge weights of the sample.

### Node strength calculation

Node strength represented the total weight of all edges surrounding a gene. We calculated this in two ways. For Figure 3B, this was calculated by summing the weights of all edges surrounding a gene. This was displayed by plotting the summed weight on the x-axis for different genes along the y-axis. Samples were colored from light shades (small ADCY5 node strength) to dark shades (large ADCY5 node strength) and maintained the same color when displaying nodes strengths of other genes. This was useful for visualizing the variability across our diseased cohorts. For subsequent analyses of node strength in Figures 4 and 5, we modified the calculation such that greater node strength would indicate greater difference from non-diseased samples. Each edge surrounding a gene were previously determined by lionessR to be red (stonger in disease) or green (stronger in control). This was colored based on behavior of the full cohort. To calculate node strength in each sample, we subtracted the sum of the green edges from the sum of the red edges.

### Assessment of co-modulation of edges around a common hubnode

We defined hubnodes as genes that were connected to at least three other genes. We tested which hubnodes represented units of network activation, in that higher co-expression of one of the edges co-occurred with higher co-expression of the other edges. For each disease network, we analyzed the disease (HCM or DCM) and control cohorts separately. We first calculated the Pearson correlation coefficient for each edge-by-edge comparison. Second, we subset all edge-edge pairs surrounding a shared hubnode, called the “All” dataset. We also created a “background” dataset with all edge-edge pairs expect those for which the same gene was shared in both edges. We randomly sampled the All and Background datasets and calculated the difference in Pearson correlation coefficient (All – Background). We did this 10,000 times to obtain the mean and 95% confidence intervals. Nodes whose 95% confidence interval bars do not cross zero are concluded to exhibit co-modulation of the edge strengths for their surrounding edges.

### Principal component analysis

The edge weights for the HCM cohort, control cohort, and individual HCM samples were analyzed by principal component analysis (prcomp, scale=TRUE, in R). Separately, the same was done for the DCM cohort, control cohort, and individual DCM samples. Linear regression compared principal component 1 (PC1) to echocardiogram measurements (LVEF for DCM, IVSd for HCM) (Figures 4B and 4D). The contribution of *ADCY5* edges to PC1 for the HCM network (Figure 5A) was computed. For each sample, the scaled edge weights were multiplied by the loadings for PC1 and all *ADCY5* edges summed.

The relative contribution of each edge to the principal component was calculated as: (loadings^2) / sum(loadings^2). This was done for the full network (not on individual samples). Edges were ordered from highest to lowest relative contribution. Visual inspection of the list revealed enrichment of *ADCY5* edges at the top of the PC1 list for highest contribution and *ANLN* edges at the top of the PC3 list. This was confirmed by plotting the cumulative contribution (sum of relative contributions) with each successive edge added (Figure 5B) alongside the cumulative contribution specifically from *ADCY5* or *ANLN* edges only. (Note that for clarity, in Figure 5B, the cumulative contribution for either ADCY5 or ANLN was not plotted as a smooth curve, but only displayed at the *ADCY5* or *ANLN* edges.)

Pearson correlation was used to obtain R^2^ values for correlation of hubnode strength to PC1 and to echocardiogram measurements and as well as to compare *ADCY5* node strength to *ADCY5* expression and *MEF2A* expression in HCM samples.

### Kinetic imaging cytometry to measure contractility

Contractility was assessed as previously described.[50, 51] On approximately day 24, cells were dissociated (TrypLE, Fisher, 50-591-353) and replated in Matrigel-coated 384 well plates (20,000 cells/well, 8 wells per drug condition), and maintained in parallel for the remainder of the differentiation. The four perimeter rows and columns of wells were not used, and filled with PBS to minimize the effect of temperature fluctuation on the assay. At the time of assay, 400nM tetramethylrhodamine, methyl ester (TMRM, Marker Gene Techonologies), a live cell stain of mitochondria, was added to the cardiomyocyte cell culture, and the cells were returned to the 37°C incubator for approximately 15 min to restabilize their temperature. Contractility was analyzed on the IC200 Kinetic Imaging Cytometer (Vala Sciences) running CyteSeer software (Vala Sciences), using a 0.75 NA 20x Nikon Apo VC objective. Cells were maintained at 37C°, 5% CO_2_ throughout the assay. Time series images were acquired over 10 second at 33 Hz frequency. A custom MatLab script was used to further process the outputs of CyteSeer and extract key metrics of contractility, including averaging multiple contractions per well into a representative peak and extracting the area under the curve (AUC) as well as average time between peaks (T.peak). [52] AUC divided by T.peak represented the total amount of deformation normalized to time. Each differentiation was assayed in 8 wells per condition (DMSO and drugs).

### *ADCY5* hub node correction with drug treatment

For each edge surrounding *ADCY5* in the HCM and DCM networks respectively, we calculated the mean edge weight in disease at baseline and after drug treatment as well as in control samples at baseline. We computed the difference as such: HCM_DMSO_ – Control_DMSO_; HCM_MYK_ – Control_DMSO_; DCM_DMSO_ – Control_DMSO_; DCM_OMEC_ – Control_DMSO_. We then converted the differences to absolute value. Box and whisker plots displayed these values for all edges in each comparison.

