## supplement for "Personalized transcriptome signatures in a cardiomyopathy stem cell biobank"

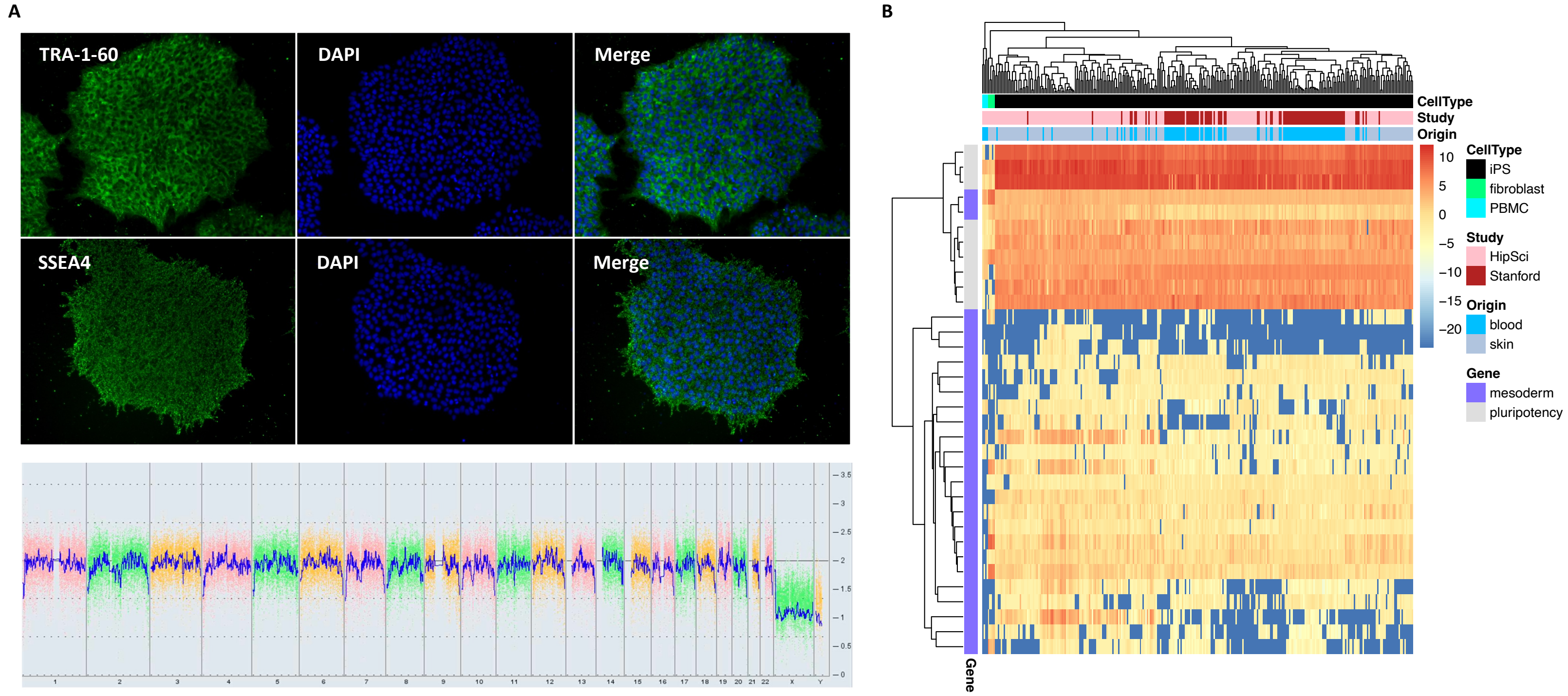

**Supplemental Figure 1. iPS cells were successfully reprogrammed. A.** Representative quality control images. All iPS lines were confirmed to express pluripotency markers TRA-1-60 and SSEA4 and show normal karyotype. Staining (10X magnification) and karyotype report shown for male line 941. **B.** Expression from 196 iPS lines from the HipSci project of the Wellcome Sanger Institute and EMBL was compared to the expression of 102 iPS lines from our biobank. Key pluripotency genes as well as mesoderm genes were selected for examination based on the iPSCORE resource. Stanford biobank lines express pluripotency genes, cluster away from PBMC and fibroblast samples, and cluster with the benchmark HipSci samples.

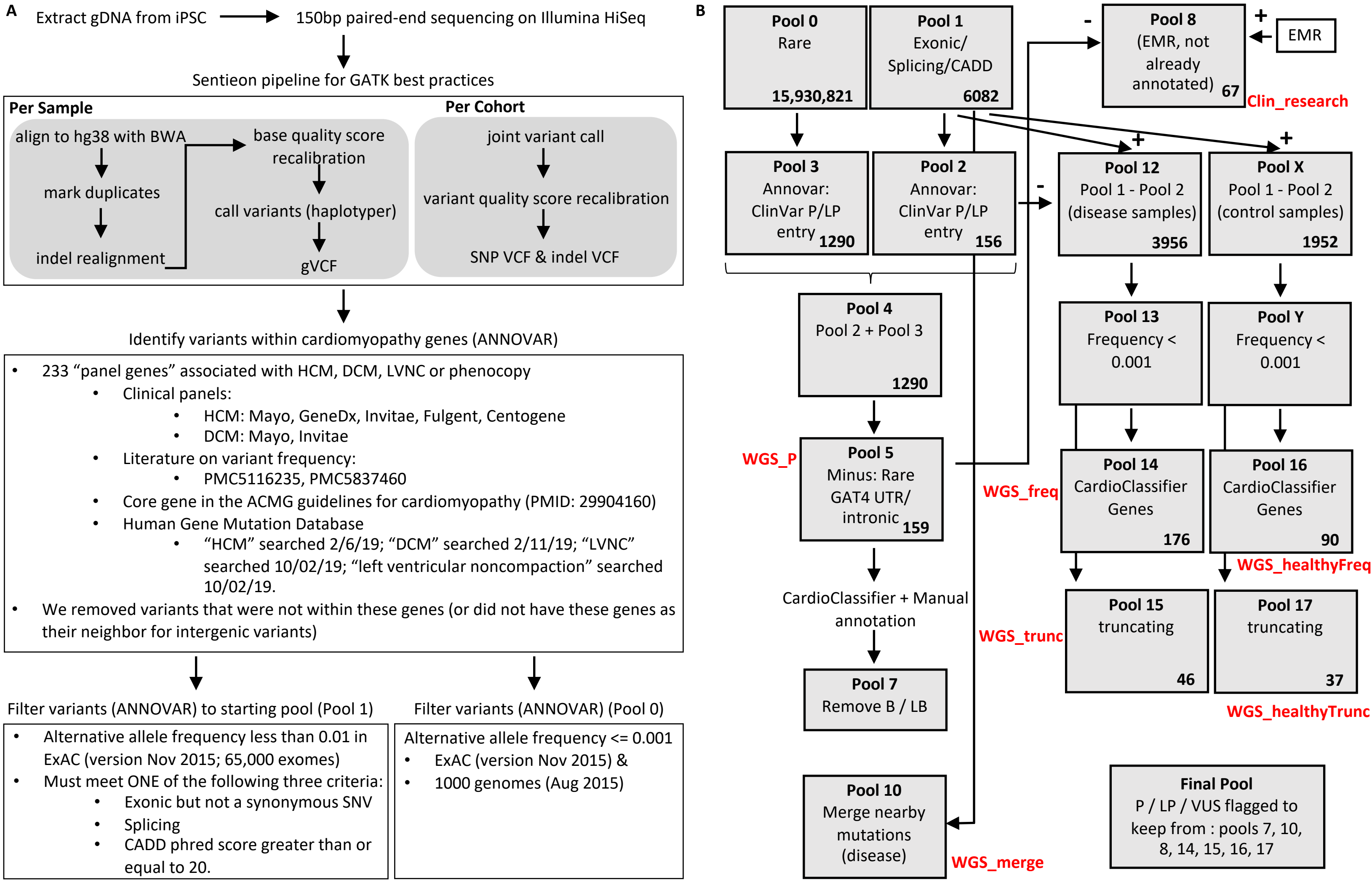

**C** **WGS_P**: variant in pool 5.
**WGS_VUS**: variant in pool 1 but not pool 5
**WGS_NotInPanel**: gene not one of the "panel genes" we curated
**WGS_class**: for variants we did not detect because they are larger than a SNP or Indel
**WGS_vqsr**: variant present in earlier cohort vcf (when we had only a subset of the WGS data) but lost due to variant quality score recalibration on larger cohort
**WGS_merge**: For insertion deletions that were broken up as a byproduct of our analysis pipeline. We had the correct sequence identified in pool 1, but it was split across multiple "mutations" and didn't make it to pool 5. Instead we found it in pool 10 merge analysis.
**WGS_AbsentDoubleCheck**: This is for a variant that we didn't find in pool 1 but we double checked if it was in the data because it was in the EMR, so we went back and checked the vcf cohort file before vqsr to see if present in the sample, and still not there. So conclusively not in our WGS.
**WGS_freq**: variant found in pool 14
**WGS_trunc**: variant found in pool 15
**WGS_healthyFreq**: variant found in pool 16
**WGS_healthyTrunc**: variant found in pool 17
**Clin_P**: variant listed in EMR and annotated as P/LP in EMR
**Clin_conflict**: variant listed in EMR and annotated as conflicting interpretations in EMR
**Clin_VUS**: variant listed in EMR and annotated as VUS in EMR
**Clin_NotInPanel**: patient underwent clinical testing but variant not listed in EMR because gene (or splice/intronic region of gene) not in the panel
**Clin_Absent**: patient underwent clinical testing but variant not listed in EMR despite gene being included in the panel. In some cases, the EMR doesn't contain VUSs. So it's possible they found the VUS but it didn't make it into the EMR (or at least the subset of the EMR data that the research team received)
**Clin_fam**: variant listed in EMR only associated with a family member and not confirmed in this patient
**Clin_research**: Didn't have ClinVar flag from Annovar (not in pool 5) nor was it flagged as P / LP in EMR. However subsequent analysis of EMR variants that weren't P/LP annotated pulled this out (pool 8)
*Clin_P and WGS_P do not mean our final annotation is P/LP.

**Supplemental Figure 2. Mutation filtering. A.** Mutations were processed as outlined. **B.** The ANNOVAR-annotated lists were further culled before manual annotation. The number of mutations in each pool is indicated in bold. The same mutation in multiple lines is counted multiple times. See Supplemental Methods for additional description. **C.** Filter shorthand used in Table S5 to demarcate how each candidate mutation was found. A subset of these are also labeled in red in the diagram (B). Abbreviations: pathogenic / likely pathogenic (P/LP), benign / likely benign (B/LB).

A

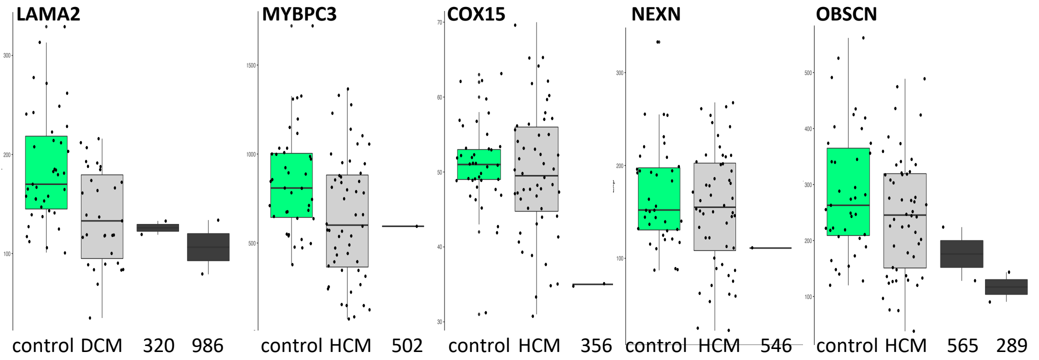

| Donor | Disease | Gene | RNA | REF count | ALT count | % ALT | Annotation | Filters |
|---|---|---|---|---|---|---|---|---|
| 320 | DCM | LAMA2 | 320dmso_r1 | 109 | 30 | 22% | LIKELEY PATHOGENIC | WGS_P |
| | | | 320dmso_r2 | 141 | 18 | 11% | | |
| 986 | DCM | LAMA2 | 986dmso_r1 | 280 | 6 | 2% | LIKELEY PATHOGENIC | WGS_P |
| | | | 986dmso_r2 | 473 | 56 | 11% | | |
| 502 | HCM | MYBPC3 | 502dmso_r1 | 1479 | 232 | 14% | PATHOGENIC | WGS_P |
| 356 | HCM | COX15 | 356dmso_r1 | 44 | 1 | 2% | UNCERTAIN SIGNIFICANCE | WGS_trunc |
| | | | 356dmso_r2 | 28 | 7 | 20% | | |
| 546 | HCM | NEXN | 546dmso_r1 | 144 | 8 | 5% | UNCERTAIN SIGNIFICANCE | WGS_trunc |
| 565 | HCM | OBSCN | 565dmso_r1 | 65 | 32 | 33% | UNCERTAIN SIGNIFICANCE | WGS_trunc |
| | | | 565dmso_r2 | 36 | 7 | 16% | | |
| 289 | HCM | OBSCN | 289dmso_r1 | Zero reads. Intronic variant | | | UNCERTAIN SIGNIFICANCE | WGS_merge |
| | | | 289dmso_r2 | | | | | |

B

**145 donors with clinical genetic testing**

| | Evaluated | P/LP Identified | | |
|---|---|---|---|---|
| | Clin & WGS | Clin & WGS | Clin Only | WGS Only |
| DCM | 53 | 28 | 2 (FBN1 [Marfan], tRNA Leu UUR) | 5 (FLNC, FLNC, LAMA2, MYBPC3, OBSL1) |
| LVNC | 10 | 1 | | |
| HCM | 82 | 29 | | 3 (MYBPC3 intronic, DSP, ALPK3) |

**2 donors with clinical genetic testing of a family member**

| | Evaluated | P/LP Identified | |
|---|---|---|---|
| | Clin on family & WGS | Clin on family & WGS | Clin Only |
| HCM | 2 | 1 | 1 misidentified (WGS revealed mutation NOT in patient, only in family member) |

**Supplemental Figure 3. WGS annotation is aided by RNA-seq, but still provides minimal improvement over clinical genetic testing.** **A.** RNA-seq from differentiated cardiomyocytes provided additional support to mutation annotation. Potentially truncating mutations (heterozygous, stopgain mutations) were evaluated for differential alleleic expression. Top boxplots show the overall (not accounting for allele) expression using our combat-corrected RNA-seq data. Bottom table shows read counts at the mutant loci exported from bam files utilizing wasp output feature in STAR to capture allelic information. The first three line-gene pairs (2 LAMA2 mutations and MYBPC3) show mutations we already annotated as P / LP before taking into account RNA data (see supplemental method for filters corresponding to WGS_P). The remaining mutations are those which only made it past our filters due to their reduced expression of the alternative allele in the presence of detectable reference allele expression (table) and their overall reduced expression in relation to the cohort (box plots). Even with this observation, there was not enough data to call these mutations as P / LP per ACMG guidelines, so they do not appear as definitive, for example in Fig 2A chart of P / LP mutations. Note that the OBSCN mutation in 565 was not deemed worthy of passing filters (see Table S5 for annotation notes), but was kept in as a comparison for OBSCN mutation in 289 which did show up as VUS to keep. (Note r1 and r2 in table refer to replicate differentiation experiments.) **B.** 145 cardiomyopathy patients had clinical genetic testing results in their EMR to compare to the WGS (left table). Indicated are the number of lines where we found a P/LP mutation via both methods, or only via one of the methods. Mutations found only in the clinical test were due to the omission of the gene from our "panel genes" used to annotate mutations. See supplemental table 5 for full explanation of why mutations were found or not.

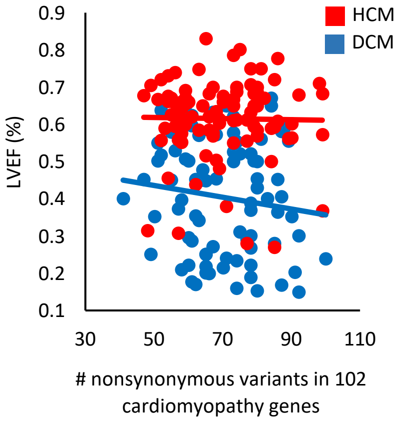

**Supplemental Figure 4. Comparison of the number of nonsynonymous variants in the Puckelwartz et al genes with left ventricular ejection fraction when combining P/LP and nopatho donors.**

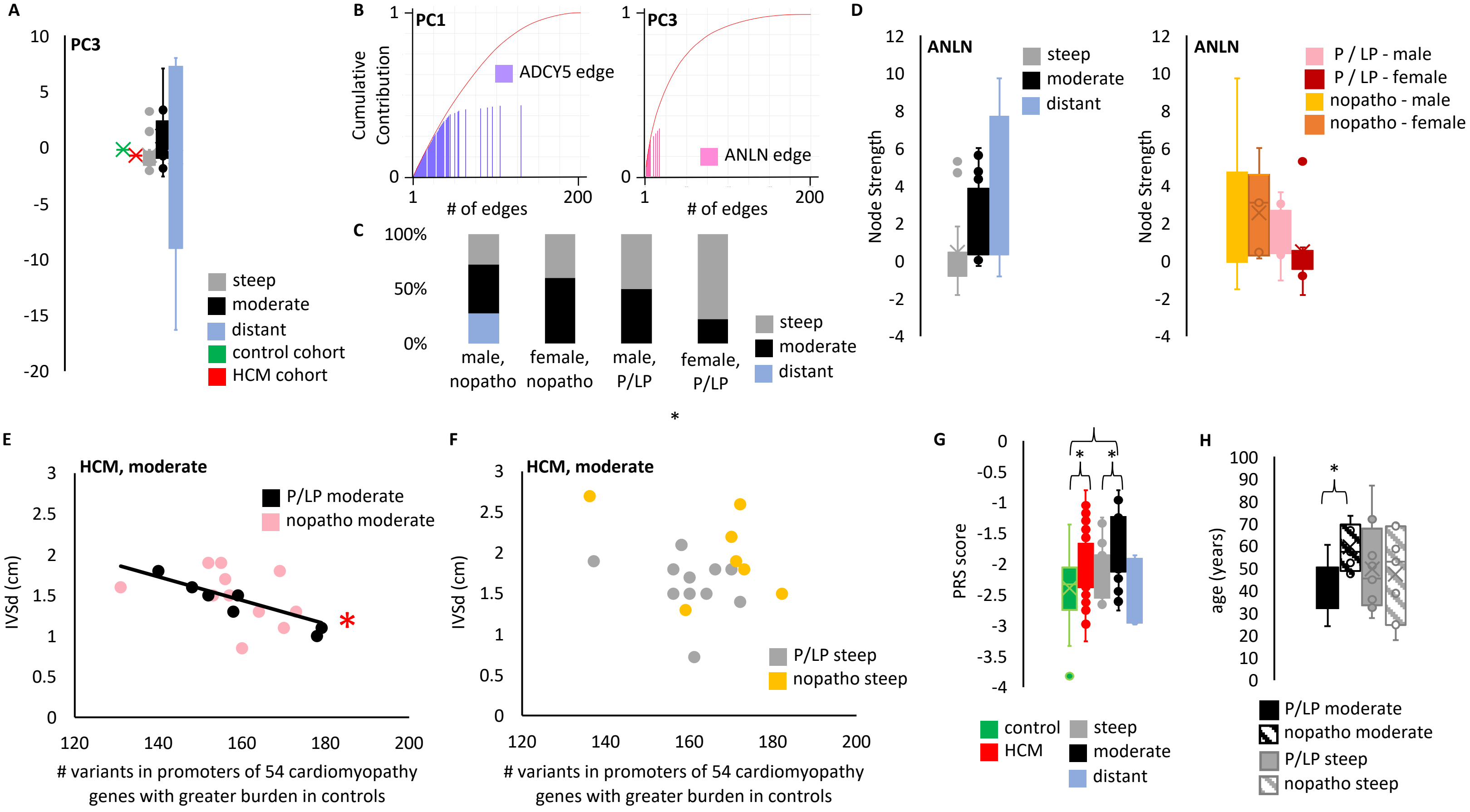

**Supplemental Figure 5. Features of demogrpahics, the transcriptional network, and genetic background distinguish HCM-moderate and HCM-steep samples. A.** PCA analysis of the co-expression correlations of the HCM network was performed as outlined in Figure 4B. We tested for principal components distinguishing steep and moderate HCM samples. Principal component 3 showed the greatest difference between steep and moderate samples. **B.** Plotted is the cumulative contribution to the principal component with each subsequent edge, ordered by edges with the greatest relative contribution first. (Left) *ADCY5* edges are highlighted in purple for their contributions to PC1. (Right) *ANLN* edges are highlighted in pink for their contribution to PC3. **C.** Demographic distribution of HCM samples between the three RNA groups. **D.** *ANLN* node strength (sum of all surrounding edge strengths) for HCM lines. **E.** Plotting the mutation burden in the same 54 promoters for all of the moderate samples, reveals a significant relationship between burden and IVSd (p-value = 0.0117), **F.** but not for steep samples. **G.** A published polygenic risk score for HCM was applied. Moderate samples on average show the expected increase in score while steep samples do not. (t-test: Control vs Moderate-HCM p-value = 0.00000257452. Control vs Steep-HCM p-value = 0.137757459. Control vs Distant-HCM p-value = 0.685843043. Control vs HCM-all p-value = 0.000000338723. Moderate-HCM vs Steep-HCM p-value = 0.011904577). **H.** Moderate HCM lines with a known pathogenic or likely pathogenic mutation come from younger donors (p-value = 0.00277).

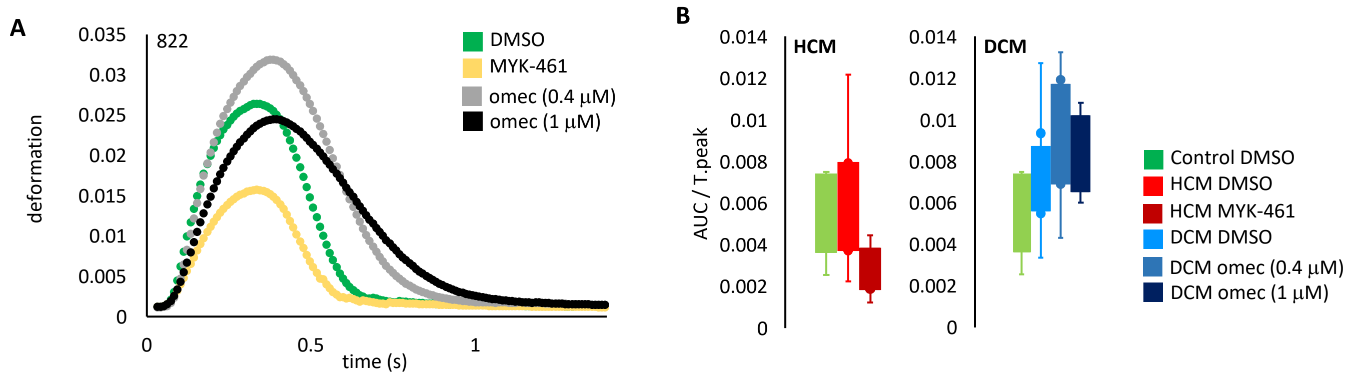

**Supplemental Figure 6. Treatment with mavacamten or omectamtiv mecarbil alters contractility. A.** Cardiomyocytes were treated with the small molecule sarcomere activator (omecamtiv mecarbil) or inhibitor (mavacamten) for 48 hours. Kinetic Image Cytometer microscopy was used to visually measure cellular deformation over time. Presented is a representative image from a control donor sample (line 822). Mavacamten reduces contractility, while omecamtiv mecarbil increases contractility. **B.** AUC (the area under the curve) and T.peak (the time between peaks) were extracted from the curves. AUC/T.peak represents the total deformation normalized to the length of a beat. Mavacamten treatment blunts and omecamtiv mecarbil treatment increases AUC/T.peak.

**Supplemental Table Legends:**

**Supplemental Table 1. Sample level demographics.**

**Supplemental Table 2. Whole genome sequencing passes quality control.** Cohort-level summary of whole genome sequencing showed appropriate values for key metrics. One control sample (line 2002) had low coverage below 20X.

**Supplemental Table 3. Sample level whole genome sequencing quality control metrics.**

**Supplemental Table 4. "Panel Genes" analyzed for potentially harboring cardiomyopathy mutations.** Column A – Gene list includes isoform entries and gene synonyms to fully capture the entries from the respective sources. Column B – ACMG guidelines list comes from Table 1 in (PMID: 29904160). ACMG identified DCM-specific genes as well as recommended DCM patients be tested for HCM and ARVC genes (indicated with "test" in annotation). Likewise, ACMG recommended testing LVNC patients for the associated cardiomyopathy, thus we used the DCM gene list (indicated with "test" in the annotation). Column C – LVNC genes from HGMD. Searched "LVNC" and "left ventricular noncompaction" on 10/2/19. Columns D-K: HCM genes. Column D – Designation in PMC5837460. Column E – HCM designation in PMC5116235. Column F – HCM genes from HGMD. Searched "HCM" on 2/6/19. Column G – Fulgent hypertrophic cardiomyopathy panel. Column H – Centogene hypertrophic cardiomyopathy panel. Column I – Mayo Clinic Inherited Disease Panel, hypertrophic cardiomyopathy designation. Column J – GeneDx hypertrophic cardiomyopathy panel. Column K – Invitae hypertrophic cardiomyopathy panel. Columns L-O: DCM genes. Column L - DCM designation in PMC5116235. Column M – DCM genes from HGMD. Searched "DCM" on 2/11/19. Column N – Invitae Cardiomyopathy Comprehensive Panel. Column O – Mayo Clinic Inherited Disease Panel, dilated cardiomyopathy designation. Abbreviations: American College of Medical Genetics (ACMG), Human Gene Mutation Database (HGMD).

**Supplemental Table 5. Candidate cardiomyopathy mutations.** Pathogenic or likely pathogenic identified in our cohort. Assigning pathogenicity is an imperfect process and these designations should be periodically re-evaluated. Column A – Mutation sequence. Column B – Gene name. Column C and D - ANNOVAR functional annotation. Column E – dbSNP ID where available (or for similar mutation at that loci as indicated). Column F – Donor ID. Column G – Disease of donor. Column H – Pathogenicity assignment. Column I – Notes on assignment where applicable. Notes use ACMG abbreviations for classifying variants. Column J – Genetic analysis platform used on the donor. All mutations list "wgs" to indicate we performed WGS on samples from this donor for this study. Some donors also list "clin" to indicate they received clinical genetic testing as denoted in their electronic medical record. Column K – Explanation for how the mutation was found (which of the clinical/WGS filters) or was not found (in cases where only found by clinical analysis or WGS but not both). (See Supplemental Figure 2 B & C for explanation of column K annotation.)

**Supplemental Table 6. Sample level RNA-seq quality control metrics.** Columns A-D provide sample name, line number, cell type (iPS or iPS-derived cardiomyocytes [cm]), and drug condition (for iPS cells, neither drug nor DMSO control was added, so condition is "ips"). Column E indicates biological replicate. For iPS cells, this indicates the same iPS clonal line harvested for RNA on a different day of culture. For cardiomyocytes this indicates the same iPS line undergoing a second 30-day cardiomyocyte differentiation process and subsequent drug treatment. Columns F and G indicate batch for library generation and sequencing pool. Columns H-N provide outputs from QC tools. For each output, the tool name is provided in parentheses.

| Table S1 | | | | | | | | | |
|---|---|---|---|---|---|---|---|---|---|
| ID | Age | Gender | Race | Southeast Asian | Other | Ethnicity | Disease | LVEF (%) | IVSD (cm) |
| 67 | 60.0 | Female | White | | | Not Hispanic or Latino | HCM | 80.00% | 1.5 |
| 70 | 24.3 | Male | White | | | Hispanic or Latino | HCM | 65.00% | 2.5 |
| 283 | 50.9 | Female | White | | | Not Hispanic or Latino | Healthy Control | | |
| 284 | 41.2 | Female | White | | | Not Hispanic or Latino | Healthy Control | | |
| 287 | 47.6 | Female | White | | | Not Hispanic or Latino | HCM | 67.10% | 1.7 |
| 289 | 67.1 | Male | White | | | Not Hispanic or Latino | HCM | 68.50% | 1.5 |
| 295 | 77.1 | Male | White | | | Not Hispanic or Latino | Healthy Control | | |
| 297 | 52.9 | Male | White | | | Not Hispanic or Latino | Healthy Control | | |
| 298 | 70.2 | Female | White | | | Not Hispanic or Latino | HCM | 67.50% | 2.2 |
| 301 | 73.5 | Male | White | | | Not Hispanic or Latino | HCM | 60.90% | 1.3 |
| 304 | 32.8 | Female | White | | | Not Hispanic or Latino | LVNC (not clinically diagnosed) | 44.80% | 0.86 |
| 310 | 73.3 | Male | African American | | | Not Hispanic or Latino | HCM | 58.50% | 1.4 |
| 319 | 87.1 | Female | White | | | Not Hispanic or Latino | HCM | 70.00% | 1.9 |
| 320 | 57.6 | Female | White | | | Not Hispanic or Latino | DCM | 62.00% | 0.89 |
| 333 | 32.1 | Male | White | | | Not Hispanic or Latino | HCM | 66.70% | 1.6 |
| 334 | 63.8 | Male | African American | | | Unknown | HCM | 56.40% | 1.3 |
| 338 | 63.2 | Male | White | | | Not Hispanic or Latino | DCM | 55.00% | 0.94 |
| 352 | 42.5 | Female | White | | | Not Hispanic or Latino | HCM | 48.10% | 1.8 |
| 356 | 48.7 | Male | White | | | Not Hispanic or Latino | HCM | 83.00% | 1.5 |
| 358 | 81.3 | Male | White | | | Not Hispanic or Latino | HCM | 50.00% | 1.4 |
| 367 | 55.6 | Male | Other | | | Hispanic or Latino | DCM | 58.00% | 1.2 |
| 371 | 35.8 | Male | White | | | Not Hispanic or Latino | HCM | 65.40% | 1.5 |
| 372 | 71.8 | Male | White | | | Not Hispanic or Latino | Healthy Control | | |
| 373 | 47.1 | Male | Asian | | | Not Hispanic or Latino | HCM | 59.30% | 1.5 |
| 374 | 53.0 | Female | White | | | Not Hispanic or Latino | HCM | 68.30% | 1.3 |
| 375 | 57.4 | Male | White | | | Not Hispanic or Latino | HCM | 63.00% | 1.6 |
| 376 | 69.8 | Female | White | | | Not Hispanic or Latino | HCM | 66.00% | 1.9 |
| 378 | 30.9 | Female | White | | | Not Hispanic or Latino | DCM | 40.00% | 1 |
| 379 | 68.5 | Male | White | | | Not Hispanic or Latino | Healthy Control | | |
| 380 | 67.0 | Male | White | | | Not Hispanic or Latino | HCM | 26.90% | 0.85 |
| 386 | 46.3 | Male | White | | | Not Hispanic or Latino | HCM | 61.20% | 2.2 |
| 388 | 55.0 | Female | White | | | Not Hispanic or Latino | HCM | 57.00% | 1.9 |
| 390 | 45.6 | Female | White | | | Not Hispanic or Latino | DCM | 29.00% | 0.81 |
| 393 | 41.9 | Male | White | | | Not Hispanic or Latino | DCM | 21.00% | 0.91 |
| 394 | 35.1 | Female | African American | | | Not Hispanic or Latino | DCM | 56.00% | 0.84 |
| 395 | 55.7 | Male | White | | | Not Hispanic or Latino | HCM | 61.40% | 1.7 |
| 397 | 17.9 | Male | Asian | | | Not Hispanic or Latino | HCM | 68.30% | 2.6 |
| 398 | 52.4 | Male | Asian | | | Not Hispanic or Latino | HCM | 36.80% | 1.3 |
| 399 | 52.9 | Female | African American | | | Not Hispanic or Latino | DCM | 16.00% | 0.95 |
| 405 | 38.8 | Male | White | | | Not Hispanic or Latino | HCM | 59.00% | 2.7 |
| 411 | 60.1 | Female | White | | | Not Hispanic or Latino | DCM | 57.00% | 1.1 |
| 413 | 69.6 | Male | Other | | Middle Eastern | Not Hispanic or Latino | HCM | 57.80% | 1.8 |
| 415 | 24.5 | Male | Other | | Filipino | Not Hispanic or Latino | LVNC | 56.80% | 0.93 |
| 419 | 36.0 | Female | Asian | | | Not Hispanic or Latino | HCM | 68.20% | 1.5 |
| 420 | 56.4 | Male | White | | | Not Hispanic or Latino | HCM | 43.80% | 1.3 |
| 421 | 32.7 | Female | White | | | Not Hispanic or Latino | HCM | 27.90% | 1.4 |
| 428 | 59.6 | Female | White | | | Not Hispanic or Latino | DCM | 57.90% | 1.3 |
| 431 | 44.5 | Male | White | | | Not Hispanic or Latino | HCM | 61.10% | 2.9 |
| 437 | 56.5 | Female | White | | | Not Hispanic or Latino | DCM | 43.00% | 0.72 |
| 438 | 51.2 | Male | White | | | Not Hispanic or Latino | HCM | 56.90% | 1.8 |
| 440 | 50.3 | Male | White | | | Not Hispanic or Latino | HCM | 55.00% | 1.8 |
| 485 | 41.6 | Male | White | | | Not Hispanic or Latino | HCM | | 1.3 |
| 487 | 70.7 | Female | White | | | Not Hispanic or Latino | HCM | 70.40% | 1.2 |
| 489 | 58.3 | Male | White | | | Not Hispanic or Latino | HCM | 59.90% | 1.4 |
| 493 | 58.5 | Female | White | | | Not Hispanic or Latino | DCM | 52.00% | 0.76 |
| 494 | 57.5 | Female | White | | | Not Hispanic or Latino | DCM | 50.00% | 0.92 |
| 495 | 40.6 | Male | White | | | Not Hispanic or Latino | DCM | 57.20% | 0.83 |
| 496 | 25.4 | Male | White | | | Not Hispanic or Latino | HCM | 63.40% | 2.8 |
| 501 | 47.1 | Male | White | | | Not Hispanic or Latino | HCM | 66.70% | 1.1 |
| 502 | 36.0 | Female | White | | | Not Hispanic or Latino | HCM | 77.60% | 1.3 |

| | | | | | | | | | |
|---|---|---|---|---|---|---|---|---|---|
| 503 | 19.1 | Male | Other | | Filipino | Not Hispanic or Latino | DCM | 24.00% | 1 |
| 505 | 61.9 | Female | Asian | | | Not Hispanic or Latino | DCM | 15.00% | 1.3 |
| 511 | 51.4 | Male | White | | | Not Hispanic or Latino | HCM | 59.50% | 1.3 |
| 512 | 24.2 | Male | Other | | | Hispanic or Latino | HCM | 38.00% | 1.1 |
| 517 | 60.4 | Female | Asian | | | Not Hispanic or Latino | HCM | 61.30% | 1 |
| 519 | 42.2 | Female | Other | | Filipino | Not Hispanic or Latino | Healthy Control | | |
| 520 | 45.6 | Female | White | | | Hispanic or Latino | Healthy Control | | |
| 521 | 46.2 | Female | White | | | Not Hispanic or Latino | Other (LQT) | | |
| 522 | 1.0 | Male | White | | | Not Hispanic or Latino | Healthy Control | | |
| 523 | 46.5 | Male | White | | | Not Hispanic or Latino | Healthy Control | | |
| 524 | 11.1 | Male | White | | | Not Hispanic or Latino | Healthy Control | | |
| 525 | 7.8 | Male | White | | | Not Hispanic or Latino | Healthy Control | | |
| 526 | 9.8 | Male | White | | | Not Hispanic or Latino | Healthy Control | | |
| 530 | 56.2 | Male | White | | | Not Hispanic or Latino | HCM | 65.00% | |
| 532 | 58.7 | Male | White | | | Not Hispanic or Latino | HCM | 74.90% | 1.8 |
| 534 | 45.9 | Female | Asian | | | Not Hispanic or Latino | HCM | 70.00% | 0.72 |
| 535 | 52.4 | Male | Asian | | | Not Hispanic or Latino | DCM | 25.00% | 0.88 |
| 539 | 42.6 | Female | Asian | Yes | | Not Hispanic or Latino | DCM | 50.10% | 1.1 |
| 540 | 65.1 | Male | White | | | Not Hispanic or Latino | HCM | 55.20% | 1.9 |
| 541 | 31.2 | Male | White | | | Not Hispanic or Latino | DCM | 31.00% | 0.62 |
| 543 | 74.5 | Male | White | | | Not Hispanic or Latino | HCM | 65.00% | 1.6 |
| 544 | 55.8 | Male | White | | | Not Hispanic or Latino | HCM (not clinically diagnosed) | 69.00% | 1.8 |
| 545 | 67.1 | Female | White | | | Not Hispanic or Latino | HCM | | 2.1 |
| 546 | 44.9 | Female | White | | | Not Hispanic or Latino | HCM | 63.20% | 2.1 |
| 547 | 83.1 | Female | Other | | Filipino | Not Hispanic or Latino | HCM | 62.00% | 1.5 |
| 548 | 44.2 | Male | White | | | Not Hispanic or Latino | HCM | 62.60% | 1.6 |
| 549 | 59.6 | Male | White | | | Not Hispanic or Latino | HCM | 61.00% | 1.2 |
| 550 | 60.1 | Male | White | | | Not Hispanic or Latino | HCM | 67.00% | 1.2 |
| 551 | 18.3 | Male | White | | | Not Hispanic or Latino | DCM | 35.40% | 0.88 |
| 552 | 72.2 | Female | White | | | Not Hispanic or Latino | Other (Fabry) | 78.00% | |
| 553 | 60.0 | Male | Asian | Yes | | Not Hispanic or Latino | HCM | 56.10% | 1.4 |
| 555 | 68.9 | Male | Other | | Filipino | Not Hispanic or Latino | HCM | 70.00% | 1.5 |
| 556 | 51.2 | Male | Asian | | | Not Hispanic or Latino | HCM | 59.00% | 0.97 |
| 557 | 46.5 | Female | White | | | Not Hispanic or Latino | Healthy Control | | |
| 558 | 47.1 | Male | White | | | Not Hispanic or Latino | Healthy Control | | |
| 559 | 8.9 | Female | White | | | Not Hispanic or Latino | Other | | |
| 560 | 12.0 | Male | White | | | Not Hispanic or Latino | Other | | |
| 561 | 16.9 | Female | White | | | Not Hispanic or Latino | Healthy Control | | |
| 562 | 27.7 | Male | White | | | Not Hispanic or Latino | HCM | 73.00% | 1.5 |
| 563 | 59.0 | Male | White | | | Not Hispanic or Latino | HCM | 57.50% | 1.8 |
| 564 | 37.7 | Female | Asian | | | Not Hispanic or Latino | HCM | 60.30% | 2.7 |
| 565 | 37.3 | Male | Other | | | Hispanic or Latino | HCM | 56.90% | 1.5 |
| 566 | 45.9 | Male | African American | | | Not Hispanic or Latino | HCM | 57.20% | 2.4 |
| 567 | 72.0 | Female | White | | | Not Hispanic or Latino | HCM | 68.80% | 1.2 |
| 568 | 62.8 | Female | Other | | | Hispanic or Latino | HCM | 50.30% | |
| 569 | 87.6 | Female | White | | | Not Hispanic or Latino | HCM | 51.60% | 2.1 |
| 570 | 79.6 | Female | White | | | Not Hispanic or Latino | HCM | 67.00% | 2.1 |
| 571 | 72.0 | Female | Native American | | | Hispanic or Latino | HCM | 58.40% | 1.7 |
| 574 | 60.7 | Female | White | | | Not Hispanic or Latino | DCM | 45.00% | 0.79 |
| 576 | 87.1 | Female | White | | | Not Hispanic or Latino | HCM | 59.40% | 1.7 |
| 578 | 73.5 | Male | White | | | Not Hispanic or Latino | HCM | 72.00% | 1.5 |
| 581 | 61.3 | Female | White | | | Not Hispanic or Latino | HCM | | |
| 582 | 57.9 | Male | Asian | Yes | | Not Hispanic or Latino | HCM | | |
| 586 | 56.4 | Female | White | | | Not Hispanic or Latino | DCM | 52.60% | 0.94 |
| 591 | 24.4 | Female | White | | | Not Hispanic or Latino | HCM | 67.60% | 1 |
| 592 | 55.2 | Male | White | | | Not Hispanic or Latino | HCM | 64.30% | 1.7 |
| 593 | 28.7 | Male | White | | | Not Hispanic or Latino | HCM | 55.60% | 0.81 |
| 596 | 38.7 | Male | White | | | Not Hispanic or Latino | DCM | 56.00% | 0.89 |
| 598 | 65.7 | Female | Other | | | Hispanic or Latino | HCM | 66.20% | 1.6 |
| 599 | 71.7 | Female | White | | | Not Hispanic or Latino | HCM | 65.00% | 2.3 |
| 601 | 36.8 | Male | White | | | Not Hispanic or Latino | DCM | 29.60% | 0.86 |
| 603 | 32.5 | Male | White | | | Not Hispanic or Latino | HCM (not clinically diagnosed) | 65.60% | 2.1 |

| | | | | | | | | | |
|---|---|---|---|---|---|---|---|---|---|
| 605 | 56.7 | Male | African American | | | Not Hispanic or Latino | DCM | 35.20% | 0.78 |
| 607 | 44.2 | Male | Other | | Maternal European ancestry, paternal Egyptian ancestry | Not Hispanic or Latino | HCM | 66.10% | 1.7 |
| 612 | 69.3 | Male | White | | | Not Hispanic or Latino | HCM | | 2 |
| 613 | 34.9 | Male | White | | | Not Hispanic or Latino | HCM | | 2 |
| 614 | 82.0 | Female | White | | | Not Hispanic or Latino | HCM | 70.70% | 2 |
| 615 | 29.1 | Female | White | | | Not Hispanic or Latino | Healthy Control | | |
| 618 | 55.1 | Male | White | | | Not Hispanic or Latino | HCM | 58.50% | 2.9 |
| 619 | 48.4 | Male | African American | | | Not Hispanic or Latino | HCM | 67.90% | 1.5 |
| 621 | 36.4 | Male | White | | | Not Hispanic or Latino | Healthy Control | | |
| 622 | 58.6 | Male | White | | | Not Hispanic or Latino | HCM | 30.80% | 1.5 |
| 624 | 45.7 | Female | White | | | Not Hispanic or Latino | DCM | 35.20% | 1.1 |
| 625 | 51.3 | Male | Other | | Paternal ancestry: Native American, Black; maternal ancestry: European | Not Hispanic or Latino | HCM | 72.00% | 2.1 |
| 628 | 54.2 | Female | White | | | Not Hispanic or Latino | DCM | 45.00% | 0.97 |
| 629 | 25.7 | Male | White | | | Not Hispanic or Latino | LVNC (not clinically diagnosed) | 71.00% | |
| 632 | 29.8 | Male | White | | | Not Hispanic or Latino | Healthy Control | | |
| 634 | 54.2 | Male | Other | | | Hispanic or Latino | HCM | 73.90% | 1.6 |
| 638 | 29.0 | Female | Asian | | | Not Hispanic or Latino | HCM | 75.00% | 1.7 |
| 643 | 60.1 | Male | Asian | | | Not Hispanic or Latino | DCM | 37.00% | 0.99 |
| 645 | 48.9 | Male | White | | | Not Hispanic or Latino | HCM | 67.70% | 1.5 |
| 649 | 63.5 | Female | White | | | Not Hispanic or Latino | HCM | 31.50% | 1.4 |
| 650 | 46.9 | Female | White | | | Unknown | DCM | 45.40% | 0.7 |
| 653 | 53.6 | Male | White | | | Not Hispanic or Latino | HCM | | |
| 656 | 70.0 | Female | White | | | Not Hispanic or Latino | HCM | 55.80% | 1.2 |
| 657 | 57.9 | Male | White | | | Not Hispanic or Latino | HCM | 55.50% | 1.3 |
| 658 | 61.0 | Female | White | | | Not Hispanic or Latino | LVNC (not clinically diagnosed) | 47.70% | 0.78 |
| 659 | 59.5 | Female | African American | | | Not Hispanic or Latino | HCM | 60.20% | 1.9 |
| 662 | 30.2 | Male | White | | | Not Hispanic or Latino | HCM | 45.60% | 1.5 |
| 668 | 51.2 | Male | White | | | Not Hispanic or Latino | DCM | 25.00% | 1.2 |
| 673 | 58.9 | Female | Other | | | Hispanic or Latino | HCM | 65.90% | 1.5 |
| 676 | 55.4 | Female | Other | | | Hispanic or Latino | HCM | 71.00% | 0.02 |
| 677 | 26.9 | Male | White | | | Not Hispanic or Latino | DCM | 50.60% | 0.77 |
| 684 | 27.8 | Male | Asian | | | Not Hispanic or Latino | LVNC | 42.70% | 0.64 |
| 686 | 50.8 | Female | White | | | Not Hispanic or Latino | HCM | | 1.4 |
| 693 | 22.9 | Male | White | | | Not Hispanic or Latino | HCM | 62.10% | 2.5 |
| 697 | 24.7 | Male | White | | | Not Hispanic or Latino | HCM | 78.60% | 1.9 |
| 709 | 59.9 | Female | White | | | Not Hispanic or Latino | DCM | 17.00% | 0.89 |
| 710 | 27.5 | Male | Asian | | | Not Hispanic or Latino | DCM | 20.30% | 1.1 |
| 711 | 67.3 | Female | White | | | Not Hispanic or Latino | HCM | 65.90% | 1.5 |
| 712 | 72.5 | Female | White | | | Not Hispanic or Latino | HCM | 74.70% | 1.8 |
| 715 | 59.8 | Female | White | | | Not Hispanic or Latino | DCM | 37.20% | 1.2 |
| 721 | 46.3 | Male | White | | | Not Hispanic or Latino | DCM | 53.00% | 0.86 |
| 728 | 56.7 | Female | White | | | Not Hispanic or Latino | DCM | 63.70% | 0.86 |
| 731 | 45.9 | Male | White | | | Not Hispanic or Latino | Healthy Control | | |
| 732 | 22.1 | Male | White | | | Not Hispanic or Latino | Other | | |
| 733 | 71.3 | Female | White | | | Not Hispanic or Latino | Other | | |
| 734 | 44.2 | Female | White | | | Not Hispanic or Latino | Other | | |
| 735 | 17.0 | Male | White | | | Not Hispanic or Latino | Other | | |
| 738 | 73.5 | Female | White | | | Not Hispanic or Latino | HCM | 64.90% | 1.8 |
| 741 | 55.8 | Male | White | | | Not Hispanic or Latino | DCM | 22.10% | 0.86 |
| 751 | 56.5 | Female | White | | | Not Hispanic or Latino | DCM | 45.20% | 0.9 |
| 754 | 79.2 | Male | White | | | Not Hispanic or Latino | DCM | 26.70% | 0.93 |
| 766 | 35.4 | Female | White | | | Not Hispanic or Latino | DCM | 40.60% | 1.1 |
| 769 | 39.8 | Female | White | | | Not Hispanic or Latino | DCM | 44.90% | 0.7 |
| 772 | 49.2 | Male | Other | | | Hispanic or Latino | DCM | 47.70% | 0.97 |
| 780 | 63.9 | Male | African American | | | Not Hispanic or Latino | DCM | 50.00% | 1.2 |
| 783 | 81.7 | Female | White | | | Not Hispanic or Latino | DCM | 27.10% | 0.76 |
| 787 | 52.0 | Female | African American | | | Not Hispanic or Latino | DCM | 23.80% | 1.3 |
| 788 | 56.2 | Male | White | | | Not Hispanic or Latino | DCM | 22.10% | 0.67 |
| 793 | 77.0 | Male | White | | | Not Hispanic or Latino | LVNC (not clinically diagnosed) | 66.50% | 1.4 |

| | | | | | | | | | |
|---|---|---|---|---|---|---|---|---|---|
| 794 | 62.3 | Female | White | | | Not Hispanic or Latino | LVNC | 54.60% | 0.85 |
| 799 | 36.6 | Male | White | | | Not Hispanic or Latino | LVNC | 68.90% | 0.95 |
| 803 | 28.5 | Male | White | | | Not Hispanic or Latino | LVNC | 43.20% | 0.71 |
| 808 | 38.2 | Female | White | | | Not Hispanic or Latino | LVNC | 37.80% | 0.6 |
| 811 | 19.7 | Female | White | | | Not Hispanic or Latino | DCM (not clinically diagnosed) | 64.90% | 0.89 |
| 813 | 49.6 | Female | White | | | Not Hispanic or Latino | DCM | 30.10% | 0.82 |
| 814 | 62.2 | Male | White | | | Not Hispanic or Latino | DCM | 55.50% | 1.1 |
| 815 | 41.9 | Male | White | | | Not Hispanic or Latino | DCM | 51.70% | 0.93 |
| 817 | 65.8 | Male | Other | | | Hispanic or Latino | DCM | 64.90% | 1.1 |
| 820 | 50.0 | Male | White | | | Not Hispanic or Latino | Healthy Control | | |
| 821 | 68.4 | Female | African American | | | Not Hispanic or Latino | Healthy Control | | |
| 822 | 62.1 | Female | White | | | Not Hispanic or Latino | Healthy Control | | |
| 823 | 62.3 | Female | White | | | Not Hispanic or Latino | Healthy Control | | |
| 827 | 58.0 | Female | White | | | Not Hispanic or Latino | Healthy Control | | |
| 839 | 55.0 | Male | Asian | Yes | | Not Hispanic or Latino | Healthy Control | | |
| 844 | 36.3 | Male | African American | | | Not Hispanic or Latino | DCM | 16.80% | 1.2 |
| 851 | 57.7 | Female | White | | | Not Hispanic or Latino | DCM | 20.10% | 1 |
| 852 | 38.2 | Female | Unknown | | | Unknown | DCM | 34.10% | 1 |
| 854 | 62.2 | Female | Other | | | Hispanic or Latino | Healthy Control | | |
| 855 | 52.2 | Female | White | | | Not Hispanic or Latino | Healthy Control | | |
| 856 | 63.2 | Female | Other | | | Hispanic or Latino | Healthy Control | | |
| 857 | 62.4 | Male | Asian | | | Not Hispanic or Latino | Healthy Control | | |
| 860 | 53.6 | Male | Asian | | | Not Hispanic or Latino | Healthy Control | | |
| 861 | 49.4 | Female | White | | | Not Hispanic or Latino | Healthy Control | | |
| 862 | 65.4 | Male | White | | | Not Hispanic or Latino | Healthy Control | | |
| 868 | 55.0 | Female | Other | | | Hispanic or Latino | Healthy Control | | |
| 869 | 53.0 | Female | White | | | Not Hispanic or Latino | Healthy Control | | |
| 875 | 36.8 | Female | Other | | | Not Hispanic or Latino | DCM | 62.40% | 0.9 |
| 885 | 46.9 | Male | White | | | Not Hispanic or Latino | Healthy Control | | |
| 886 | 5.4 | Male | White | | | Not Hispanic or Latino | DCM | 52.00% | 0.51 |
| 887 | 57.0 | Female | White | | | Not Hispanic or Latino | LVNC | 42.10% | 0.89 |
| 888 | 55.1 | Female | Other | | | Hispanic or Latino | DCM (not clinically diagnosed) | 30.00% | 0.68 |
| 910 | 69.1 | Male | White | | | Not Hispanic or Latino | DCM | 28.70% | 1 |
| 912 | 53.9 | Male | White | | | Not Hispanic or Latino | DCM | 67.00% | 0.92 |
| 914 | 63.8 | Male | African American | | | Not Hispanic or Latino | DCM | 18.90% | 0.8 |
| 915 | 63.4 | Male | White | | | Not Hispanic or Latino | DCM | 59.70% | 1.2 |
| 916 | 72.0 | Male | Other | | | Hispanic or Latino | Other | | |
| 917 | 48.3 | Female | White | | | Not Hispanic or Latino | DCM (not clinically diagnosed) | 64.90% | 0.88 |
| 919 | 62.2 | Male | Asian | | | Not Hispanic or Latino | Other | | |
| 920 | 43.0 | Male | White | | | Not Hispanic or Latino | DCM | 39.60% | 0.6 |
| 923 | 54.0 | Female | African American | | | Not Hispanic or Latino | Other | | |
| 925 | 25.7 | Male | Asian | | | Not Hispanic or Latino | DCM | 57.80% | 1.2 |
| 927 | 54.2 | Female | African American | | | Not Hispanic or Latino | Other | | |
| 928 | 73.8 | Female | Asian | Yes | | Not Hispanic or Latino | DCM | 38.90% | 0.75 |
| 930 | 58.2 | Male | African American | | | Not Hispanic or Latino | Other | | |
| 931 | 68.1 | Male | Other | | | Hispanic or Latino | Other | | |
| 932 | 48.0 | Female | White | | | Not Hispanic or Latino | LVNC | 38.40% | 0.85 |
| 933 | 42.3 | Female | Asian | Yes | | Not Hispanic or Latino | Healthy Control | | |
| 934 | 63.7 | Male | Other | | | Hispanic or Latino | Other | | |
| 935 | 54.3 | Male | Other | | Palestinian | Not Hispanic or Latino | DCM | 45.10% | 1.2 |
| 936 | 63.1 | Male | African American | | | Not Hispanic or Latino | Other | | |
| 937 | 64.8 | Male | White | | | Not Hispanic or Latino | LVNC | 40.20% | 0.82 |
| 938 | 35.8 | Female | White | | | Not Hispanic or Latino | DCM | 55.30% | 0.87 |
| 941 | 51.8 | Male | Other | | | Hispanic or Latino | Other | | |
| 942 | 55.8 | Male | Other | | | Hispanic or Latino | DCM | 28.00% | 1 |
| 944 | 46.9 | Male | Asian | | | Not Hispanic or Latino | Healthy Control | | |
| 946 | 56.1 | Male | Asian | Yes | | Not Hispanic or Latino | DCM | 21.80% | 1 |
| 950 | 55.8 | Male | African American | | | Not Hispanic or Latino | Other | | |
| 954 | 55.7 | Female | Pacific Islander | | | Not Hispanic or Latino | DCM | 19.90% | 0.8 |
| 955 | 68.6 | Female | White | | | Not Hispanic or Latino | Healthy Control | | |
| 957 | 78.2 | Female | Other | | | Hispanic or Latino | DCM | 21.40% | 0.89 |
| 959 | 61.1 | Male | Other | | | Hispanic or Latino | Other | | |

| | | | | | | | | | |
|---|---|---|---|---|---|---|---|---|---|
| 960 | 65.3 | Male | White | | | Not Hispanic or Latino | Healthy Control | | |
| 961 | 82.8 | Male | Asian | | | Not Hispanic or Latino | Other | | |
| 962 | 65.0 | Female | Asian | | | Not Hispanic or Latino | Healthy Control | | |
| 963 | 48.5 | Female | African American | | | Not Hispanic or Latino | DCM | 45.10% | 0.79 |
| 964 | 66.4 | Female | African American | | | Not Hispanic or Latino | Other | | |
| 965 | 61.1 | Female | White | | | Not Hispanic or Latino | DCM | 45.40% | 1.1 |
| 967 | 60.1 | Female | Asian | | | Not Hispanic or Latino | Healthy Control | | |
| 969 | 42.9 | Female | African American | | | Not Hispanic or Latino | DCM (not clinically diagnosed) | 23.80% | 1 |
| 974 | 35.9 | Female | Other | | | Hispanic or Latino | DCM | 15.20% | 0.84 |
| 979 | 72.9 | Male | Asian | | | Not Hispanic or Latino | Other | | |
| 986 | 33.3 | Male | White | | | Not Hispanic or Latino | DCM | 40.10% | 0.88 |
| 989 | 49.6 | Female | Asian | | | Not Hispanic or Latino | Healthy Control | | |
| 990 | 57.2 | Female | Asian | | | Not Hispanic or Latino | Healthy Control | | |
| 991 | 59.9 | Male | Asian | | | Not Hispanic or Latino | Healthy Control | | |
| 995 | 60.1 | Male | White | | | Not Hispanic or Latino | DCM | 63.30% | 0.82 |
| 1074 | 71.0 | Male | White | | | Not Hispanic or Latino | DCM | | |
| 2002 | 66.1 | Female | White | | | Not Hispanic or Latino | Healthy Control | | |
| 2003 | 80.4 | Female | Other | | Filipino | Not Hispanic or Latino | Healthy Control | | |
| 2006 | 65.4 | Female | White | | | Not Hispanic or Latino | DCM | 22.80% | 0.8 |
| 2007 | 71.7 | Male | White | | | Not Hispanic or Latino | DCM | 25.00% | 0.9 |
| 2009 | 36.0 | Male | Asian | Yes | | Not Hispanic or Latino | Healthy Control | | |
| 2012 | 63.9 | Female | White | | | Not Hispanic or Latino | DCM | 17.70% | 1.3 |
| 2013 | 70.3 | Male | White | | | Not Hispanic or Latino | Healthy Control | | |
| 2015 | 51.1 | Female | Native Hawaiian | | | Not Hispanic or Latino | DCM | 55.90% | 1.6 |
| 2020 | 54.5 | Female | White | | | Not Hispanic or Latino | DCM | 50.10% | 0.81 |
| 2025 | 37.9 | Male | White | | | Not Hispanic or Latino | DCM | 59.30% | 0.97 |
| 2032 | 65.1 | Female | Asian | | | Not Hispanic or Latino | Other | | |
| 2035 | 78.9 | Male | Asian | | | Not Hispanic or Latino | Other | | |
| 2036 | 23.8 | Male | White | | | Not Hispanic or Latino | DCM | 50.20% | 0.78 |
| 2038 | 61.8 | Female | White | | | Not Hispanic or Latino | Healthy Control | | |
| 2039 | 60.9 | Male | White | | | Not Hispanic or Latino | DCM | 36.40% | 0.91 |
| 2051 | 69.8 | Male | Asian | | | Not Hispanic or Latino | Other | | |
| 2055 | 53.9 | Male | White | | | Not Hispanic or Latino | Healthy Control | | |
| 2059 | 37.4 | Male | Other | | | Hispanic or Latino | DCM | 23.30% | 1 |
| 2061 | 79.6 | Male | Asian | | | Not Hispanic or Latino | Other | | |
| 2062 | 47.6 | Male | Asian | Yes | | Not Hispanic or Latino | DCM | 24.20% | 0.8 |
| 2063 | 56.5 | Male | Other | | | Hispanic or Latino | Other | | |
| 2064 | 29.1 | Male | African American | | | Not Hispanic or Latino | LVNC | 68.60% | 1 |
| 2065 | 52.1 | Male | Asian | | | Not Hispanic or Latino | Other | | |
| 2073 | 54.7 | Male | White | | | Not Hispanic or Latino | Healthy Control | | |
| 2086 | 67.7 | Male | White | | | Not Hispanic or Latino | Healthy Control | | |
| 2136 | 50.8 | Female | White | | | Not Hispanic or Latino | Healthy Control | | |
| 2137 | 51.1 | Male | White | | | Not Hispanic or Latino | Healthy Control | | |
| 2138 | 20.5 | Male | White | | | Not Hispanic or Latino | Healthy Control | | |
| 2156 | 62.0 | Female | White | | | Not Hispanic or Latino | Other | | |
| 2157 | 66.0 | Male | Asian | East Asian | | Not Hispanic or Latino | Other | | |
| 2159 | 23.7 | Female | White | | | Not Hispanic or Latino | Healthy Control | | |
| 2161 | 63.0 | Male | Asian | East Asian | | Not Hispanic or Latino | Other | | |
| 2162 | 60.0 | Female | Asian | East Asian | | Not Hispanic or Latino | Other | | |
| 2163 | 62.0 | Female | Asian | East Asian | | Not Hispanic or Latino | Other | | |
| 2167 | 57.1 | Male | White | | | Not Hispanic or Latino | Healthy Control | | |
| 2184 | 27.3 | Male | White | | | Not Hispanic or Latino | Healthy Control | | |
| 2186 | 25.2 | Female | Other | | | Hispanic or Latino | Healthy Control | | |
| 2189 | 66.0 | Female | Asian | East Asian | | Not Hispanic or Latino | Healthy Control | | |
| 2194 | 73.0 | Female | Asian | East Asian | | Not Hispanic or Latino | Healthy Control | | |
| 2197 | 49.0 | Male | Asian | East Asian | | | Healthy Control | | |
| 2198 | 43.0 | Female | Asian | East Asian | | Not Hispanic or Latino | Healthy Control | | |

**Table S2**

| Parameter | Average | Min | Max |
|---|---|---|---|
| Total SNPs+Indels (millions) | 4.28 | 4.14 | 5.14 |
| TiTv dbSNP | 2.11 | 2.1 | 2.11 |
| TiTv novel | 1.23 | 1.16 | 1.32 |
| Ins/Del dbSNP | 0.89 | 0.88 | 0.9 |
| Ins/Del novel | 0.65 | 0.62 | 0.71 |
| SNP reference bias | 0.54 | 0.53 | 0.55 |
| Coverage | 28.2 | 18.2 | 44.1 |

**Table S3**

| Sample | Mean Coverage | TOTAL_SNPS | TOTAL_INDELS | TOTAL SNPS+INDELS | DBSNP_TITV | NOVEL_TITV | DBSNP_INS_DEL_RA | NOVEL_INS_DEL_RA | SNP_REFERENCE_BIAS |
|---|---|---|---|---|---|---|---|---|---|
| 67 | 25.44 | 3865486 | 388247 | 4.253733 | 2.107593 | 1.23858 | 0.890661 | 0.638943 | 0.537844 |
| 70 | 23.19 | 3828412 | 382274 | 4.210686 | 2.106071 | 1.228261 | 0.891072 | 0.644689 | 0.534503 |
| 283 | 36.17 | 3829867 | 380709 | 4.210576 | 2.110035 | 1.200518 | 0.894566 | 0.643686 | 0.54179 |
| 284 | 31.89 | 3871754 | 386279 | 4.258033 | 2.107034 | 1.227879 | 0.889223 | 0.653137 | 0.537624 |
| 287 | 34.52 | 3882478 | 386449 | 4.268927 | 2.105272 | 1.242289 | 0.892151 | 0.647025 | 0.538323 |
| 289 | 37.34 | 3857871 | 383927 | 4.241798 | 2.10567 | 1.23896 | 0.890383 | 0.651265 | 0.538698 |
| 295 | 29.57 | 3796688 | 377214 | 4.173902 | 2.107108 | 1.240403 | 0.888625 | 0.64378 | 0.537759 |
| 297 | 31.98 | 3809100 | 380368 | 4.189468 | 2.104918 | 1.239576 | 0.891807 | 0.637438 | 0.536682 |
| 298 | 37.4 | 3876330 | 386103 | 4.262433 | 2.106327 | 1.249724 | 0.891905 | 0.650325 | 0.539467 |
| 301 | 35.73 | 3851380 | 384547 | 4.235927 | 2.106246 | 1.212786 | 0.894259 | 0.631118 | 0.540042 |
| 304 | 27.54 | 3829586 | 381061 | 4.210647 | 2.108784 | 1.202409 | 0.892226 | 0.641577 | 0.539387 |
| 310 | 30.35 | 4567574 | 466073 | 5.033647 | 2.108913 | 1.305366 | 0.901376 | 0.692395 | 0.533929 |
| 319 | 33.7 | 3842382 | 384319 | 4.226701 | 2.105461 | 1.225773 | 0.888295 | 0.647666 | 0.539921 |
| 320 | 27.66 | 3828211 | 381067 | 4.209278 | 2.107453 | 1.208738 | 0.89129 | 0.631135 | 0.537449 |
| 333 | 33.08 | 3846001 | 385641 | 4.231642 | 2.105652 | 1.217931 | 0.889489 | 0.639115 | 0.536976 |
| 334 | 26.94 | 4539149 | 459202 | 4.998351 | 2.108793 | 1.274533 | 0.900696 | 0.692829 | 0.539129 |
| 338 | 25.41 | 3771298 | 376690 | 4.147988 | 2.107833 | 1.231012 | 0.892455 | 0.665522 | 0.538626 |
| 352 | 27.69 | 3851862 | 382114 | 4.233976 | 2.110064 | 1.223389 | 0.891415 | 0.65331 | 0.540764 |
| 356 | 26.48 | 3817058 | 380414 | 4.197472 | 2.108222 | 1.215969 | 0.894968 | 0.643274 | 0.541525 |
| 358 | 26.88 | 3830591 | 381725 | 4.212316 | 2.105795 | 1.212063 | 0.893891 | 0.652212 | 0.541063 |
| 367 | 24.63 | 3883036 | 387434 | 4.27047 | 2.107139 | 1.244366 | 0.894333 | 0.649871 | 0.541837 |
| 371 | 31.33 | 3806069 | 377211 | 4.18328 | 2.107589 | 1.202123 | 0.891575 | 0.634756 | 0.539503 |
| 372 | 29 | 3838397 | 380415 | 4.218812 | 2.107548 | 1.230043 | 0.89707 | 0.63331 | 0.541775 |
| 373 | 28.9 | 3810013 | 382101 | 4.192114 | 2.1009 | 1.25359 | 0.895177 | 0.653 | 0.540677 |
| 374 | 25.36 | 3803379 | 378204 | 4.181583 | 2.109199 | 1.2174 | 0.893731 | 0.633373 | 0.543782 |
| 375 | 28.2 | 3815187 | 380136 | 4.195323 | 2.105844 | 1.218664 | 0.891363 | 0.655241 | 0.540591 |
| 376 | 24.96 | 3808029 | 380024 | 4.188053 | 2.109415 | 1.185671 | 0.890712 | 0.640431 | 0.539756 |
| 378 | 26.11 | 3809831 | 380944 | 4.190775 | 2.10438 | 1.234936 | 0.897818 | 0.660326 | 0.542999 |
| 379 | 25.74 | 3816406 | 379607 | 4.196013 | 2.106534 | 1.206741 | 0.893598 | 0.646443 | 0.54365 |
| 380 | 33.5 | 3799298 | 377974 | 4.177272 | 2.106133 | 1.215582 | 0.891763 | 0.634943 | 0.5418 |
| 386 | 25.81 | 3862996 | 385764 | 4.24876 | 2.103686 | 1.225071 | 0.892932 | 0.654074 | 0.540708 |
| 388 | 36.7 | 3859489 | 387226 | 4.246715 | 2.107124 | 1.216851 | 0.889108 | 0.642081 | 0.540811 |
| 390 | 26.19 | 3818074 | 381305 | 4.199379 | 2.105701 | 1.203953 | 0.891328 | 0.645467 | 0.541641 |
| 393 | 28.07 | 3789902 | 377128 | 4.16703 | 2.1074 | 1.200328 | 0.892675 | 0.630952 | 0.541161 |
| 394 | 27.21 | 4442563 | 448050 | 4.890613 | 2.107751 | 1.256437 | 0.897492 | 0.68364 | 0.533704 |
| 395 | 26.19 | 3827900 | 381402 | 4.209302 | 2.106218 | 1.212087 | 0.892995 | 0.651665 | 0.542105 |
| 397 | 33.93 | 3824652 | 385079 | 4.209731 | 2.100397 | 1.237899 | 0.895077 | 0.63608 | 0.538475 |
| 398 | 27.85 | 3822294 | 386824 | 4.209118 | 2.102102 | 1.219922 | 0.897102 | 0.638609 | 0.538087 |
| 399 | 27.21 | 4443232 | 447455 | 4.890687 | 2.110574 | 1.270187 | 0.894918 | 0.681006 | 0.531214 |
| 405 | 26.84 | 3831139 | 381983 | 4.213122 | 2.108506 | 1.210883 | 0.895527 | 0.63806 | 0.544523 |
| 411 | 25.99 | 3826925 | 382148 | 4.209073 | 2.107811 | 1.203943 | 0.892342 | 0.642667 | 0.539379 |
| 413 | 25.11 | 3893890 | 390137 | 4.284027 | 2.110658 | 1.238391 | 0.894336 | 0.646681 | 0.537091 |
| 415 | 26.3 | 3783216 | 377237 | 4.160453 | 2.104172 | 1.247392 | 0.896798 | 0.649642 | 0.545155 |
| 419 | 28.22 | 3838520 | 384924 | 4.223444 | 2.105307 | 1.235152 | 0.89693 | 0.647422 | 0.540715 |
| 420 | 28.13 | 3801878 | 380055 | 4.181933 | 2.111993 | 1.211612 | 0.896068 | 0.644403 | 0.538099 |
| 421 | 23.18 | 3803803 | 381998 | 4.185801 | 2.109551 | 1.207608 | 0.893596 | 0.642835 | 0.541241 |
| 428 | 25.13 | 3828988 | 383660 | 4.212648 | 2.105813 | 1.205958 | 0.892658 | 0.652015 | 0.543261 |
| 431 | 29.15 | 3807726 | 380675 | 4.188401 | 2.104029 | 1.194757 | 0.894054 | 0.639604 | 0.542527 |
| 437 | 21.86 | 3826730 | 385295 | 4.212025 | 2.102667 | 1.270204 | 0.89772 | 0.64711 | 0.544585 |
| 438 | 25.97 | 3767731 | 378286 | 4.146017 | 2.106493 | 1.215289 | 0.888435 | 0.635732 | 0.539752 |
| 440 | 30.63 | 3805329 | 383152 | 4.188481 | 2.107261 | 1.203936 | 0.890269 | 0.641613 | 0.542092 |
| 485 | 25.05 | 3802239 | 377851 | 4.18009 | 2.105147 | 1.216388 | 0.890806 | 0.63851 | 0.541127 |
| 487 | 26.49 | 3834297 | 381940 | 4.216237 | 2.110778 | 1.209485 | 0.892065 | 0.643414 | 0.539073 |
| 489 | 28.03 | 3790860 | 381810 | 4.17267 | 2.107582 | 1.207447 | 0.889059 | 0.649689 | 0.540451 |
| 493 | 29.53 | 3822639 | 382163 | 4.204802 | 2.108175 | 1.214611 | 0.892799 | 0.641319 | 0.539926 |
| 494 | 30.74 | 3851757 | 384228 | 4.235985 | 2.104174 | 1.208384 | 0.892503 | 0.664936 | 0.545395 |
| 495 | 28.13 | 3800731 | 379065 | 4.179796 | 2.103594 | 1.209282 | 0.891596 | 0.644301 | 0.540053 |
| 496 | 24.93 | 3835848 | 381616 | 4.217464 | 2.103464 | 1.224702 | 0.894555 | 0.64255 | 0.542548 |
| 501 | 26.79 | 3805984 | 379187 | 4.185171 | 2.108168 | 1.214124 | 0.895 | 0.646061 | 0.541465 |
| 502 | 28.38 | 3816016 | 379792 | 4.195808 | 2.109008 | 1.196669 | 0.89343 | 0.630074 | 0.541394 |
| 503 | 26.98 | 3812483 | 386036 | 4.198519 | 2.102646 | 1.238666 | 0.895694 | 0.660883 | 0.539001 |
| 505 | 28.62 | 3842253 | 384278 | 4.226531 | 2.102267 | 1.224656 | 0.898917 | 0.644978 | 0.541134 |
| 511 | 34.69 | 3814384 | 384474 | 4.198858 | 2.103256 | 1.201406 | 0.887418 | 0.659274 | 0.536799 |
| 512 | 29.19 | 4173092 | 422220 | 4.595312 | 2.106993 | 1.224877 | 0.895594 | 0.663914 | 0.538195 |
| 517 | 31.59 | 3850802 | 388315 | 4.239117 | 2.103965 | 1.22385 | 0.89824 | 0.654465 | 0.542618 |
| 519 | 30.23 | 3887689 | 390804 | 4.278493 | 2.107442 | 1.262333 | 0.898159 | 0.658905 | 0.543801 |
| 520 | 27.79 | 3820731 | 384084 | 4.204815 | 2.108835 | 1.206372 | 0.893628 | 0.658013 | 0.543652 |
| 521 | 26.95 | 3785202 | 380925 | 4.166127 | 2.107634 | 1.194008 | 0.893517 | 0.646585 | 0.545334 |

| | | | | | | | | | |
|---|---|---|---|---|---|---|---|---|---|
| 522 | 25.97 | 3836231 | 384139 | 4.22037 | 2.104721 | 1.158367 | 0.892003 | 0.637142 | 0.545678 |
| 523 | 27.82 | 3793839 | 382572 | 4.176411 | 2.107399 | 1.20372 | 0.892614 | 0.645277 | 0.542637 |
| 524 | 23.42 | 3830452 | 386266 | 4.216718 | 2.110057 | 1.225578 | 0.891775 | 0.65153 | 0.538526 |
| 525 | 27.54 | 3825651 | 385777 | 4.211428 | 2.107736 | 1.194155 | 0.891874 | 0.639347 | 0.537862 |
| 526 | 32.1 | 3850623 | 386808 | 4.237431 | 2.10756 | 1.200182 | 0.891663 | 0.654872 | 0.54312 |
| 530 | 24.96 | 3825035 | 384629 | 4.209664 | 2.107619 | 1.236389 | 0.892922 | 0.649051 | 0.539035 |
| 532 | 28.9 | 3834270 | 387925 | 4.222195 | 2.104662 | 1.222159 | 0.889006 | 0.650445 | 0.539012 |
| 534 | 30.94 | 3839582 | 387445 | 4.227027 | 2.100565 | 1.229256 | 0.894893 | 0.639619 | 0.545071 |
| 535 | 32.76 | 3830711 | 387805 | 4.218516 | 2.103791 | 1.211751 | 0.895592 | 0.647654 | 0.542611 |
| 539 | 26.84 | 3908943 | 395044 | 4.303987 | 2.104456 | 1.250362 | 0.890793 | 0.647418 | 0.539707 |
| 540 | 26.29 | 3816922 | 384730 | 4.201652 | 2.106054 | 1.226716 | 0.89269 | 0.658613 | 0.540805 |
| 541 | 22.95 | 3807942 | 384818 | 4.19276 | 2.106595 | 1.213495 | 0.895739 | 0.658285 | 0.536662 |
| 543 | 29.42 | 3860519 | 389094 | 4.249613 | 2.109464 | 1.282401 | 0.893189 | 0.673575 | 0.542947 |
| 544 | 32.85 | 3805785 | 383016 | 4.188801 | 2.104809 | 1.207845 | 0.890776 | 0.64982 | 0.542579 |
| 545 | 35.99 | 3897834 | 391611 | 4.289445 | 2.109452 | 1.213006 | 0.893553 | 0.645139 | 0.54324 |
| 546 | 32.86 | 3833700 | 385444 | 4.219144 | 2.107427 | 1.212437 | 0.891248 | 0.638137 | 0.539308 |
| 547 | 29.37 | 3909019 | 393870 | 4.302889 | 2.103948 | 1.25148 | 0.891123 | 0.63492 | 0.546762 |
| 548 | 34.24 | 3808719 | 382718 | 4.191437 | 2.105549 | 1.206377 | 0.892468 | 0.640486 | 0.542945 |
| 549 | 28.47 | 3817118 | 385032 | 4.20215 | 2.106903 | 1.220783 | 0.892073 | 0.652719 | 0.537423 |
| 550 | 25.21 | 3787803 | 380149 | 4.167952 | 2.105273 | 1.212243 | 0.891055 | 0.635496 | 0.539559 |
| 551 | 27.44 | 3795814 | 380273 | 4.176087 | 2.104869 | 1.205229 | 0.889416 | 0.632515 | 0.539188 |
| 552 | 26.03 | 3821429 | 383357 | 4.204786 | 2.108959 | 1.211425 | 0.891284 | 0.636873 | 0.542022 |
| 553 | 28.81 | 3893195 | 391950 | 4.285145 | 2.103827 | 1.235532 | 0.896148 | 0.641754 | 0.539449 |
| 555 | 23.57 | 3854511 | 385432 | 4.239943 | 2.10077 | 1.298928 | 0.896019 | 0.64874 | 0.542697 |
| 556 | 24.03 | 3781002 | 379775 | 4.160777 | 2.101626 | 1.254001 | 0.895576 | 0.635404 | 0.538869 |
| 557 | 24.06 | 3846542 | 385067 | 4.231609 | 2.105981 | 1.255402 | 0.887823 | 0.643221 | 0.545515 |
| 558 | 26.54 | 3803647 | 381289 | 4.184936 | 2.110767 | 1.20577 | 0.890883 | 0.633439 | 0.540937 |
| 559 | 23.18 | 3843252 | 384410 | 4.227662 | 2.110204 | 1.249361 | 0.887667 | 0.643555 | 0.542591 |
| 560 | 24.12 | 3790983 | 380042 | 4.171025 | 2.108526 | 1.215334 | 0.889465 | 0.627849 | 0.544148 |
| 561 | 26.93 | 3821394 | 384001 | 4.205395 | 2.107658 | 1.210161 | 0.892801 | 0.631184 | 0.541711 |
| 562 | 32.94 | 3848628 | 386667 | 4.235295 | 2.102583 | 1.206539 | 0.886982 | 0.638416 | 0.542396 |
| 563 | 29.79 | 3835231 | 384125 | 4.219356 | 2.107384 | 1.20587 | 0.891102 | 0.633974 | 0.540726 |
| 564 | 25.86 | 3910750 | 394413 | 4.305163 | 2.10601 | 1.247344 | 0.892693 | 0.640235 | 0.540474 |
| 565 | 26.25 | 3904871 | 392755 | 4.297626 | 2.105643 | 1.233533 | 0.892223 | 0.648475 | 0.536367 |
| 566 | 29.9 | 4481349 | 458166 | 4.939515 | 2.104717 | 1.273546 | 0.900365 | 0.698483 | 0.536503 |
| 567 | 27.28 | 3868075 | 388503 | 4.256578 | 2.105942 | 1.221261 | 0.888441 | 0.625881 | 0.538438 |
| 568 | 23.09 | 3927390 | 393885 | 4.321275 | 2.108249 | 1.246773 | 0.892455 | 0.642471 | 0.535475 |
| 569 | 33.36 | 3882393 | 387339 | 4.269732 | 2.10757 | 1.238163 | 0.893834 | 0.646267 | 0.541069 |
| 570 | 23.41 | 3859167 | 386966 | 4.246133 | 2.109229 | 1.251372 | 0.89511 | 0.633816 | 0.538878 |
| 571 | 23.01 | 3890722 | 390868 | 4.28159 | 2.105344 | 1.243896 | 0.892582 | 0.639692 | 0.542237 |
| 574 | 35.09 | 3825306 | 384523 | 4.209829 | 2.105436 | 1.204961 | 0.892018 | 0.649255 | 0.54006 |
| 576 | 26.78 | 3877799 | 387132 | 4.264931 | 2.11033 | 1.223344 | 0.895579 | 0.649503 | 0.537401 |
| 578 | 27.59 | 3857943 | 390520 | 4.248463 | 2.109807 | 1.241243 | 0.891085 | 0.651591 | 0.539558 |
| 581 | 32.45 | 3842893 | 385223 | 4.228116 | 2.110049 | 1.232327 | 0.89504 | 0.64782 | 0.538189 |
| 582 | 22.6 | 3875823 | 389128 | 4.264951 | 2.105326 | 1.234633 | 0.893575 | 0.638391 | 0.537623 |
| 586 | 25.58 | 3910729 | 391649 | 4.302378 | 2.106296 | 1.225464 | 0.890934 | 0.636839 | 0.540722 |
| 591 | 42.27 | 3854850 | 385403 | 4.240253 | 2.102411 | 1.189051 | 0.887279 | 0.620887 | 0.543314 |
| 592 | 23.64 | 3771960 | 377880 | 4.14984 | 2.107221 | 1.200805 | 0.891639 | 0.63171 | 0.538138 |
| 593 | 29.8 | 3809639 | 382064 | 4.191703 | 2.104736 | 1.189447 | 0.889219 | 0.638838 | 0.542429 |
| 596 | 26.07 | 3812393 | 381623 | 4.194016 | 2.107019 | 1.236356 | 0.890694 | 0.638514 | 0.538522 |
| 598 | 25.2 | 3912782 | 391583 | 4.304365 | 2.10516 | 1.277945 | 0.895017 | 0.646107 | 0.543212 |
| 599 | 29.29 | 3833225 | 386820 | 4.220045 | 2.106146 | 1.210429 | 0.889112 | 0.648225 | 0.543433 |
| 601 | 26.92 | 3819953 | 380441 | 4.200394 | 2.10799 | 1.230056 | 0.892585 | 0.642124 | 0.541883 |
| 603 | 30.27 | 3831902 | 382382 | 4.214284 | 2.106339 | 1.215151 | 0.894639 | 0.646612 | 0.539815 |
| 605 | 29.06 | 4510934 | 458461 | 4.969395 | 2.108411 | 1.285913 | 0.899328 | 0.687617 | 0.538074 |
| 607 | 26.73 | 3916564 | 390319 | 4.306883 | 2.108225 | 1.234554 | 0.891664 | 0.647615 | 0.535882 |
| 612 | 26.34 | 3784560 | 377544 | 4.162104 | 2.107633 | 1.217379 | 0.892541 | 0.639654 | 0.538223 |
| 613 | 28.5 | 3825515 | 385245 | 4.21076 | 2.108854 | 1.206779 | 0.890842 | 0.645852 | 0.541725 |
| 614 | 29.25 | 3856346 | 384836 | 4.241182 | 2.107023 | 1.211555 | 0.8931 | 0.654982 | 0.539638 |
| 615 | 28.35 | 3856744 | 388331 | 4.245075 | 2.107714 | 1.221741 | 0.89356 | 0.646838 | 0.543247 |
| 618 | 26.75 | 3839784 | 383804 | 4.223588 | 2.108727 | 1.216159 | 0.893379 | 0.644285 | 0.540514 |
| 619 | 26.18 | 4390841 | 442477 | 4.833318 | 2.109203 | 1.307434 | 0.904122 | 0.673301 | 0.534724 |
| 621 | 29.24 | 3812250 | 380176 | 4.192426 | 2.104332 | 1.19544 | 0.889541 | 0.639876 | 0.540797 |
| 622 | 44.08 | 3853677 | 385667 | 4.239344 | 2.106287 | 1.17681 | 0.892597 | 0.628101 | 0.53944 |
| 624 | 25.2 | 3913293 | 391243 | 4.304536 | 2.105206 | 1.23332 | 0.892238 | 0.647348 | 0.536931 |
| 625 | 32.01 | 4209477 | 422397 | 4.631874 | 2.107116 | 1.238076 | 0.891726 | 0.671096 | 0.534952 |
| 628 | 27.61 | 3824518 | 381317 | 4.205835 | 2.10537 | 1.199179 | 0.889969 | 0.643713 | 0.540393 |
| 629 | 27.53 | 3784588 | 377565 | 4.162153 | 2.108907 | 1.191896 | 0.89276 | 0.642813 | 0.541017 |
| 632 | 29.63 | 3770296 | 375701 | 4.145997 | 2.106126 | 1.201898 | 0.892774 | 0.650098 | 0.542592 |
| 634 | 29.12 | 3887804 | 387370 | 4.275174 | 2.107587 | 1.22666 | 0.895854 | 0.641691 | 0.5401 |

| | | | | | | | | | |
|---|---|---|---|---|---|---|---|---|---|
| 638 | 30.71 | 3835297 | 386434 | 4.221731 | 2.103132 | 1.24516 | 0.894849 | 0.629635 | 0.54072 |
| 643 | 26.08 | 3817469 | 381806 | 4.199275 | 2.10542 | 1.247206 | 0.895654 | 0.652053 | 0.544903 |
| 645 | 26.98 | 3834538 | 379705 | 4.214243 | 2.107282 | 1.228562 | 0.897472 | 0.649825 | 0.539896 |
| 649 | 26.23 | 3828762 | 381739 | 4.210501 | 2.110279 | 1.231667 | 0.89537 | 0.632459 | 0.539301 |
| 650 | 26.46 | 3833779 | 381343 | 4.215122 | 2.105403 | 1.22131 | 0.895253 | 0.635385 | 0.541663 |
| 653 | 26.7 | 3793193 | 378458 | 4.171651 | 2.109115 | 1.186682 | 0.894606 | 0.649128 | 0.540083 |
| 656 | 27.43 | 3843135 | 382379 | 4.225514 | 2.105129 | 1.188062 | 0.893282 | 0.625676 | 0.539161 |
| 657 | 40.27 | 3881205 | 384262 | 4.265467 | 2.107609 | 1.212607 | 0.895591 | 0.631509 | 0.541441 |
| 658 | 27.28 | 3836038 | 381958 | 4.217996 | 2.111441 | 1.199604 | 0.891239 | 0.645597 | 0.5383 |
| 659 | 32.03 | 4660964 | 477322 | 5.138286 | 2.108337 | 1.286658 | 0.896534 | 0.692436 | 0.535334 |
| 662 | 30.22 | 3804626 | 380418 | 4.185044 | 2.107186 | 1.206357 | 0.894023 | 0.636877 | 0.540741 |
| 668 | 27.79 | 3909008 | 389544 | 4.298552 | 2.108544 | 1.212731 | 0.894985 | 0.64117 | 0.540493 |
| 673 | 25.96 | 3902109 | 389966 | 4.292075 | 2.107548 | 1.231599 | 0.891815 | 0.642479 | 0.540487 |
| 676 | 32.43 | 3960817 | 395098 | 4.355915 | 2.104344 | 1.238416 | 0.893655 | 0.648337 | 0.541656 |
| 677 | 25.53 | 3793272 | 378370 | 4.171642 | 2.108459 | 1.211237 | 0.894156 | 0.64019 | 0.540349 |
| 684 | 27.57 | 3806731 | 380392 | 4.187123 | 2.102967 | 1.23109 | 0.895332 | 0.645529 | 0.542447 |
| 686 | 34.61 | 3871440 | 386493 | 4.257933 | 2.108428 | 1.203141 | 0.890061 | 0.645413 | 0.539228 |
| 693 | 31.75 | 3817961 | 385251 | 4.203212 | 2.105214 | 1.206914 | 0.891205 | 0.645446 | 0.541408 |
| 697 | 27.41 | 3802475 | 379793 | 4.182268 | 2.108186 | 1.216027 | 0.894134 | 0.64037 | 0.542827 |
| 709 | 25.88 | 3829984 | 381249 | 4.211233 | 2.106866 | 1.211807 | 0.892542 | 0.650972 | 0.540667 |
| 710 | 25.68 | 3794209 | 378589 | 4.172798 | 2.103474 | 1.245024 | 0.898944 | 0.643411 | 0.544585 |
| 711 | 29.43 | 3822856 | 382482 | 4.205338 | 2.106544 | 1.219798 | 0.887774 | 0.633657 | 0.54233 |
| 712 | 29.68 | 3839701 | 383224 | 4.222925 | 2.106655 | 1.206222 | 0.891059 | 0.632786 | 0.544616 |
| 715 | 25.72 | 3820932 | 379077 | 4.200009 | 2.109648 | 1.22381 | 0.896945 | 0.632976 | 0.53785 |
| 721 | 30.44 | 3802696 | 379282 | 4.181978 | 2.105902 | 1.219929 | 0.889961 | 0.636922 | 0.538213 |
| 728 | 26.28 | 3808330 | 378885 | 4.187215 | 2.105303 | 1.200984 | 0.89187 | 0.635124 | 0.540561 |
| 731 | 29.59 | 3814465 | 380187 | 4.194652 | 2.105844 | 1.193265 | 0.892704 | 0.640149 | 0.543361 |
| 732 | 32.93 | 3821945 | 382717 | 4.204662 | 2.103794 | 1.202808 | 0.890998 | 0.63874 | 0.541675 |
| 733 | 28.91 | 3822661 | 385138 | 4.207799 | 2.107165 | 1.203422 | 0.89008 | 0.648408 | 0.541821 |
| 734 | 26.79 | 3823773 | 385917 | 4.20969 | 2.107886 | 1.193524 | 0.890846 | 0.648085 | 0.541399 |
| 735 | 29.26 | 3787080 | 376863 | 4.163943 | 2.104602 | 1.18223 | 0.893065 | 0.643538 | 0.539439 |
| 738 | 26.47 | 3833706 | 381804 | 4.21551 | 2.105333 | 1.21321 | 0.893067 | 0.633539 | 0.542162 |
| 741 | 26.52 | 3773213 | 376201 | 4.149414 | 2.108472 | 1.202642 | 0.89286 | 0.645234 | 0.538942 |
| 751 | 26.93 | 3832555 | 380915 | 4.21347 | 2.105334 | 1.221458 | 0.891894 | 0.637425 | 0.540528 |
| 754 | 25.86 | 3777257 | 376101 | 4.153358 | 2.107428 | 1.186297 | 0.89243 | 0.632284 | 0.539684 |
| 766 | 26 | 3810414 | 378437 | 4.188851 | 2.108048 | 1.215452 | 0.890563 | 0.636882 | 0.545412 |
| 769 | 25.36 | 3850884 | 384403 | 4.235287 | 2.110066 | 1.217092 | 0.896717 | 0.634842 | 0.539603 |
| 772 | 27.12 | 3894285 | 390667 | 4.284952 | 2.108776 | 1.240082 | 0.891155 | 0.652184 | 0.539295 |
| 780 | 26.87 | 4355411 | 439259 | 4.79467 | 2.108996 | 1.25982 | 0.89713 | 0.689374 | 0.535491 |
| 783 | 28.96 | 3854184 | 385507 | 4.239691 | 2.109019 | 1.19946 | 0.891605 | 0.663109 | 0.54181 |
| 787 | 28.99 | 4565749 | 464826 | 5.030575 | 2.106888 | 1.274097 | 0.898159 | 0.688595 | 0.536379 |
| 788 | 28.75 | 3829166 | 383719 | 4.212885 | 2.108684 | 1.209587 | 0.898888 | 0.660521 | 0.53919 |
| 793 | 34.01 | 3825620 | 383923 | 4.209543 | 2.105314 | 1.201893 | 0.890046 | 0.663957 | 0.542139 |
| 794 | 27.43 | 3827898 | 380366 | 4.208264 | 2.106177 | 1.228077 | 0.889475 | 0.651323 | 0.540138 |
| 799 | 28.59 | 3811031 | 382141 | 4.193172 | 2.107031 | 1.202882 | 0.894151 | 0.668527 | 0.543686 |
| 803 | 29.5 | 3835116 | 383934 | 4.21905 | 2.105679 | 1.203638 | 0.895445 | 0.655018 | 0.539636 |
| 808 | 32.37 | 3853406 | 384938 | 4.238344 | 2.107079 | 1.199046 | 0.892463 | 0.648097 | 0.545236 |
| 811 | 26.86 | 3808075 | 379845 | 4.18792 | 2.106194 | 1.209609 | 0.890556 | 0.642831 | 0.544597 |
| 813 | 27.72 | 3824873 | 380710 | 4.205583 | 2.109501 | 1.210834 | 0.896081 | 0.633994 | 0.538786 |
| 814 | 26.12 | 3800165 | 384741 | 4.184906 | 2.103301 | 1.203481 | 0.886889 | 0.636498 | 0.54112 |
| 815 | 25.68 | 3772464 | 374962 | 4.147426 | 2.108308 | 1.205864 | 0.8938 | 0.642816 | 0.538434 |
| 817 | 25.74 | 3868590 | 385653 | 4.254243 | 2.109776 | 1.223976 | 0.892605 | 0.646127 | 0.540561 |
| 820 | 25.4 | 3837481 | 382680 | 4.220161 | 2.109411 | 1.25101 | 0.892965 | 0.667541 | 0.546537 |
| 821 | 29.58 | 4533284 | 461500 | 4.994784 | 2.108013 | 1.286832 | 0.896515 | 0.683479 | 0.533349 |
| 822 | 28.03 | 3843263 | 382839 | 4.226102 | 2.106321 | 1.240383 | 0.891284 | 0.669077 | 0.543836 |
| 823 | 23.94 | 3812004 | 385327 | 4.197331 | 2.103924 | 1.215284 | 0.892494 | 0.655282 | 0.542583 |
| 827 | 31.28 | 3843880 | 387840 | 4.23172 | 2.109215 | 1.206206 | 0.891692 | 0.636188 | 0.544738 |
| 839 | 25.25 | 3883640 | 393393 | 4.277033 | 2.105147 | 1.232287 | 0.890112 | 0.642709 | 0.536011 |
| 844 | 26.65 | 4435311 | 454940 | 4.890251 | 2.109754 | 1.271354 | 0.897051 | 0.690763 | 0.533552 |
| 851 | 27.56 | 3826600 | 382877 | 4.209477 | 2.107953 | 1.217555 | 0.889127 | 0.635102 | 0.54404 |
| 852 | 27.8 | 3939680 | 398700 | 4.33838 | 2.109442 | 1.208303 | 0.891454 | 0.647631 | 0.537364 |
| 854 | 24.36 | 3938294 | 395025 | 4.333319 | 2.10731 | 1.271783 | 0.893031 | 0.631582 | 0.536866 |
| 855 | 24.53 | 3855180 | 385096 | 4.240276 | 2.10752 | 1.177986 | 0.88915 | 0.64307 | 0.530569 |
| 856 | 24.63 | 3929270 | 397513 | 4.326783 | 2.104165 | 1.247947 | 0.89601 | 0.645249 | 0.535176 |
| 857 | 28.51 | 3825906 | 387491 | 4.213397 | 2.101339 | 1.227999 | 0.894502 | 0.654233 | 0.541927 |
| 860 | 28.51 | 3829289 | 388095 | 4.217384 | 2.10102 | 1.238526 | 0.894055 | 0.653761 | 0.53627 |
| 861 | 26.2 | 3881156 | 392537 | 4.273693 | 2.10956 | 1.230353 | 0.891503 | 0.650129 | 0.543321 |
| 862 | 31.37 | 3804286 | 385087 | 4.189373 | 2.107746 | 1.19358 | 0.889284 | 0.640538 | 0.539729 |
| 868 | 27.71 | 3860065 | 390358 | 4.250423 | 2.108272 | 1.227886 | 0.892655 | 0.648479 | 0.539664 |
| 869 | 28.71 | 3829284 | 382926 | 4.21221 | 2.105147 | 1.207921 | 0.889173 | 0.637708 | 0.544986 |

| | | | | | | | | | |
|---|---|---|---|---|---|---|---|---|---|
| 875 | 26.34 | 3827458 | 387209 | 4.214667 | 2.109535 | 1.196381 | 0.889167 | 0.633637 | 0.541586 |
| 885 | 31.94 | 3830412 | 383818 | 4.21423 | 2.108475 | 1.205188 | 0.89346 | 0.659092 | 0.539889 |
| 886 | 30.98 | 3927483 | 393653 | 4.321136 | 2.107541 | 1.224575 | 0.895532 | 0.665896 | 0.537371 |
| 887 | 32.22 | 3827302 | 386977 | 4.214279 | 2.107049 | 1.202784 | 0.890489 | 0.651258 | 0.540356 |
| 888 | 29.75 | 3875682 | 392388 | 4.26807 | 2.101976 | 1.227327 | 0.895601 | 0.649028 | 0.537535 |
| 910 | 27.35 | 3799459 | 383497 | 4.182956 | 2.108686 | 1.205253 | 0.887732 | 0.648289 | 0.537793 |
| 912 | 26.59 | 3793392 | 383325 | 4.176717 | 2.104856 | 1.20212 | 0.8902 | 0.639903 | 0.542617 |
| 914 | 25.89 | 4529520 | 463926 | 4.993446 | 2.110399 | 1.269314 | 0.900788 | 0.698838 | 0.530748 |
| 915 | 30.67 | 3821838 | 385273 | 4.207111 | 2.105823 | 1.205663 | 0.889687 | 0.644363 | 0.542042 |
| 916 | 31.32 | 3879382 | 392947 | 4.272329 | 2.102128 | 1.214595 | 0.893824 | 0.630561 | 0.540059 |
| 917 | 29.41 | 3826866 | 386865 | 4.213731 | 2.108888 | 1.210192 | 0.890677 | 0.643044 | 0.544117 |
| 919 | 25.31 | 3816653 | 386067 | 4.20272 | 2.100337 | 1.248674 | 0.895572 | 0.650343 | 0.544646 |
| 920 | 35.83 | 3819082 | 382962 | 4.202044 | 2.103858 | 1.196014 | 0.892262 | 0.632934 | 0.538295 |
| 923 | 26.63 | 4594787 | 469361 | 5.064148 | 2.106969 | 1.293591 | 0.898675 | 0.686058 | 0.533193 |
| 925 | 29.29 | 3817545 | 383783 | 4.201328 | 2.103195 | 1.226656 | 0.896028 | 0.638915 | 0.542062 |
| 927 | 28.3 | 4435269 | 451680 | 4.886949 | 2.108922 | 1.26457 | 0.895766 | 0.68591 | 0.529563 |
| 928 | 31.72 | 3898933 | 391993 | 4.290926 | 2.104267 | 1.23861 | 0.892991 | 0.649254 | 0.538731 |
| 930 | 26.01 | 4594498 | 469505 | 5.064003 | 2.10998 | 1.292553 | 0.897416 | 0.693991 | 0.531665 |
| 931 | 24.51 | 3867248 | 386752 | 4.254 | 2.105355 | 1.236173 | 0.890057 | 0.64164 | 0.539414 |
| 932 | 28.56 | 3844962 | 386334 | 4.231296 | 2.106904 | 1.215651 | 0.889525 | 0.651249 | 0.537434 |
| 933 | 23.66 | 3913740 | 392235 | 4.305975 | 2.107384 | 1.265489 | 0.895362 | 0.635799 | 0.53638 |
| 934 | 24.31 | 3868710 | 388950 | 4.25766 | 2.106026 | 1.290452 | 0.893838 | 0.655312 | 0.541492 |
| 935 | 28.72 | 3760777 | 375496 | 4.136273 | 2.106981 | 1.264217 | 0.894634 | 0.64246 | 0.538148 |
| 936 | 27.44 | 4494471 | 456735 | 4.951206 | 2.108193 | 1.27969 | 0.90088 | 0.689716 | 0.530343 |
| 937 | 28.98 | 3795086 | 381887 | 4.176973 | 2.107486 | 1.198482 | 0.889781 | 0.632029 | 0.538819 |
| 938 | 25.3 | 3788614 | 379569 | 4.168183 | 2.105821 | 1.220828 | 0.889541 | 0.642086 | 0.536965 |
| 941 | 31.9 | 3942853 | 395964 | 4.338817 | 2.105176 | 1.238997 | 0.894315 | 0.640987 | 0.537277 |
| 942 | 31.91 | 3902680 | 393493 | 4.296173 | 2.102762 | 1.225319 | 0.888332 | 0.640989 | 0.538704 |
| 944 | 30.12 | 3836670 | 385791 | 4.222461 | 2.10155 | 1.239069 | 0.896669 | 0.629867 | 0.538951 |
| 946 | 29 | 3871471 | 389723 | 4.261194 | 2.105068 | 1.267692 | 0.894257 | 0.646991 | 0.532008 |
| 950 | 32.15 | 4550262 | 464423 | 5.014685 | 2.106607 | 1.264858 | 0.899475 | 0.689228 | 0.533507 |
| 954 | 25.16 | 3807769 | 383015 | 4.190784 | 2.102697 | 1.286218 | 0.8993 | 0.632907 | 0.548368 |
| 955 | 31.45 | 3852081 | 386036 | 4.238117 | 2.106851 | 1.193252 | 0.889265 | 0.630958 | 0.539979 |
| 957 | 23.66 | 3887583 | 390174 | 4.277757 | 2.108257 | 1.250352 | 0.890767 | 0.644916 | 0.538453 |
| 959 | 31.42 | 3913660 | 392376 | 4.306036 | 2.103446 | 1.223602 | 0.892265 | 0.641814 | 0.539481 |
| 960 | 25.03 | 3843149 | 384778 | 4.227927 | 2.10832 | 1.24239 | 0.892722 | 0.652274 | 0.536566 |
| 961 | 28.44 | 3831715 | 383906 | 4.215621 | 2.101104 | 1.226139 | 0.893285 | 0.636445 | 0.541845 |
| 962 | 24.74 | 3819767 | 383234 | 4.203001 | 2.104028 | 1.235808 | 0.893866 | 0.643692 | 0.538925 |
| 963 | 26.9 | 4333990 | 437751 | 4.771741 | 2.107381 | 1.271744 | 0.89237 | 0.674357 | 0.534742 |
| 964 | 24.07 | 4075803 | 408375 | 4.484178 | 2.105699 | 1.277153 | 0.890698 | 0.658857 | 0.535069 |
| 965 | 24.88 | 3825490 | 381984 | 4.207474 | 2.107167 | 1.222251 | 0.89105 | 0.639069 | 0.537589 |
| 967 | 24.91 | 3813135 | 381431 | 4.194566 | 2.101481 | 1.259115 | 0.897784 | 0.640016 | 0.538754 |
| 969 | 26.71 | 4636812 | 472025 | 5.108837 | 2.107606 | 1.306827 | 0.899555 | 0.694704 | 0.533238 |
| 974 | 24.08 | 3897968 | 390365 | 4.288333 | 2.106943 | 1.261719 | 0.895142 | 0.647626 | 0.537005 |
| 979 | 24.23 | 4061999 | 407474 | 4.469473 | 2.103407 | 1.260202 | 0.882023 | 0.633614 | 0.538196 |
| 986 | 23.99 | 3801930 | 380341 | 4.182271 | 2.109952 | 1.233919 | 0.890487 | 0.632106 | 0.53478 |
| 989 | 31.62 | 3847243 | 385767 | 4.23301 | 2.10491 | 1.244298 | 0.892531 | 0.636492 | 0.539322 |
| 990 | 27.09 | 3800286 | 380381 | 4.180667 | 2.101255 | 1.24739 | 0.894367 | 0.636498 | 0.543477 |
| 991 | 29.34 | 3826294 | 383741 | 4.210035 | 2.09948 | 1.244451 | 0.893114 | 0.643211 | 0.542404 |
| 995 | 29.07 | 3792613 | 379800 | 4.172413 | 2.107034 | 1.228796 | 0.891907 | 0.637686 | 0.538604 |
| 1074 | 33.63 | 3827240 | 386022 | 4.213262 | 2.10393 | 1.188613 | 0.889481 | 0.639597 | 0.541014 |
| 2002 | 18.15 | 3768940 | 375476 | 4.144416 | 2.107516 | 1.245797 | 0.888442 | 0.64507 | 0.535658 |
| 2003 | 27.18 | 3840875 | 385585 | 4.22646 | 2.103674 | 1.276914 | 0.891841 | 0.633773 | 0.540628 |
| 2006 | 27.48 | 3821112 | 382408 | 4.20352 | 2.107558 | 1.232536 | 0.89056 | 0.637035 | 0.53904 |
| 2007 | 25.75 | 3772621 | 377563 | 4.150184 | 2.107726 | 1.21746 | 0.890069 | 0.642528 | 0.538732 |
| 2009 | 25.67 | 3868526 | 387946 | 4.256472 | 2.106341 | 1.26533 | 0.893326 | 0.639467 | 0.537393 |
| 2012 | 27.03 | 3865646 | 386435 | 4.252081 | 2.106597 | 1.239821 | 0.890128 | 0.628021 | 0.539849 |
| 2013 | 30.01 | 3875716 | 387676 | 4.263392 | 2.105278 | 1.225523 | 0.8909 | 0.640775 | 0.535001 |
| 2015 | 25.16 | 3954241 | 396506 | 4.350747 | 2.107726 | 1.271803 | 0.891587 | 0.643197 | 0.537859 |
| 2020 | 25.46 | 3804341 | 379368 | 4.183709 | 2.108446 | 1.22486 | 0.889306 | 0.632436 | 0.540296 |
| 2025 | 28.89 | 3788214 | 379769 | 4.167983 | 2.108456 | 1.208386 | 0.89146 | 0.638097 | 0.536575 |
| 2032 | 25.75 | 3808157 | 383192 | 4.191349 | 2.10413 | 1.242658 | 0.896046 | 0.63512 | 0.535662 |
| 2035 | 23.15 | 3778218 | 379144 | 4.157362 | 2.102358 | 1.244352 | 0.895803 | 0.635294 | 0.540638 |
| 2036 | 21.22 | 3768710 | 375440 | 4.14415 | 2.10814 | 1.235207 | 0.890786 | 0.624195 | 0.536605 |
| 2038 | 24.48 | 3835087 | 384067 | 4.219154 | 2.108017 | 1.25847 | 0.889718 | 0.642276 | 0.535311 |
| 2039 | 21.36 | 3835354 | 383360 | 4.218714 | 2.106013 | 1.281718 | 0.893383 | 0.641826 | 0.536325 |
| 2051 | 23.12 | 3789540 | 379165 | 4.168705 | 2.103565 | 1.25458 | 0.895512 | 0.624804 | 0.537538 |
| 2055 | 27.36 | 3767378 | 376810 | 4.144188 | 2.106596 | 1.231305 | 0.890295 | 0.642811 | 0.533947 |
| 2059 | 28.06 | 3854475 | 385739 | 4.240214 | 2.107919 | 1.245391 | 0.893225 | 0.638775 | 0.538721 |
| 2061 | 28.27 | 3806904 | 382543 | 4.189447 | 2.10208 | 1.242881 | 0.895462 | 0.639939 | 0.53584 |

| | | | | | | | | | |
|---|---|---|---|---|---|---|---|---|---|
| 2062 | 31.65 | 3890294 | 389226 | 4.27952 | 2.103634 | 1.250413 | 0.892529 | 0.636379 | 0.538738 |
| 2063 | 25.27 | 3878027 | 387026 | 4.265053 | 2.103857 | 1.255942 | 0.892539 | 0.629154 | 0.536371 |
| 2064 | 24.18 | 4579893 | 465633 | 5.045526 | 2.11001 | 1.324896 | 0.901485 | 0.706886 | 0.533794 |
| 2065 | 27.12 | 3789703 | 379182 | 4.168885 | 2.105612 | 1.261827 | 0.896477 | 0.640076 | 0.536299 |
| 2073 | 26.23 | 3774070 | 376594 | 4.150664 | 2.10134 | 1.233172 | 0.89075 | 0.640496 | 0.53985 |
| 2086 | 26.08 | 3798956 | 381851 | 4.180807 | 2.104584 | 1.221077 | 0.889724 | 0.643531 | 0.540344 |
| 2136 | 32.45 | 3824152 | 384411 | 4.208563 | 2.105538 | 1.19925 | 0.889623 | 0.639935 | 0.543434 |
| 2137 | 29.61 | 3786308 | 380457 | 4.166765 | 2.104759 | 1.21047 | 0.892229 | 0.642239 | 0.540445 |
| 2138 | 30.25 | 3833546 | 385815 | 4.219361 | 2.105622 | 1.230349 | 0.888805 | 0.648133 | 0.540289 |
| 2156 | 29.51 | 3835306 | 385803 | 4.221109 | 2.103131 | 1.244325 | 0.892857 | 0.64969 | 0.542338 |
| 2157 | 27.05 | 3819272 | 384715 | 4.203987 | 2.103261 | 1.244555 | 0.8904 | 0.634469 | 0.541714 |
| 2159 | 33.26 | 3845646 | 386501 | 4.232147 | 2.107949 | 1.206342 | 0.892064 | 0.644201 | 0.5394 |
| 2161 | 30.1 | 3823240 | 385619 | 4.208859 | 2.101917 | 1.238887 | 0.893712 | 0.64875 | 0.545441 |
| 2162 | 28.1 | 3825922 | 385551 | 4.211473 | 2.103573 | 1.237383 | 0.893807 | 0.642141 | 0.541395 |
| 2163 | 29.62 | 3843054 | 387093 | 4.230147 | 2.102129 | 1.241833 | 0.895986 | 0.638045 | 0.541262 |
| 2167 | 35.75 | 3831137 | 385839 | 4.216976 | 2.106951 | 1.218269 | 0.894462 | 0.636788 | 0.544814 |
| 2184 | 31.54 | 3812631 | 382933 | 4.195564 | 2.10654 | 1.214868 | 0.890871 | 0.641188 | 0.53965 |
| 2186 | 27.57 | 3946744 | 397822 | 4.344566 | 2.105613 | 1.249412 | 0.893405 | 0.649139 | 0.538511 |
| 2189 | 30.6 | 3833674 | 386746 | 4.22042 | 2.103503 | 1.225446 | 0.894576 | 0.643086 | 0.541181 |
| 2194 | 33.05 | 3841762 | 388162 | 4.229924 | 2.102078 | 1.230876 | 0.894781 | 0.635206 | 0.542586 |
| 2197 | 33.15 | 3821946 | 386378 | 4.208324 | 2.101011 | 1.225057 | 0.892394 | 0.647141 | 0.540498 |

**Table S4**

| Gene | ACMG - HCM/DCM/LVNC | HGMD ("LVNC" 10/02/19 & "left ventricular noncompaction" 10/02/19) | PMC5837460 - HCM | PMC5116235 - HCM | HGMD ("HCM" 2/6/19) | Fulgent HCM | Centogene HCM | Mayo_inheritence_disease HCM | GeneDx HCM | Invitae HCM | PMC5116235 - DCM | HGMD ("DCM" 2/11/19) | Invitae DCM | Mayo_inheritence_disease DCM |
|---|---|---|---|---|---|---|---|---|---|---|---|---|---|---|
| A2ML1 | | | | | | fulgent | | | | Invitae-RASopathy | | | Invitae-RASopathy | |
| ABCC9 | | | | | (cardiomyopathy, hypertrophic) | fulgent | | | GeneDx | | no excess | (Cardiomyopathy, dilated) | Invitae-primary panel | mayo_AD_DCM, Cantu syndrome |
| ACAD9 | | | | | (cardiomyopathy, hypertrophic) | | | | | | | | | |
| ACADVL | | | | | | fulgent | | | | Invitae-AR syndromic pediatric cardiomyopathy | | | Invitae-AR syndromic pediatric cardiomyopathy | |
| ACE | | | | | (Hypertrophic cardiomyopathy, association with) | | | | | | | | | |
| ACTA1 | | | rare, recessive, phenocopy | | (Hypertrophic cardiomyopathy, myocardial noncompaction & transmural crypts); (cardiomyopathy, hypertrophic); (Nemaline myopathy and hypertrophic cardiomyopathy) | | | | | | | (Fibre type disproportion, congenital and dilated cardiomyopathy);(Nemaline myopathy and dilated cardiomyopathy); (Cardiomyopathy, dilated) | | |
| ACTC1 | (HCM, test DCM/LVNC); (secondary finding) | (Cardiomyopathy, hypertrophic/dilated with left ventricular noncompaction);(Left ventricular noncompaction with arrhythmias) | sarcomere, validated | Nontruncating excess | (cardiomyopathy, hypertrophic) | fulgent | centogene | mayo_AD_CHD, DCM, HCM, LVNC | GeneDx | Invitae-primary panel | no excess | (Cardiomyopathy, dilated) | Invitae-primary panel | mayo_AD_CHD, DCM, HCM, LVNC |
| ACTN2 | (HCM, test DCM/LVNC) | (Ventricular fibrillation, left ventricular noncompaction and sudden death) | moderate: no excess but human evidence -> very rare that mutation in this gene is causitive, most mutations benign | no excess | (cardiomyopathy, hypertrophic) | fulgent | centogene | mayo_AD_DCM, HCM | GeneDx | Invitae-primary panel | no excess | (Cardiomyopathy, dilated) | Invitae-primary panel | mayo_AD_DCM, HCM |
| ADRB2 | | | | | | | | | | | | (Cardiomyopathy, dilated) | | |
| AGL | | | | | | fulgent | | | | Invitae-primary panel | | | Invitae-primary panel | |
| AGTR1 | | | | | (Hypertrophic cardiomyopathy, association with) | | | | | | | | | |
| AKAP9 | | | | | (cardiomyopathy, hypertrophic) | | | | GeneDx | | | (Cardiomyopathy, dilated) | | |
| ALMS1 | | | | | | | | | GeneDx | | | | | |
| ALPK3 | | | | | (cardiomyopathy, hypertrophic) | | | | GeneDx | | | | | |
| ANKRD1 | | | functional data only | no excess | (cardiomyopathy, hypertrophic) | fulgent | centogene | mayo_AD_HCM, DCM | GeneDx | Invitae-preliminary evidence | no excess | (Dilated cardiomyopathy); (Cardiomyopathy, dilated) | Invitae-preliminary evidence | mayo_AD_HCM, DCM |
| B2M | | | | | (cardiomyopathy, hypertrophic) | | | | | | | | | |
| BAG3 | (DCM, test LVNC) | | | | (cardiomyopathy, hypertrophic) | fulgent | | | GeneDx | Invitae-primary panel | | (Cardiomyopathy, dilated) | Invitae-primary panel | |
| BRAF | | | | | (cardiomyopathy, hypertrophic) | fulgent | | | GeneDx | Invitae-RASopathy | | | Invitae-RASopathy | |
| C2orf40 | | | | | | | | | | | | (Cardiomyopathy, dilated) | | |
| CACNA1C | | | HCM+arrythmia -> likely cause arrythmia only | | (cardiomyopathy, hypertrophic); (Long QT syndrome with hypertrophic cardiomyopathy) | fulgent | | | | Invitae-primary panel | | | Invitae-primary panel | |
| CALM2 | | | | | (Long QT syndrome & hypertrophic cardiomyopathy) | | | | | | | | | |
| CALR3 | | | no evidence (same rare variant frequency as EXAC) | | | fulgent | centogene | | | Invitae-preliminary evidence | | (Cardiomyopathy, dilated) | Invitae-preliminary evidence | |
| CASK | | | | | | | | | | | | (Developmental delay, hearing loss and dilated cardiomyopathy) | | |
| CASQ2 | | | no evidence (same rare variant frequency as EXAC) | | (cardiomyopathy, hypertrophic) | | | | | | no excess | (Cardiomyopathy, dilated) | | |
| CASZ1 | | | | | | | | | | | | (Cardiomyopathy, dilated) | | |
| CAV3 | | | functional data only | | (cardiomyopathy, hypertrophic); (Hypertrophic cardiomyopathy) | fulgent | centogene | mayo_AD, AR_HCM, LQTS, LGMD, Tateyama-type distal myopathy, rippling muscle disease | GeneDx | Invitae-primary panel | no excess | (Dilated cardiomyopathy and limb girdle muscular dystrophy 1C) | Invitae-primary panel | |
| CAVIN4 | | | | | | | | | | | | (Cardiomyopathy, dilated) | | |
| CBL | | | | | (cardiomyopathy, hypertrophic) | fulgent | | | | Invitae-RASopathy | | | Invitae-RASopathy | |
| cflartv7 | | | | | (cardiomyopathy, hypertrophic) | | | | | | | | | |
| CHRM2 | | | | | | | | | GeneDx | | | (Cardiomyopathy, dilated) | Invitae-preliminary evidence | |
| CMYA5 | | | | | (cardiomyopathy, hypertrophic) | | | | | | | | | |
| CNBP | | | | | | | | | | | | (Cardiomyopathy, dilated) | | |
| COA5 | | | | | (Hypertrophic cardiomyopathy, fatal neonatal) | | | | | | | | | |
| COX15 | | | rare, recessive, phenocopy | | (Hypertrophic cardiomyopathy, early onset) | | | | | | | | | |
| CPT2 | | | | | | fulgent | | | | Invitae-AR syndromic pediatric cardiomyopathy | | | Invitae-AR syndromic pediatric cardiomyopathy | |
| CRB1 | | | | | | | | | | | | (CRB1-related maculopathy with cystoid macular oedema) | | |
| CRYAB | | | moderate: | | (cardiomyopathy, hypertrophic) | fulgent | centogene | | GeneDx | | no excess | (Cardiomyopathy, dilated) | Invitae-primary panel | mayo_AD, AR_DCM, myofibrillar myopathy |
| CSRP3 | (HCM, test DCM/LVNC) | | strong: excess + human genetic evidence | no excess | (cardiomyopathy, hypertrophic) | fulgent | centogene | mayo_AD_HCM, DCM | GeneDx | Invitae-primary panel | no excess | (Cardiomyopathy, dilated) | Invitae-primary panel | mayo_AD_HCM, DCM |
| CTF1 | | | | | | fulgent | | | | | | (Cardiomyopathy, dilated) | Invitae-preliminary evidence | |
| CTLA4 | | | | | | | | | | | | (Dilated cardiomyopathy, association with) | | |
| CTNNA3 | | | | | (Hypertrophic cardiomyopathy or arrhythmogenic cardiomyopathy) | | | | GeneDx | | | (Cardiomyopathy, dilated) | Invitae-preliminary evidence | |
| DES | (ARVC, test DCM/LVNC) | | | | (cardiomyopathy, hypertrophic); (Hypertrophic cardiomyopathy with conduction system disease) | fulgent | centogene | mayo_AD, AR_DCM, ARVC, myofibrillar myopathy, RCM with AV block, neurogenic scapuloperoneal syndrome Kaeser type, LGMD | GeneDx | Invitae-primary panel | no excess | (Cardiomyopathy, dilated) | Invitae-primary panel | mayo_AD, AR_DCM, ARVC, myofibrillar myopathy, RCM with AV block, neurogenic scapuloperoneal syndrome Kaeser type, LGMD |
| DLG1 | | | | | (cardiomyopathy, hypertrophic) | | | | | | | | | |
| DMD | | (Bradycardia, left bundle branch block & left ventricular noncompaction) | | | | fulgent | | | GeneDx | | | (Cardiomyopathy, dilated, X-linked);(Muscular dystrophy, Becker / dilated cardiomyopathy, X-linked); (Muscular dystrophy, Becker with dilated cardiomyopathy & progressive heart failure); (Cardiomyopathy, dilated) | Invitae-primary panel | |
| DNAJC19 | | | | | | | | | | | | | Invitae-AR syndromic pediatric cardiomyopathy | |
| DOLK | | | | | | | | | GeneDx | | | (Severe ichthyosis, distal digital constrictions and dilated cardiomyopathy); (Cardiomyopathy, dilated) | Invitae-primary panel | |
| DSC2 | (ARVC, test DCM/LVNC); (secondary finding) | | | | | fulgent | | | GeneDx | | no excess | (Cardiomyopathy, dilated) | Invitae-primary panel | |
| DSG2 | (ARVC, test DCM/LVNC); (secondary finding) | | | | (cardiomyopathy, hypertrophic) | fulgent | | | GeneDx | | no excess | (Cardiomyopathy, dilated) | Invitae-primary panel | |
| DSP | (ARVC, test DCM/LVNC); (secondary finding) | | | | (Hypertrophic cardiomyopathy); (cardiomyopathy, hypertrophic); (arrhythmogenic CM or HCM) | fulgent | | | GeneDx | | Truncating excess | (Dilated cardiomyopathy, woolly hair, keratoderma); (Cardiomyopathy, dilated) | Invitae-primary panel | |
| DTNA | | (Left ventricular noncompaction with CHD) | | | | fulgent | | | GeneDx | | no excess | (Cardiomyopathy, dilated) | Invitae-preliminary evidence | |
| EEF1A2 | | | | | | | | | | | | (Dilated cardiomyopathy, failure to thrive, global developmental delay, epilepsy & early death) | | |
| EGFR | | | | | | | | | | | | (Dilated cardiomyopathy, association with) | | |
| ELAC2 | | | | | (Hypertrophic cardiomyopathy and complex I deficiency) | fulgent | | | | Invitae-AR syndromic pediatric cardiomyopathy | | | Invitae-AR syndromic pediatric cardiomyopathy | |
| EMD | | (Bradycardia, left bundle branch block & left ventricular noncompaction) | | | | fulgent | | | GeneDx | | no excess | (X-linked dilated cardiomyopathy with conduction defects & arrhythmias); (Cardiomyopathy, dilated) | Invitae-primary panel | |

| | | | | | | | | | | | | | | |
|---|---|---|---|---|---|---|---|---|---|---|---|---|---|---|
| EYA4 | | | | | | | | | GeneDx | | | (Dilated cardiomyopathy and sensorineural deafness); (Cardiomyopathy, dilated) | Invitae-primary panel | |
| FBXO32 | | | | | | | | | | | | (Cardiomyopathy, dilated) | | |
| FBXO7 | | | | | (Mean cell haemoglobin, association with) | | | | | | | | | |
| FHL1 | | | strong: excess + human genetic evidence | Nontruncating excess | (cardiomyopathy, hypertrophic); (X-linked myopathy with hypertrophic cardiomyopathy) | fulgent | | | GeneDx | Invitae-primary panel | no excess | | Invitae-primary panel | |
| FHL2 | | | functional data only | | (cardiomyopathy, hypertrophic) | | centogene | | | | no excess | (Cardiomyopathy, dilated) | Invitae-preliminary evidence | |
| FHOD3 | | | | | (cardiomyopathy, hypertrophic) | | | | | | | (Cardiomyopathy, dilated) | | |
| FKRP | | | | | | | | | GeneDx | | | (Cardiomyopathy, dilated) | Invitae-primary panel | |
| FKTN | | (Left ventricular noncompaction) | | | | fulgent | | | GeneDx | | | (Cardiomyopathy, dilated) | Invitae-primary panel | |
| FLNC | | (Cardiomyopathy, left ventricular noncompaction) | moderate: genetic evidence, excess not tested | | (cardiomyopathy, hypertrophic) | fulgent | centogene | | GeneDx | Invitae-primary panel | | (Cardiomyopathy, dilated) | Invitae-primary panel | |
| FLT1 | | | | | | | | | | | | (Cardiomyopathy, dilated) | | |
| FOXD4 | | | | | | | | | | | | (Dilated cardiomyopathy, OCD and suicidality) | | |
| FXN | | | functional data only | | (cardiomyopathy, hypertrophic) | fulgent | | | | | | | | |
| GAA | | | metabolic mimic | | (Glycogen storage disease type Ⅱ with hypertrophic cardiomyopathy);(Glycogen storage disease 2, atypical infantile-onset with hypertrophic cardiomyopathy); (cardiomyopathy, hypertrophic); (Glycogen storage disease 2 with hypertrophic cardiomyopathy) | fulgent | | | GeneDx | Invitae-primary panel | | | Invitae-primary panel | |
| GATA2 | | | | | | | | | | | | (GATA2 deficiency with melanoma);(MonoMAC and Emberger syndromes); (GATA2 deficiency) | | |
| GATA4 | | (Left ventricular noncompaction) | | | | fulgent | | | GeneDx | Invitae-preliminary evidence | | (Dilated cardiomyopathy, sporadic);(Cardiomyopathy, dilated) | Invitae-preliminary evidence | |
| GATA5 | | | | | | | | | | | | (Cardiomyopathy, dilated) | | |
| GATA6 | | | | | | | | | | | | (Cardiomyopathy, dilated) | Invitae-preliminary evidence | |
| GATAD1 | | | | | | fulgent | | | GeneDx | | | (Cardiomyopathy, dilated) | Invitae-preliminary evidence | |
| GLA | (HCM, test DCM/LVNC); (secondary finding) | | metabolic mimic | Nontruncating excess | (cardiomyopathy, hypertrophic); (Hypertrophic cardiomyopathy with Fabry disease) | fulgent | centogene | mayo_X-linked_Fabry disease | GeneDx | Invitae-primary panel | no excess | | Invitae-primary panel | |
| GTPBP3 | | | | | (Hypertrophic cardiomyopathy, lactic acidosis & encephalopathy) | | | | | | | | | |
| gtpbp3i5 | | | | | (Hypertrophic cardiomyopathy, lactic acidosis & encephalopathy) | | | | | | | | | |
| HAND1 | | | | | | | | | | | | (Cardiomyopathy, dilated) | | |
| HCN4 | | (Bradycardia & left ventricular noncompaction cardiomyopathy); (Bradycardia, left bundle branch block & left ventricular noncompaction) | | | | | | | GeneDx | | | | Invitae-primary panel | |
| HFE | | | | | (Mean cell haemoglobin, association with) | | | | GeneDx | | | | | |
| HIF1A | | | | | (More severe septal hypertrophy and diastolic dysfunction, in hypertrophic cardiomyopathy, association with) | | | | | | | | | |
| HRAS | | | | | | fulgent | | | GeneDx | Invitae-RASopathy | | | Invitae-RASopathy | |
| HSPB6 | | | | | | | | | | | | (Cardiomyopathy, dilated) | | |
| ILK | | | | | (cardiomyopathy, hypertrophic) | fulgent | | | GeneDx | | | (Cardiomyopathy, dilated) | Invitae-preliminary evidence | |
| INS-IGF2 | | | | | (cardiomyopathy, hypertrophic) | | | | | | | | | |
| ISL1 | | | | | | | | | | | | (Cardiomyopathy, dilated) | | |
| JPH2 | | | functional data only | | (cardiomyopathy, hypertrophic) | fulgent | centogene | | GeneDx | Invitae-preliminary evidence | | (Cardiomyopathy, dilated) | Invitae-preliminary evidence | |
| JUP | (ARVC, test DCM/LVNC) | | | | | | | | | | no excess | (Cardiomyopathy, dilated) | Invitae-primary panel | |
| KCNE1 | | | | | | | | | | | | (Cardiomyopathy, dilated) | | |
| KCNH2 | | | | | | | | | | | | (Cardiomyopathy, dilated) | | |
| KCNJ12 | | | | | | | | | | | | (Cardiomyopathy, dilated) | | |
| KCNQ1 | | (Long QT syndrome & left ventricular noncompaction) | HCM+arrythmia -> likely cause arrythmia only | | (cardiomyopathy, hypertrophic) | fulgent | | | GeneDx | | | (Cardiomyopathy, dilated) | | |
| KLF10 | | | functional data only | | (cardiomyopathy, hypertrophic) | | | | | | | (Cardiomyopathy, dilated) | | |
| KRAS | | | | | | fulgent | | | GeneDx | Invitae-RASopathy | | | Invitae-RASopathy | |
| KRT1 | | | | | (Ichthyosis hystrix, Curth-Macklin type) | | | | | | | | | |
| LAMA2 | | | | | | | | | | | | (Cardiomyopathy, dilated) | | |
| LAMA4 | | | | | (cardiomyopathy, hypertrophic) | fulgent | | | GeneDx | | no excess | (Cardiomyopathy, dilated) | Invitae-preliminary evidence | mayo_AD_DCM |
| LAMP2 | (HCM, test DCM/LVNC) | | metabolic mimic | no excess | (cardiomyopathy, hypertrophic) | fulgent | centogene | mayo_X-linked_Danon disease | GeneDx | Invitae-primary panel | no excess | (Cardiomyopathy, dilated) | Invitae-primary panel | mayo_X-linked_Danon disease |
| LDB3 | | | functional data only | | (cardiomyopathy, hypertrophic) | fulgent | centogene | | GeneDx | Invitae-preliminary evidence | no excess | (Cardiomyopathy, dilated) | Invitae-preliminary evidence | mayo_AD_DCM, LVNC, myofibrillar myopathy |
| ldb3i2 | | | | | | | | | | | | (Cardiomyopathy, dilated) | | |
| ldb3i6 | | | | | | | | | | | | (Cardiomyopathy, dilated) | | |
| ldb3z4 | | (Left ventricular noncompaction) | | | (cardiomyopathy, hypertrophic) | | | | | | | (Cardiomyopathy, dilated) | | |
| LMNA | (DCM, test LVNC) | (Cardiomyopathy, left ventricular noncompaction);(Left ventricular noncompaction) | no evidence (variant too common) | | (cardiomyopathy, hypertrophic) | fulgent | | | GeneDx | | Nontruncating excess and Truncating excess | (Atrioventricular block with normal QRS interval and dilated cardiomyopathy); (Dilated cardiomyopathy / ventricular arrhythmia); (Dilated cardiomyopathy and limb-girdle muscular dystrophy 1B); (Lipodystrophy, mandibular dysplasia and dilated cardiomyopathy);(Ovarian failure and dilated cardiomyopathy);(Partial lipodystrophy, dilated cardiomyopathy & conduction system disease); (Cardiomyopathy, dilated) | Invitae-primary panel | mayo_AD, AR_DCM, EMD, LGMD, congenital muscular dystrophy (see OMIM for full listing) |
| lmna2 | | | | | | | | | | | | (Cardiomyopathy, dilated) | | |
| LRRC10 | | | | | | | | | GeneDx | | | (Cardiomyopathy, dilated) | Invitae-preliminary evidence | |
| MAP2K1 | | | | | (cardiomyopathy, hypertrophic) | fulgent | | | GeneDx | Invitae-RASopathy | | | Invitae-RASopathy | |
| MAP2K2 | | | | | (cardiomyopathy, hypertrophic) | fulgent | | | GeneDx | Invitae-RASopathy | | | Invitae-RASopathy | |
| MED12 | | | | | | | | | | | | | Invitae-preliminary evidence | |
| MIB1 | | | | | (cardiomyopathy, hypertrophic) | | | | GeneDx | | | | | |
| MIB2 | | (Left ventricular noncompaction) | | | | | | | | | | | | |
| MIR196A2 | | | | | | | | | | | | (Dilated cardiomyopathy, association with) | | |
| MIR499A | | | | | | | | | | | | (Dilated cardiomyopathy, association with) | | |
| MRPL3 | | | rare, recessive, phenocopy | | (cardiomyopathy, hypertrophic) | | | | | | | | | |
| MRPL44 | | | | | (Hypertrophic cardiomyopathy, childhood-onset) | | | | | | | | | |
| MTND1 | | | | | | | | | GeneDx | | | | | |
| MTND5 | | | | | | | | | GeneDx | | | | | |
| MTND6 | | | | | | | | | GeneDx | | | | | |

| | | | | | | | | | | | | | | |
|---|---|---|---|---|---|---|---|---|---|---|---|---|---|---|
| MTO1 | | | | | (Hypertrophic cardiomyopathy, lactic acidosis and respiratory chain deficiency);(Hypertrophic cardiomyopathy & lactic acidosis) | fulgent | | | | Invitae-AR syndromic pediatric cardiomyopathy | | | Invitae-AR syndromic pediatric cardiomyopathy | |
| MTTD | | | | | | | | | GeneDx | | | | | |
| MTTG | | | | | | | | | GeneDx | | | | | |
| MTTH | | | | | | | | | GeneDx | | | | | |
| MTTI | | | | | | | | | GeneDx | | | | | |
| MTTK | | | | | | | | | GeneDx | | | | | |
| MTTL1 | | | | | | | | | GeneDx | | | | | |
| MTTL2 | | | | | | | | | GeneDx | | | | | |
| MTTM | | | | | | | | | GeneDx | | | | | |
| MTTQ | | | | | | | | | GeneDx | | | | | |
| MTTS1 | | | | | | | | | GeneDx | | | | | |
| MTTS2 | | | | | | | | | GeneDx | | | | | |
| MURC | | | | | | | | | GeneDx | | | | | |
| MYBPC3 | (HCM, test DCM/LVNC); (secondary finding) | (Cardiomyopathy, dilated with left ventricular noncompaction); (Cardiomyopathy, left ventricular noncompaction); (Hypertrophic cardiomyopathy and left ventricular noncompaction);(Left ventricular noncompaction) | sarcomere, validated | Nontruncating excess and Truncating excess | (Hypertrophic cardiomyopathy, in Potocki Schaffer syndrome); (Hypertrophic cardiomyopathy, apical);(Hypertrophic cardiomyopathy with inclusion body myositis);(Hypertrophic cardiomyopathy and left ventricular noncompaction); (Hypertrophic cardiomyopathy and atrial fibrillation); (cardiomyopathy, hypertrophic); (arrhythmogenic CM or HCM) | fulgent | centogene | mayo_AD_HCM, DCM | GeneDx | Invitae-primary panel | no excess | (Dilated cardiomyopathy, hearing loss & developmental delay); (Dilated cardiomyopathy); (Cardiomyopathy, dilated) | Invitae-primary panel | mayo_AD_HCM, DCM |
| MYBPHL | | | | | | | | | | | | (Dilated cardiomyopathy & arrhythmias) | | |
| MYH15 | | | | | (cardiomyopathy, hypertrophic) | | | | | | | | | |
| MYH6 | | | weak: excess but no strong segregation data | | (cardiomyopathy, hypertrophic); (Hypertrophic cardiomyopathy) | fulgent | centogene | | GeneDx | Invitae-preliminary evidence | no excess | (Cardiomyopathy, dilated) | Invitae-preliminary evidence | mayo__HCM, DCM |
| MYH7 | (HCM);(DCM, test LVNC);(secondary finding) | (Cardiomyopathy, left ventricular noncompaction);(Ebstein anomaly, left ventricular noncompaction, and ventricular septal defect); (Left ventricular noncompaction); (Cardiomyopathy, left ventricular noncompaction, with heart block);(Hypertrophic cardiomyopathy with left ventricular noncompaction cardiomyopathy and coronary artery-left ventricular fistulae);(Laing distal myopathy and left ventricular noncompaction cardiomyopathy) | sarcomere, validated | Nontruncating excess | (Hypertrophic cardiomyopathy, long QT and atrial fibrillation); (Hypertrophic cardiomyopathy with left ventricular noncompaction cardiomyopathy and coronary artery-left ventricular fistulae); (cardiomyopathy, hypertrophic); (Hypertrophic cardiomyopathy with cranial, axial and proximal myopathy) | fulgent | centogene | mayo_AD_HCM, DCM, LVNC, myopathy | GeneDx | Invitae-primary panel | Nontruncating excess | (Core myopathy, dilated cardiomyopathy, respiratory failure & scoliosis); (Cardiomyopathy, dilated) | Invitae-primary panel | mayo_AD_HCM, DCM, LVNC, myopathy |
| MYH7B | | (Cardiomyopathy, left ventricular noncompaction) | | | | | | | | | | | | |
| MYL2 | (HCM, test DCM/LVNC); (secondary finding) | | sarcomere, validated | Nontruncating excess | (cardiomyopathy, hypertrophic) | fulgent | centogene | mayo_AD_HCM | GeneDx | Invitae-primary panel | no excess | (Cardiomyopathy, dilated) | Invitae-primary panel | |
| MYL3 | (HCM, test DCM/LVNC); (secondary finding) | | sarcomere, validated | Nontruncating excess | (cardiomyopathy, hypertrophic); (Hypertrophic cardiomyopathy) | fulgent | centogene | mayo_AD, AR_HCM | GeneDx | Invitae-primary panel | no excess | (Cardiomyopathy, dilated) | Invitae-primary panel | |
| MYLK2 | | | functional data only | | (cardiomyopathy, hypertrophic) | fulgent | centogene | mayo_AD_HCM | GeneDx | Invitae-preliminary evidence | no excess | (Cardiomyopathy, dilated) | Invitae-preliminary evidence | |
| MYO6 | | | rare, recessive, phenocopy | | | | | | | | | | | |
| MYOM1 | | | functional data only | | (cardiomyopathy, hypertrophic) | fulgent | | | | Invitae-preliminary evidence | | (Cardiomyopathy, dilated) | Invitae-preliminary evidence | |
| MYOM3 | | | | | | | | | | | | (Cardiomyopathy, dilated) | | |
| MYOZ2 | | | moderate: no excess but human evidence -> very rare that mutation in this gene is causitive, most mutations benign | no excess | (cardiomyopathy, hypertrophic) | fulgent | | mayo_AD_HCM | GeneDx | Invitae-preliminary evidence | no excess | | Invitae-preliminary evidence | |
| MYPN | | | functional data only | | (cardiomyopathy, hypertrophic) | fulgent | centogene | | GeneDx | Invitae-preliminary evidence | | (Cardiomyopathy, dilated) | Invitae-preliminary evidence | mayo_AD_HCM, DCM |
| NAA10 | | | | | (Developmental delay & hypertrophic cardiomyopathy) | | | | | | | | | |
| NCOA6 | | | | | | | | | | | | (Cardiomyopathy, dilated) | | |
| NDUFAF1 | | | | | (Hypertrophic cardiomyopathy, fatal infantile) | | | | | | | | | |
| NDUFV2 | | | | | (Hypertrophic cardiomyopathy and encephalopathy) | | | | | | | (Cardiomyopathy, dilated) | | |
| NEB | | | | | (cardiomyopathy, hypertrophic) | | | | | | | | | |
| NEBL | | | | | (cardiomyopathy, hypertrophic) | fulgent | | | GeneDx | | | (Cardiomyopathy, dilated) | Invitae-preliminary evidence | |
| NEXN | | (Cardiomyopathy, left ventricular noncompaction) | functional data only | no excess | (cardiomyopathy, hypertrophic) | fulgent | centogene | mayo_AD_HCM, DCM | GeneDx | Invitae-preliminary evidence | no excess | (Cardiomyopathy, dilated) | Invitae-preliminary evidence | mayo_AD_HCM, DCM |
| NF1 | | | | | | fulgent | | | | Invitae-RASopathy | | | Invitae-RASopathy | |
| NFKB1 | | | | | | | | | | | | (Dilated cardiomyopathy risk) | | |
| NKX2-5 | | | | | | | | | GeneDx | | | (Dilated cardiomyopathy & arrhythmias); (Cardiomyopathy, dilated) | Invitae-preliminary evidence | |
| NPPA | | | | | | | | | | | | | Invitae-preliminary evidence | |
| NRAP | | | | | | | | | | | | (Cardiomyopathy, dilated) | | |
| NRAS | | | | | | fulgent | | | GeneDx | Invitae-RASopathy | | | Invitae-RASopathy | |
| OBSCN | | | no evidence (variant too common) | | (cardiomyopathy, hypertrophic) | | | | | | | (Cardiomyopathy, dilated) | | |
| obscntv2 | | | | | (cardiomyopathy, hypertrophic) | | | | | | | (Cardiomyopathy, dilated) | | |
| OBSL1 | | | | | | | | | | | | (Cardiomyopathy, dilated) | | |
| obsl1tv3 | | | | | | | | | | | | (Cardiomyopathy, dilated) | | |
| OPA1 | | | | | (Infantile mitochondrial encephalomyopathy, hypertrophic cardiomyopathy with optic atrophy) | | | | | | | | | |
| PARS2 | | | | | | | | | | | | (Microcephaly, early infantile epileptic encephalopathy, Leigh-like syndrome, dilated cardiomyopathy, and renal insufficiency) | | |
| PCCA | | | | | | | | | | | | (Propionic acidaemia with dilated cardiomyopathy, adult onset) | | |
| PCCB | | | | | | | | | | | | (Propionic acidaemia with dilated cardiomyopathy, adult onset) | | |
| PDLIM3 | | | no evidence | | (cardiomyopathy, hypertrophic) | fulgent | centogene | | GeneDx | Invitae-preliminary evidence | | (Cardiomyopathy, dilated) | Invitae-preliminary evidence | |
| PKD2 | | | | | | | | | | | | (Polycystic kidney disease and dilated cardiomyopathy) | | |
| PKP2 | (ARVC, test DCM/LVNC); (secondary finding) | | | | (cardiomyopathy, hypertrophic); (Hypertrophic cardiomyopathy) | fulgent | | | GeneDx | | no excess | (Cardiomyopathy, dilated) | Invitae-primary panel | |
| PLEKHM2 | | Cardiomyopathy, dilated with left ventricular noncompaction | | | | | | | | | | | Invitae-preliminary evidence | |
| PLN | (HCM);(DCM, test LVNC) | | strong: excess + human genetic evidence | Truncating excess | (cardiomyopathy, hypertrophic) | fulgent | centogene | mayo_AD_HCM, DCM | GeneDx | Invitae-primary panel | no excess | (Cardiomyopathy, dilated) | Invitae-primary panel | mayo_AD_HCM, DCM |
| PPCS | | | | | | | | | | | | (Cardiomyopathy, dilated) | | |
| PRDM16 | | (Left ventricular noncompaction) | | | | | | | GeneDx | | | (Cardiomyopathy, dilated) | Invitae-preliminary evidence | |

| | | | | | | | | | | | | | | |
|---|---|---|---|---|---|---|---|---|---|---|---|---|---|---|
| PRKAG2 | (HCM, test DCM/LVNC); (secondary finding) | | metabolic mimic | Nontruncating excess | (cardiomyopathy, hypertrophic) | fulgent | centogene | mayo_AD_HCM, Wolff-Parkinson-White syndrome | GeneDx | Invitae-primary panel | no excess | (Cardiomyopathy, dilated) | Invitae-primary panel | |
| PRNP | | | | | (cardiomyopathy, hypertrophic) | | | | | | | | | |
| PSEN1 | | | | | | | | | | | | (Cardiomyopathy, dilated) | | |
| PSEN2 | | | | | | | | | | | | (Dilated cardiomyopathy & heart failure); (Cardiomyopathy, dilated) | | |
| PTEN | | (Cardiomyopathy, left ventricular noncompaction) | | | | | | | | | | | | |
| PTPN11 | | | | | (Noonan syndrome with mulitple lentigines & hypertrophic cardiomyopathy); (cardiomyopathy, hypertrophic); (Noonan syndrome with juvenile myelomonocytic leukaemia & hypertrophic cardiomyopathy) | fulgent | | | GeneDx | Invitae-RASopathy | | | Invitae-RASopathy | |
| RAF1 | | | | | (cardiomyopathy, hypertrophic); (Perinatal problems & hypertrophic cardiomyopathy) | fulgent | | mayo_AD_Noonan/multiple lentigines syndrome | GeneDx | Invitae-RASopathy | | (Cardiomyopathy, dilated) | Invitae-primary panel | mayo_AD_Noonan/multiple lentigines syndrome, DCM |
| RANGRF | | | | | | | | | | | | (Cardiomyopathy, dilated) | | |
| RASA1 | | | | | | fulgent | | | | Invitae-RASopathy | | | Invitae-RASopathy | |
| RBM20 | (DCM, test LVNC) | | | | | fulgent | | | GeneDx | | | (Cardiomyopathy, dilated) | Invitae-primary panel | mayo_AD_DCM |
| RHO | | | | | | | | | | | | (Cystoid macular edema in retinitis pigmentosa) | | |
| RIT1 | | | | | (cardiomyopathy, hypertrophic) | fulgent | | | GeneDx | Invitae-RASopathy | | | Invitae-RASopathy | |
| RMND1 | | | | | | | | | | | | (Chronic kidney disease, dilated cardiomyopathy & neurological involvement) | | |
| RRAGC | | | | | | | | | | | | (Cardiomyopathy, dilated) | | |
| RRAS | | | | | | fulgent | | | | Invitae-RASopathy | | | Invitae-RASopathy | |
| RTKN2 | | | | | | | | | | | | (Cardiomyopathy, dilated) | | |
| RYR2 | (ARVC, test DCM/LVNC); (secondary finding) | (Left ventricular noncompaction) | | | (cardiomyopathy, hypertrophic) | fulgent | | | GeneDx | | no excess | (Cardiomyopathy, dilated) | Invitae-primary panel | |
| SCN5A | (DCM, test LVNC); (secondary finding) | | | | (cardiomyopathy, hypertrophic); (Hypertrophic cardiomyopathy) | fulgent | | | GeneDx | | Truncating excess | (Dilated cardiomyopathy and long QT syndrome);(Long QT syndrome 3 & dilated cardiomyopathy);(Long QT syndrome and dilated cardiomyopathy);(LQT3, conduction disorder, dilated cardiomyopathy);(Dilated cardiomyopathy); (Cardiomyopathy, dilated) | Invitae-primary panel | mayo_AD_Brugada syndrome, DCM, Heart block, LQTS, SSS, SIDS |
| scn5aib | | | | | | | | | | | | (Cardiomyopathy, dilated) | | |
| scn5aie | | | | | | | | | | | | (Cardiomyopathy, dilated) | | |
| SDHA | | | | | | | | | | | | | Invitae-AR syndromic pediatric cardiomyopathy | |
| SGCB | | | | | | | | | | | | (Cardiomyopathy, dilated) | | |
| SGCD | | | | | | fulgent | | | GeneDx | | no excess | (Cardiomyopathy, dilated) | Invitae-primary panel | mayo_AD, AR_DCM, LGMD |
| sgcdp | | | | | | | | | | | | (Cardiomyopathy, dilated) | | |
| SHOC2 | | | | | (Noonan syndrome with foetal distress & hypertrophic cardiomyopathy) | fulgent | | | GeneDx | Invitae-RASopathy | | | Invitae-RASopathy | |
| SLC22A5 | | | | | | | | | | | | | Invitae-primary panel | |
| SLC25A3 | | | | | (Muscular hypotonia & hypertrophic cardiomyopathy) | | | | | | | | | |
| SLC25A4 | | | | | (Mitochondrial myopathy & hypertrophic cardiomyopathy) | | centogene | | | | | | | |
| SLC25A5 | | | rare, recessive, phenocopy | | | | | | | | | | | |
| SLC35A2 | | | | | (Congenital disorder of glycosylation 2m with hypertrophic cardiomyopathy, hearing loss and short stature) | | | | | | | | | |
| SMC1A | | (Cornelia de Lange syndrome, left ventricular noncompaction cardiomyopathy, hyperopia and cleft lip) | | | | | | | | | | | | |
| SNTA1 | | | | | | | | | | | | (Cardiomyopathy, dilated) | | |
| SOD2 | | | | | (Multiple chemical sensitivity, association with) | | | | | | | | | |
| SOS1 | | | | | (cardiomyopathy, hypertrophic) | fulgent | centogene | | GeneDx | Invitae-RASopathy | | | Invitae-RASopathy | |
| SOS2 | | | | | | fulgent | | | | Invitae-RASopathy | | | Invitae-RASopathy | |
| SPEG | | | | | | | | | | | | (Centronuclear myopathy with dilated cardiomyopathy) | | |
| SPRED1 | | | | | | fulgent | | | | Invitae-RASopathy | | | Invitae-RASopathy | |
| SRI | | | no evidence (variant too common) | | | | | | | | | | | |
| STN1 | | | | | (Mean corpuscular hemoglobin, association with) | | | | | | | | | |
| SYNE1 | | | | | | | | | | | | (Cardiomyopathy, dilated) | | |
| SYNE2 | | | | | (cardiomyopathy, hypertrophic) | | | | | | | | | |
| SYNM | | | | | | | | | | | | (Cardiomyopathy, dilated) | | |
| TAF1A | | | | | | | | | | | | (Cardiomyopathy, dilated) | | |
| TAZ | | (Left ventricular noncompaction) | | | (cardiomyopathy, hypertrophic) | fulgent | | | GeneDx | | no excess | (Dilated cardiomyopathy, recurrent cardiac insufficiency, and multiple respiratory chain complex deficiency);(Barth syndrome with dilated cardiomyopathy); (Cardiomyopathy, dilated) | Invitae-primary panel | mayo_X-linked_Barth syndrome, LVNC, DCM |
| TBX20 | | | | | | | | | GeneDx | | | (Cardiomyopathy, dilated) | | |
| TBX5 | | | | | | | | | | | | (Cardiomyopathy, dilated) | | |
| TCAP | | | functional data only | | (cardiomyopathy, hypertrophic) | fulgent | centogene | mayo_AD, AR_HCM, DCM, LGMD | GeneDx | Invitae-primary panel | Nontruncating excess | (Cardiomyopathy, dilated) | Invitae-primary panel | mayo_AD, AR_HCM, DCM, LGMD |
| TCF21 | | | | | | | | | | | | (Cardiomyopathy, dilated) | | |
| TFRC | | | | | (Mean cell haemoglobin, association with) | | | | | | | | | |
| TGFB3 | | | | | | | | | GeneDx | | | (Cardiomyopathy, dilated) | Invitae-preliminary evidence | |
| TK2 | | | | | (Hypertrophic cardiomyopathy, regression of gross motor development, leucoencephalopathy and hepatic steatosis) | | | | | | | | | |
| TMED4 | | | | | | | | | | | | (Cardiomyopathy, dilated) | | |
| TMEM43 | (ARVC, test DCM/LVNC); (secondary finding) | | | | (cardiomyopathy, hypertrophic) | fulgent | | | GeneDx | | no excess | (Cardiomyopathy, dilated) | Invitae-primary panel | |
| TMEM70 | | | | | | | | | | | | | Invitae-AR syndromic pediatric cardiomyopathy | |
| TMPO | | | | | (cardiomyopathy, hypertrophic) | fulgent | | | GeneDx | | | (Cardiomyopathy, dilated) | Invitae-preliminary evidence | |
| TMPRSS6 | | | | | (Mean cell haemoglobin, association with) | | | | | | | | | |
| TNNC1 | (HCM);(DCM, test LVNC) | | weak: genetic evidence | no excess | (cardiomyopathy, hypertrophic) | fulgent | centogene | mayo_AD_HCM, DCM | GeneDx | Invitae-primary panel | Nontruncating excess | (Cardiomyopathy, dilated) | Invitae-primary panel | mayo_AD_HCM, DCM |
| TNNI3 | (HCM);(DCM, test LVNC);(secondary finding) | (Left ventricular noncompaction) | sarcomere, validated | Nontruncating excess | (cardiomyopathy, hypertrophic) | fulgent | centogene | mayo_AD, AR_DCM, HCM, RCM | GeneDx | Invitae-primary panel | no excess | (Cardiomyopathy, dilated) | Invitae-primary panel | mayo_AD, AR_DCM, HCM, RCM |
| TNNI3K | | | | | | | | | | | | (Conduction system disease, atrial tachyarrhythmia & dilated cardiomyopathy) | | |
| TNNT2 | (HCM);(DCM, test LVNC);(secondary finding) | (Cardiomyopathy, left ventricular noncompaction) | sarcomere, validated | Nontruncating excess and Truncating excess | (cardiomyopathy, hypertrophic) | fulgent | centogene | mayo_AD_HCM, DCM, RCM, LVNC | GeneDx | Invitae-primary panel | Nontruncating excess | (Dilated cardiomyopathy); (Cardiomyopathy, dilated) | Invitae-primary panel | mayo_AD_HCM, DCM, RCM, LVNC |
| tnnt2tv1 | | | | | (cardiomyopathy, hypertrophic) | | | | | | | | | |
| TOR1AIP1 | | | | | | | | | GeneDx | | | | | |

| | | | | | | | | | | | | | | |
|---|---|---|---|---|---|---|---|---|---|---|---|---|---|---|
| TPM1 | (HCM);(DCM, test LVNC);(secondary finding) | (Ebstein anomaly, left ventricular noncompaction & heart failure, early onset) | sarcomere, validated | Nontruncating excess | (cardiomyopathy, hypertrophic); (Hypertrophic cardiomyopathy with electrical instability typical of Brugada syndrome) | fulgent | centogene | mayo_AD_HCM, DCM, LVNC | GeneDx | Invitae-primary panel | Nontruncating excess | (Cardiomyopathy, dilated) | Invitae-primary panel | mayo_AD_HCM, DCM, LVNC |
| tpm1.3 | | | | | (cardiomyopathy, hypertrophic) | | | | | | | | | |
| tpm1.7b | | | | | (cardiomyopathy, hypertrophic) | | | | | | | | | |
| TRIM54 | | | no evidence | | (cardiomyopathy, hypertrophic); (Proximal muscle weakness & hypertrophic cardiomyopathy) | | | | | | | | | |
| TRIM55 | | | weak: excess but no strong segregation data, putative modifier | | (cardiomyopathy, hypertrophic) | | | | | | | | | |
| TRIM63 | | | weak: excess but no strong segregation data, putative modifier | | (cardiomyopathy, hypertrophic); (Proximal muscle weakness & hypertrophic cardiomyopathy) | | centogene | | | | | | | |
| TTN | (DCM, test LVNC) | | | | (cardiomyopathy, hypertrophic) | fulgent | centogene | mayo_AD, AR_HCM, DCM, myopathy | GeneDx | | Truncating excess | (Cardiomyopathy, dilated) | Invitae-primary panel | mayo_AD, AR_HCM, DCM, ARVC myopathy |
| ttnic | | (Cardiomyopathy, left ventricular noncompaction) | | | | | | | | | | (Dilated cardiomyopathy and heart failure); (Cardiomyopathy, dilated) | | |
| ttnltv | | | | | (cardiomyopathy, hypertrophic) | | | | | | | (Cardiomyopathy, dilated) | | |
| ttnnovex1 | | | | | | | | | | | | (Cardiomyopathy, dilated) | | |
| ttnnovex2 | | | | | | | | | | | | (Cardiomyopathy, dilated) | | |
| ttnnovex3 | | (Cardiomyopathy, left ventricular noncompaction) | | | | | | | | | | (Cardiomyopathy, dilated) | | |
| ttntvn2b | | | | | (cardiomyopathy, hypertrophic) | | | | | | | (Cardiomyopathy, dilated) | | |
| TTR | (HCM, test DCM/LVNC) | | metabolic mimic | no excess | (cardiomyopathy, hypertrophic) | fulgent | centogene | mayo_AD_Transthyretin-related amyloidosis | GeneDx | Invitae-primary panel | no excess | (Cardiomyopathy, dilated) | Invitae-primary panel | mayo_AD_Transthyretin-related amyloidosis |
| TXNRD2 | | | | | | | | | GeneDx | | | (Cardiomyopathy, dilated) | Invitae-preliminary evidence | |
| VCL | | (Cardiomyopathy, left ventricular noncompaction, with heart block) | no evidence (no specific phenotype) | | (cardiomyopathy, hypertrophic) | fulgent | centogene | mayo_AD_HCM, DCM | GeneDx | Invitae-primary panel | Truncating excess | (Cardiomyopathy, dilated) | Invitae-primary panel | mayo_AD_HCM, DCM |
| VEGFA | | | | | (More severe septal hypertrophy and diastolic dysfunction, in hypertrophic cardiomyopathy, association with) | | | | | | | | | |
| VPS13A | | | | | | | | | | | | (Chorea-acanthocytosis with dilated cardiomyopathy & myopathy) | | |
| xirp2tv1 | | | | | | | | | | | | (Dilated cardiomyopathy, modifier of) | | |
| YWHAE | | (Left ventricular noncompaction & hypoplasia of the corpus callosum) | | | | | | | | | | | | |
| ZBTB17 | | | | | | | | | | | | (Cardiomyopathy, dilated) | | |
| ZMPSTE24 | | | | | | | | | | | | (Metabolic syndrome, ectopic fat accumulation & dilated cardiomyopathy) | | |

**Table S5**

| ID (hg38, UCSC) | Gene.refGene | Func.refGene | ExonicFunc.refGene | avsnp150 | Patient | DiseasePatient | Pathogenicity | Comment | Tested | Found |
|---|---|---|---|---|---|---|---|---|---|---|
| chr1_228212169_228212170_TC_AA | OBSCN | exonic | stopgain | | 289 | HCM | UNCERTAIN SIGNIFICANCE | VUS but keep in list. This is two SNPs next to eachother. Each individually is common ~0.001 in Ashkenazi Jewish backgrounds, but indiviudally they change a hydrophobic amino acid to another hydrophobic amino acid (Phe to Tyr or Leu). Our two together cause a change to a stop codon (orig: TTC, each snp individually TAC or TTA, our is TAA). However, if these variants are in linkage disequilabrium, then it's possible they could be too common to be causative? PMID: 34601892 On truncating OBSCN mutations in HCM from 2021. Truncating OBSCN mutations in general enriched in HCM population and HCM patients with OBSCN truncating variants had higher chance of death than HCM patients without. All the variants in the study were of aa1150-8866 while ours is aa129. This is much more upstream. It could be an issue of nonsense mediated decay and not enough protein, but also an issue of dominant negative. The paper suggests the truncated protein may lack the motifs to interact wth signaling molecules or other members of the sarcomere. We checked the RNA and OBSCN is reduced in HCM vs Control, and even more reduced in 289. Additionally, this variant reported (PMID: 34957489)) in rhabdomyolysis patient (skeletal muscle disease). Patient had a second mutation in the other allele (all 6 patients in this paper had biallelic OBSCN mutaitons) that was a splicing variant. Like the other patients in this study, they did not have cardiomyopathy. However, they are all young. Patient is only 19 (age 12 at first onset), so cardiomyopathy may come later? | clin, wgs | Clin_NotInPanel, WGS_merge |
| chr12_21805252_21805254_TAA_ATTT | ABCC9 | exonic | frameshift stopgain | | 298 | HCM | UNCERTAIN SIGNIFICANCE | VUS but keep in list. This same mutation is in the literature (PMID: 15034580) in a DCM patient. Expression of the mutated gene in frog oocytes shows 70% reduction in channel trafficing. Mutation is in exon38, the c terminus. ATP-induced channel inhibition was blunted. "Thus, in contrast to the catalytic reaction in the wild type, where the rate-limiting step is ADP dissociation (k4), the Fs1524 ATPase is characterized by rate-limiting Pi dissociation". No segregation analysis. This same paper found a second DCM patient with a different mutation (missense) in exon38 (near the catalytic ATPase pocket) that also affected the channel function. (Since this is just one functional study, not enough for PS3). This variant also reported in PMID: 31638414. This was a study of very early onset atrial fibrilation. It was done at Stanford, but the patient in question was a white male. Our patient is a white famale, so it is not the same patient. It is possible they are family though. The patient from the paper also has a variant in ALMS1 listed as VUS/likely benign. PMID: 24439875 - Reports on a male with brugada syndrome (diagnosed at age 39) with this mutation. This paper finds 11 probands for Brugada/early repolarization with a total of 8 ABCC9 mutations. Didn't examine our mutation further, but for the others they did functional studies showing that 3 of the variants like the DCM paper blunted the ATP channel inhibition and were gain of function. These other mutations were missense SNPs. | clin, wgs | Clin_NotInPanel, WGS_merge |
| chr11_47348424_47348424_C_T | MYBPC3 | exonic | nonsynonymous SNV | rs397516074 | 319 | HCM | PATHOGENIC | This variant found in 7 patients in our cohort: 3 unrelated HCM patients (319, 431, 582) + 2 related HCM patients (711, 712) + 2 unrelated DCM patients (811, 814). Note that we have an additional HCM patient (693) where the dad carryied this variant but our patient did not (hence they were sick for a different reason). This does not mean this variant wasn't pathogenic in the dad, as 693 could have gotten their pathogenic mutation from their mom. | wgs | WGS_P |
| chr6_129460287_129460287_C_T | LAMA2 | exonic | stopgain | rs398123383 | 320 | DCM | LIKELEY PATHOGENIC | Gene not in CardioClassifier | clin, wgs | Clin_NotInPanel, WGS_P |
| chr14_23425814_23425814_G_A | MYH7 | exonic | nonsynonymous SNV | rs121913630 | 333 | HCM | PATHOGENIC | | clin, wgs | Clin_P, WGS_P |
| chr15_63042893_63042893_G_A | TPM1 | exonic | nonsynonymous SNV | rs397516382 | 334 | HCM | UNCERTAIN SIGNIFICANCE | VUS but keep in list. Multiple clinvar entries and references to presence in cohorts in the literature. | clin, wgs | Clin_P, WGS_P |
| chr10_99727098_99727098_G_C | COX15 | exonic | stopgain | rs149718203 | 356 | HCM | UNCERTAIN SIGNIFICANCE | VUS but keep in list. ClinVar has 3 Pathogenic entries for Leigh Syndrome. The original reporting of the leigh syndrome patient did not have cardiomyopathy, though Leigh Syndrome can co-occur with cardiomyopathy. Literature does show the variant affects mitrochondrial output, etc. However, Leigh Syndrome is very rare and incidence in some of the studies on gnomAD show a higher frequency | clin, wgs | Clin_NotInPanel, WGS_Trunc |
| chr2_178578066_178578066_G_A | TTN | exonic | stopgain | rs371678190 | 367 | DCM | LIKELEY PATHOGENIC | | wgs | WGS_P |
| chr11_47348541_47348541_C_G | MYBPC3 | exonic | nonsynonymous SNV | rs397516068 | 371 | HCM | LIKELEY PATHOGENIC | | clin, wgs | Clin_P, WGS_P |
| chr11_47333306_47333306_-_G | MYBPC3 | exonic | frameshift insertion | rs730880669 | 373 | HCM | LIKELEY PATHOGENIC | | clin, wgs | Clin_P, WGS_P |
| chr2_178633009_178633009_A_- | TTN | exonic | stopgain | . | 399 | DCM | LIKELEY PATHOGENIC | | wgs | WGS_Freq |
| chr11_47346212_47346212_-_T | MYBPC3 | exonic | frameshift insertion | rs730880723 | 419 | HCM | LIKELEY PATHOGENIC | | clin, wgs | Clin_P, WGS_P |
| chr6_118558947_118558947_G_A | PLN | exonic | nonsynonymous SNV | rs754782171 | 419 | HCM | UNCERTAIN SIGNIFICANCE | VUS but keep in list. Also reported in the literature. | clin, wgs | Clin_NotInPanel, WGS_Freq |
| chr14_23425768_23425768_C_G | MYH7 | exonic | nonsynonymous SNV | rs730880894 | 420 | HCM | UNCERTAIN SIGNIFICANCE | VUS but keep in list. Reported in literature in other HCM patients. Found in 1 HCM patient (PMID:27247418). Our mutation is p. Ser738Thr, p.Ser738Arg was found in 1 HCM patient and her HCM son (PMID: 31416728) and not in her two children lacking HCM. Note that Ser and Thr are both polar while Arg (from the paper) is positive so it's a bigger change. pSer738Cys is also in ClinVar. The Arg and Cys versions are also rare. We find this variant in our patient 697 as well. | wgs | WGS_P |
| chr14_23425980_23425980_C_T | MYH7 | exonic | nonsynonymous SNV | rs121913638 | 421 | HCM | PATHOGENIC | | clin, wgs | Clin_P, WGS_P |
| chr2_178553559_178553560_CA_- | TTN | exonic | frameshift deletion | . | 428 | DCM | LIKELEY PATHOGENIC | | wgs | WGS_Freq |
| chr11_47348424_47348424_C_T | MYBPC3 | exonic | nonsynonymous SNV | rs397516074 | 431 | HCM | PATHOGENIC | This variant found in 7 patients in our cohort: 3 unrelated HCM patients (319, 431, 582) + 2 related HCM patients (711, 712) + 2 unrelated DCM patients (811, 814). Note that we have an additional HCM patient (693) where the dad carryied this variant but our patient did not (hence they were sick for a different reason). This does not mean this variant wasn't pathogenic in the dad, as 693 could have gotten their pathogenic mutation from their mom. | clin, wgs | Clin_P, WGS_P |
| chr2_178632166_178632167_TT_- | TTN | exonic | frameshift deletion | rs794729316 | 437 | DCM | LIKELEY PATHOGENIC | | clin, wgs | clin_research, Clin_VUS, WGS_VUS, WGS_Freq |
| chr11_47346379_47346379_C_T | MYBPC3 | intronic | . | rs397516083 | 438 | HCM | LIKELEY PATHOGENIC | | clin, wgs | Clin_NotInPanel, WGS_P |
| chr14_23424839_23424839_C_T | MYH7 | exonic | nonsynonymous SNV | rs36211715 | 440 | HCM | PATHOGENIC | | wgs | WGS_P |
| chr10_110812295_110812295_C_T | RBM20 | exonic | nonsynonymous SNV | rs747880281 | 495 | DCM | LIKELEY PATHOGENIC | CardioClassifier said VUS and only had PM2, PP2. I added PS3 due to (PMID: 32905764) | wgs | WGS_Freq |
| chr6_7580339_7580342_GAAG_- | DSP | exonic | frameshift deletion | . | 496 | HCM | LIKELEY PATHOGENIC | | clin, wgs | Clin_NotInPanel, WGS_Trunc |
| chr11_47335120_47335120_G_A | MYBPC3 | exonic | stopgain | rs387907267 | 502 | HCM | PATHOGENIC | | clin, wgs | Clin_P, WGS_P |
| chr10_110812298_110812298_G_A | RBM20 | exonic | nonsynonymous SNV | rs267607001 | 503 | DCM | LIKELEY PATHOGENIC | | clin, wgs | Clin_P, WGS_P |
| chr14_23426810_23426810_G_A | MYH7 | exonic | nonsynonymous SNV | rs727503263 | 512 | HCM | LIKELEY PATHOGENIC | CardioClassifier was PM2, PP3, PM1=VUS. ClinVar was also PS4 = likely pathogenic. ClinVar PS4 cites multiple publications of patients with this mutation. | clin, wgs | Clin_P, WGS_P |
| chr11_47332094_47332104_ACACCGTGCCT_CAGG | MYBPC3 | | | rs1085307897 | 517 | HCM | LIKELEY PATHOGENIC | No functional data. Likely does not cause NMD (nonsense mediated decay) but alters final amino acid sequence. Two other pathogenic/likely pathogenic ClinVar entries downtream that are frameshift or termination, however only one has good annotation. Because of the R1271X downstream variant good annotation keep this as likely pathogenic. | clin, wgs | Clin_P, WGS_merge, WGS_Freq |
| chr14_23431468_23431468_C_T | MYH7 | exonic | nonsynonymous SNV | rs3218713 | 534 | HCM | PATHOGENIC | | clin, wgs | Clin_P, WGS_P |
| chr2_178620017_178620017_-_T | TTN | exonic | frameshift insertion | rs1060500443 | 541 | DCM | LIKELEY PATHOGENIC | | wgs | WGS_Freq |

| | | | | | | | | | | |
|---|---|---|---|---|---|---|---|---|---|---|
| chr14_23417152_23417152_C_G | MYH7 | | | rs876661374 | 543 | HCM | UNCERTAIN SIGNIFICANCE | VUS but keep in list. ClinVar entry by Invitae maps out that: "Variants that disrupt the donor or acceptor splice site typically lead to a loss of protein function (PMID: 16199547), however the current clinical and genetic evidence is not sufficient to establish whether loss-of-function variants in MYH7 cause disease. Disruption of this splice site has been observed in individual(s) with autosomal dominant distal myopathy and/or hypertrophic cardiomyopathy (PMID: 30297972, 32403337)." We do not have cardiomyocyte RNA-seq data for this sample. | clin, wgs | clin_research, WGS_vqsr |
| chr11_47342698_47342698_G_A | MYBPC3 | exonic | nonsynonymous SNV | rs375882485 | 544 | HCM | PATHOGENIC | | clin, wgs | Clin_P, WGS_P |
| chr11_47333226_47333226_-_C | MYBPC3 | exonic | frameshift insertion | rs397516014 | 546 | HCM | LIKELEY PATHOGENIC | | clin, wgs | Clin_P, WGS_P |
| chr1_77933402_77933402_C_T | NEXN | exonic | stopgain | rs750076188 | 546 | HCM | UNCERTAIN SIGNIFICANCE | VUS but keep in list. | clin, wgs | Clin_NotInPanel, WGS_Trunc |
| chr14_23424107_23424107_G_C | MYH7 | exonic | nonsynonymous SNV | rs121913631 | 548 | HCM | PATHOGENIC | | wgs | WGS_P |
| chr11_47333192_47333192_A_C | MYBPC3 | splicing | . | rs387906397 | 550 | HCM | PATHOGENIC | Mutation also in patient 591 and 612 (also with HCM) | clin, wgs | Clin_P, WGS_P |
| chrX_101398942_101398942_T_C | GLA | exonic | nonsynonymous SNV | rs28935197 | 552 | Other (Fabry is HCM lookalike) | LIKELEY PATHOGENIC | | clin, wgs | Clin_P, WGS_P |
| chr11_47351255_47351256_GA_- | MYBPC3 | exonic | frameshift deletion | rs1057517766 | 562 | HCM | LIKELEY PATHOGENIC | | clin, wgs | Clin_P, WGS_P |
| chr14_23429278_23429278_C_T | MYH7 | exonic | nonsynonymous SNV | rs121913624 | 564 | HCM | PATHOGENIC | | clin, wgs | Clin_P, WGS_P |
| chr14_23425316_23425316_C_T | MYH7 | exonic | nonsynonymous SNV | rs3218716 | 565 | HCM | PATHOGENIC | | clin, wgs | Clin_P, WGS_P |
| chr1_228340511_228340511_C_G | OBSCN | rs576488972 | exonic | stopgain | 565 | HCM | UNCERTAIN SIGNIFICANCE | VUS: not in clinvar, not enough data, high freq in 1000genomes substudy, but only 694 patients so not sure; no LOF P/LP mutations in this gene in ClinVar to help annotate. But there is literature on truncating OBSCN mutations in HCM (PMID: 346018920). Also this patient has a pathogenic mutation in MYH7. (This plus high frequency suggests at most it would be modifying). Keep in because of other OBSCN stopgain variant in our cohort to compare to. | clin, wgs | Clin_NotInPanel, WGS_Trunc |
| chr18_3151790_3151790_G_A | MYOM1 | exonic | stopgain | rs765191680 | 568 | HCM | UNCERTAIN SIGNIFICANCE | VUS but keep in list. No pathogenic / likely pathogenic mutations in MYOM1 in ClinVar. Very few in literature. This truncation occurs only 35% of the way into the protein. We do not have RNA-seq data for this line to check expression. There is not enough data to link loss of fuction MYOM1 to HCM. Additionally, PMID: 33452765 shows knock out in hES-derived cardiomyoyctes have an atrophy phenotype, not an HCM phenotype. | clin, wgs | clin_research, WGS_VUS |
| chr14_23424855_23424855_T_C | MYH7 | exonic | nonsynonymous SNV | rs730880749 | 569 | HCM | LIKELEY PATHOGENIC | CardioClassifier was PM2, PP3, PM1=VUS; ClinVAR was PM2, PM1, PS4 = likely pathogenic. ClinVar was not PP3, however by my evaluation, including CADD score, it is bad. ClinVar PS4 cites PMID: 27247418. This is the stanford structure paper. I do not count this alone, as it didn't solve causality, but they also site communication with GeneDx and Invitae to increase HCM count. They also site PP1 for segregation in 2 families but site private communication with GeneDx and Inivate. This submission by ClinGen Cardiomyopathy Variant Curation Expert Panel an FDA recognized database. This variant found in this patient's sister, patient 570, also in our study. | clin, wgs | Clin_VUS, WGS_P |
| chr14_23424855_23424855_T_C | MYH7 | exonic | nonsynonymous SNV | rs730880749 | 570 | HCM | LIKELEY PATHOGENIC | CardioClassifier was PM2, PP3, PM1=VUS; ClinVAR was PM2, PM1, PS4 = likely pathogenic. ClinVar was not PP3, however by my evaluation, including CADD score, it is bad. ClinVar PS4 cites PMID: 27247418. This is the stanford structure paper. I do not count this alone, as it didn't solve causality, but they also site communication with GeneDx and Invitae to increase HCM count. They also site PP1 for segregation in 2 families but site private communication with GeneDx and Inivate. This submission by ClinGen Cardiomyopathy Variant Curation Expert Panel an FDA recognized database. This variant found in this patient's sister, patient 559, also in our study. | clin, wgs | Clin_VUS, WGS_P |
| chr11_47338680_47338680_C_T | MYBPC3 | splicing | . | rs727504334 | 571 | HCM | LIKELEY PATHOGENIC | | clin, wgs | Clin_P, WGS_P |
| chr10_119672446_119672446_C_A | BAG3 | exonic | stopgain | rs876661342 | 574 | DCM | LIKELEY PATHOGENIC | | clin, wgs | Clin_P, WGS_P |
| chr14_23424876_23424876_G_A | MYH7 | exonic | nonsynonymous SNV | rs2754158 | 581 | HCM | UNCERTAIN SIGNIFICANCE | VUS but keep in list. No ClinVar entry. | clin, wgs | Clin_Absent, WGS_P |
| chr11_47348424_47348424_C_T | MYBPC3 | exonic | nonsynonymous SNV | rs397516074 | 582 | HCM | PATHOGENIC | This variant found in 7 patients in our cohort: 3 unrelated HCM patients (319, 431, 582) + 2 related HCM patients (711, 712) + 2 unrelated DCM patients (811, 814). Note that we have an additional HCM patient (693) where the dad carryied this variant but our patient did not (hence they were sick for a different reason). This does not mean this variant wasn't pathogenic in the dad, as 693 could have gotten their pathogenic mutation from their mom. | clin, wgs | Clin_P, WGS_P |
| chr11_47333192_47333192_A_C | MYBPC3 | splicing | . | rs387906397 | 591 | HCM | PATHOGENIC | Mutation also in patient 550 and 612 (also with HCM) | clin, wgs | Clin_P, WGS_P |
| chr14_23426809_23426809_C_T | MYH7 | exonic | nonsynonymous SNV | rs730880883 | 592 | HCM | UNCERTAIN SIGNIFICANCE | VUS but keep in list. Also reported in the literature. | wgs | WGS_Freq |
| chr11_47342621_47342621_-_AGTG | MYBPC3 | exonic | frameshift insertion | rs730880712 | 593 | HCM | LIKELEY PATHOGENIC | | clin, wgs | Clin_P, WGS_VUS, WGS_Freq |
| chr2_178537711_178537711_C_A | TTN | exonic | stopgain | rs886038825 | 596 | DCM | LIKELEY PATHOGENIC | This mutation also found in another DCM patient, #754. | clin, wgs | Clin_P, WGS_P |
| chr11_47346331_47346331_C_T | MYBPC3 | exonic | stopgain | rs727503211 | 598 | HCM | LIKELEY PATHOGENIC | | clin, wgs | Clin_P, WGS_P |
| chr1_201363390_201363390_C_T | TNNT2 | exonic | nonsynonymous SNV | rs45501500 | 601 | DCM | UNCERTAIN SIGNIFICANCE | VUS but keep in list. Multiple ClinVar entries and some functional data that may not be clinically relevant + occurs in several cohort studies | clin, wgs | Clin_VUS, WGS_P |
| chr14_23426833_23426833_C_T | MYH7 | exonic | nonsynonymous SNV | rs371898076 | 603 | HCM | PATHOGENIC | | clin, wgs | Clin_P, WGS_P |
| pkp2dup | PKP2 | | | | 605 | DCM | UNCERTAIN SIGNIFICANCE | VUS but keep in list. PKP2 deletion has been found to segregate in ARVC family (PMID: 23486541) and has been found in other unrelated probands as well as patients with partial deletios (PMID: 29038103). We haven't found literature for a duplication. The deletion could affect the phenotype even if not caustive. | clin, wgs | clin_research, WGS_class |
| chr14_23424907_23424909_CTT_- | MYH7 | exonic | nonframeshift deletion | rs397516155 | 607 | HCM | LIKELEY PATHOGENIC | | clin, wgs | Clin_P, WGS_P |
| chr11_47333192_47333192_A_C | MYBPC3 | splicing | . | rs387906397 | 612 | HCM | PATHOGENIC | Mutation also in patient 591 and 550 (also with HCM) | clin, wgs | Clin_P, WGS_P |
| chr2_151494190_151494191_CT_- | NEB | exonic | frameshift deletion | rs755863625 | 612 | HCM | UNCERTAIN SIGNIFICANCE (rasopathy) | VUS but keep in list. VUS for Rasopathy. Nemaline myopathy more often inherited as recessive, but can be dominant. Usually not associated with cardiomyopathy but examples of both HCM and DCM. Multiple pathogenic ClinVar entries for nemaline myopathy for this variant. Downstream truncating variants also cause the disease. | clin, wgs | Clin_NotInPanel, WGS_Trunc |
| chr11_47343281_47343281_C_T | MYBPC3 | intronic | . | rs587776699 | 613 | HCM | LIKELEY PATHOGENIC | Variant not in CardioClassifier. This is based on ClinVar. Somewhat depenent on the assays from only one study. | wgs | WGS_P |
| chr15_84857913_84857913_C_T | ALPK3 | exonic | stopgain | rs749465164 | 613 | HCM | PATHOGENIC | | clin, wgs | Clin_NotInPanel, WGS_Trunc |
| chr2_178570469_178570469_T_- | TTN | exonic | frameshift deletion | rs1131691542 | 624 | DCM | LIKELEY PATHOGENIC | | clin, wgs | Clin_P, WGS_P |
| chr19_4117563_4117563_C_- | MAP2K2 | exonic | frameshift deletion | rs777549760 | 634 | HCM | UNCERTAIN SIGNIFICANCE | VUS but keep in list as potential modifier. There are 3 ClinVar entries for cardiac/rasopathies. Not good population data or mechanistic data. Likely loss of function (LOF) but not clear link for LOF to disease. Sarcomeric mutations are more common for adult-onset HCM, while these rasopathy genes are more common in pediatric cases. In pediatric cases, map2k2 rasopathy associated with cardiofaciouctaneuous syndrome, associated with HCM in 15-40% of cases and pulmonvary valve stenosis in 65-100% of cases. In these pediatric rasopothies the HCM is associated wth more severe left ventricular outflow tract obstruction (like our patient has) (PMID: 33718303), however the EMR we were given doesn't indicate childhood onset, nor any other rasopathy symptoms. Thus, keep this mutation in the pool as a modifier, but not as a likely P/LP for the patient's HCM. | clin, wgs | Clin_NotInPanel, WGS_Trunc |
| chr11_47332569_47332569_G_- | MYBPC3 | exonic | frameshift deletion | rs397516029 | 638 | HCM | LIKELEY PATHOGENIC | | clin, wgs | Clin_P, WGS_P |
| chr1_156136024_156136024_C_T | LMNA | exonic | stopgain | . | 643 | DCM | LIKELEY PATHOGENIC | | wgs | WGS_Freq |
| chr2_219568063_219568063_-_T | OBSL1 | exonic | frameshift insertion | rs762334954 | 650 | DCM | PATHOGENIC (Three M syndrome 2) | Pathogenic for Three M syndrome 2. Annotation dependent on ClinVar, as we don't study this disease. However, this is a recessive disorder and our patient is a heterozygote. From what we can tell, this syndrome has normal lifespan and cardiac dysfunction is not a major symptom. No OBSL1 variants in ClinVar for cardiomyopathy, so for DCM, we consider this a VUS. | clin, wgs | Clin_NotInPanel, WGS_Trunc |

| | | | | | | | | | | |
|---|---|---|---|---|---|---|---|---|---|---|
| chr14_23424951_23424951_A_G | MYH7 | exonic | nonsynonymous SNV | rs730880746 | 662 | HCM | UNCERTAIN SIGNIFICANCE | VUS but keep in list. Also reported in the literature. | clin, wgs | Clin_VUS, WGS_Freq |
| chr14_23431641_23431641_C_T | MYH7 | exonic | nonsynonymous SNV | rs1057517773 | 676 | HCM | UNCERTAIN SIGNIFICANCE | VUS but keep in list. Reported in literature in other HCM patients - PMID: 27247418, 27532257. These papers also report the p.Ala226Val mutation at the same residue. These are statistics papers looking at mutations in large cohorts. They don't determine pathogenicity. Also found in 1 HCM patient in PMID: 27600940. Paper DOI: 10.3390 /ijms17081239 - Also found in a patient with early-onset HCM in conjunction with a likely benign missense mutation in GLA (rs28935490). And found in 1 HCM patient in PMID: 26656175. (This paper and the last one with the DOI are both Italian.) This group did a follow-up paper (PMID: 27054166) that found the mutation likely causes "steric hindrance" | clin, wgs | Clin_P, WGS_P |
| chr2_178598977_178598977_-_T | TTN | exonic | frameshift insertion | rs397517626 | 677 | DCM | LIKELEY PATHOGENIC | | clin, wgs | Clin_VUS, WGS_P |
| chr1_201364341_201364341_C_T | TNNT2 | exonic | nonsynonymous SNV | rs397516466 | 684 | LVNC | LIKELEY PATHOGENIC | Note patient from paper cited in ClinVar with functional data was 70 at age of onset vs the average age of TNNT2 patients in their 20s. CardioClassifer said VUS since didn't have the PS3 functional note. | clin, wgs | Clin_VUS, WGS_P |
| chr14_23425768_23425768_C_G | MYH7 | exonic | nonsynonymous SNV | rs730880894 | 697 | HCM | UNCERTAIN SIGNIFICANCE | VUS but keep in list. Reported in literature in other HCM patients. Found in 1 HCM patient (PMID:27247418). Our mutation is p. Ser738Thr, p.Ser738Arg was found in 1 HCM patient and her HCM son (PMID: 31416728) and not in her two children lacking HCM. Note that Ser and Thr are both polar while Arg (from the paper) is positive so it's a bigger change. pSer738Cys is also in ClinVar. The Arg and Cys versions are also rare. We find this variant in our patient 420 as well. | clin, wgs | Clin_VUS, WGS_P |
| chr2_178740646_178740646_G_T | TTN | exonic | stopgain | rs370912401 | 709 | DCM | LIKELEY PATHOGENIC | | wgs | WGS_Freq |
| chr7_128837228_128837228_C_T | FLNC | exonic | stopgain | . | 710 | DCM | LIKELEY PATHOGENIC | | clin, wgs | Clin_NotInPanel, WGS_Trunc |
| chr11_47348424_47348424_C_T | MYBPC3 | exonic | nonsynonymous SNV | rs397516074 | 711 | HCM | PATHOGENIC | This variant found in 7 patients in our cohort: 3 unrelated HCM patients (319, 431, 582) + 2 related HCM patients (711, 712) + 2 unrelated DCM patients (811, 814). Note that we have an additional HCM patient (693) where the dad carryied this variant but our patient did not (hence they were sick for a different reason). This does not mean this variant wasn't pathogenic in the dad, as 693 could have gotten their pathogenic mutation from their mom. | clin_fam, wgs | clin_fam, WGS_P |
| chr11_47348424_47348424_C_T | MYBPC3 | exonic | nonsynonymous SNV | rs397516074 | 712 | HCM | PATHOGENIC | This variant found in 7 patients in our cohort: 3 unrelated HCM patients (319, 431, 582) + 2 related HCM patients (711, 712) + 2 unrelated DCM patients (811, 814). Note that we have an additional HCM patient (693) where the dad carryied this variant but our patient did not (hence they were sick for a different reason). This does not mean this variant wasn't pathogenic in the dad, as 693 could have gotten their pathogenic mutation from their mom. | wgs | WGS_P |
| chr2_178546612_178546612_G_A | TTN | exonic | stopgain | rs1060500435 | 715 | DCM | LIKELEY PATHOGENIC | | clin, wgs | Clin_P, WGS_P |
| chr2_178569734_178569735_TA_- | TTN | exonic | frameshift deletion | rs794729342 | 721 | DCM | LIKELEY PATHOGENIC | | clin, wgs | Clin_VUS, WGS_P |
| chr14_23424148_23424148_T_C | MYH7 | exonic | nonsynonymous SNV | rs397516161 | 738 | HCM | PATHOGENIC | | clin, wgs | Clin_P, WGS_P |
| chr1_156136093_156136093_C_T | LMNA | exonic | nonsynonymous SNV | rs397517889 | 741 | DCM | LIKELEY PATHOGENIC | CardioClassifer said VUS since didn't have the PS4 note. There are multiple pathogenic / likely pathogenic ClinVar entries and multiple published cohorts where this mutation, and another missense mutation at this amino acid, are documented | clin, wgs | Clin_P, WGS_P |
| chr2_178537711_178537711_C_A | TTN | exonic | stopgain | rs886038825 | 754 | DCM | LIKELEY PATHOGENIC | This mutation also found in another DCM patient, #596. | clin, wgs | Clin_VUS, WGS_P |
| chr15_48445467_48445467_G_T | FBN1 | exonic | stopgain | rs363806 | 766 | DCM | LIKELEY PATHOGENIC (Marfan syndrome) | for Marfan syndrome | clin, wgs | Clin_P, WGS_NotInPanel |
| chrM_3243_3243_A_G | tRNA Leu (UUR) | | | rs199474657 | 769 | DCM | LIKELEY PATHOGENIC | Multiple ClinVar entries show this is pathogenic for MELAS. This patient has a family history of MELAS symptoms as well as some of her own MELAS symptoms. Heart failure most likely developed due to that. | clin, wgs | Clin_P, WGS_NotInPanel |
| chrX_136206492_136206492_G_- | FHL1 | exonic | frameshift deletion | rs1060502840 | 772 | DCM | LIKELEY PATHOGENIC | | clin, wgs | Clin_P, WGS_vqsr |
| chr2_178563474_178563474_C_- | TTN | exonic | frameshift deletion | rs794729349 | 780 | DCM | LIKELEY PATHOGENIC | | clin, wgs | Clin_VUS, WGS_P |
| chr2_178542803_178542803_-_T | TTN | exonic | frameshift insertion | rs794729365 | 783 | DCM | LIKELEY PATHOGENIC | | wgs | WGS_P |
| chr2_178607429_178607429_T_- | TTN | exonic | frameshift deletion | . | 788 | DCM | LIKELEY PATHOGENIC | | wgs | WGS_Freq |
| chr11_47348424_47348424_C_T | MYBPC3 | exonic | nonsynonymous SNV | rs397516074 | 811 | DCM | PATHOGENIC | This variant found in 7 patients in our cohort: 3 unrelated HCM patients (319, 431, 582) + 2 related HCM patients (711, 712) + 2 unrelated DCM patients (811, 814). Note that we have an additional HCM patient (693) where the dad carryied this variant but our patient did not (hence they were sick for a different reason). This does not mean this variant wasn't pathogenic in the dad, as 693 could have gotten their pathogenic mutation from their mom. | clin, wgs | Clin_P, WGS_P |
| chr11_47348424_47348424_C_T | MYBPC3 | exonic | nonsynonymous SNV | rs397516074 | 814 | DCM | PATHOGENIC | This variant found in 7 patients in our cohort: 3 unrelated HCM patients (319, 431, 582) + 2 related HCM patients (711, 712) + 2 unrelated DCM patients (811, 814). Note that we have an additional HCM patient (693) where the dad carryied this variant but our patient did not (hence they were sick for a different reason). This does not mean this variant wasn't pathogenic in the dad, as 693 could have gotten their pathogenic mutation from their mom. | clin, wgs | Clin_NotInPanel, WGS_P |
| chr10_119651774_119651781_GACCGGCT_- | BAG3 | exonic | frameshift deletion | rs727505283 | 815 | DCM | LIKELEY PATHOGENIC | | clin, wgs | Clin_P, WGS_P |
| chr1_99851059_99851060_AG_- | AGL | exonic | frameshift deletion | rs113994127 | 820 | Healthy Control | LIKELEY PATHOGENIC | Likely pathogenic for Glycogen Storage Disease II. This is based on ClinVar entries: low frequency in dbSNP, ClinVar shows: likely loss of function in a gene known to cause this disease, found in multiple cases - mostly homozygous our compound heterozyogtes. Our patient is also heterozygous, so likely asymptomatic. Also in ClinVar in healthy screen. | wgs | WGS_healthyTrunc |
| chr6_7582719_7582719_-_A | DSP | exonic | frameshift insertion | rs1554108609 (this rsID for same protein change, different cDNA change) | 852 | DCM | LIKELEY PATHOGENIC | | clin, wgs | Clin_P, WGS_VUS, WGS_Freq |
| chr3_38606682_38606682_C_T | SCN5A | exonic | nonsynonymous SNV | rs199473101 | 869 | Healthy Control | LIKELEY PATHOGENIC | | wgs | WGS_P |
| chr6_118558957_118558959_AAG_- | PLN | exonic | nonframeshift deletion | rs397516784 | 875 | DCM | PATHOGENIC | CardioClassifier said VUS and only had PM2, PM4. | clin, wgs | Clin_P, WGS_P |
| chr2_178546392_178546392_T_- | TTN | exonic | frameshift deletion | . | 888 | DCM | LIKELEY PATHOGENIC | | wgs | WGS_Freq |
| chr2_178630240_178630240_C_T | TTN | splicing | . | rs771562210 | 910 | DCM | LIKELEY PATHOGENIC | | clin, wgs | Clin_VUS, WGS_P |
| chr7_128841565_128841565_C_T | FLNC | exonic | stopgain | . | 912 | DCM | LIKELEY PATHOGENIC | | clin, wgs | Clin_NotInPanel, WGS_Trunc |
| chr1_156134811_156134811_C_T | LMNA | exonic | nonsynonymous SNV | rs794728591 | 915 | DCM | LIKELEY PATHOGENIC | CardioClassifier said VUS and only had PM2, PP3, PP2 | clin, wgs | Clin_conflict, WGS_P |
| chr1_156136350_156136350_C_T | LMNA | exonic | stopgain | rs267607618 | 920 | DCM | LIKELEY PATHOGENIC | | clin, wgs | Clin_P, WGS_P |
| chr6_7565521_7565521_G_A | DSP | splicing | . | rs727504443 | 925 | DCM | LIKELEY PATHOGENIC | | clin, wgs | Clin_P, WGS_P |
| chr2_178570804_178570804_G_A | TTN | exonic | stopgain | rs794729382 | 928 | DCM | LIKELEY PATHOGENIC | | clin, wgs | Clin_P, WGS_P |
| chr14_23400861_23400861_G_A | MYH6 | exonic | stopgain | rs773659892 | 928 | DCM | UNCERTAIN SIGNIFICANCE | VUS but keep in list. Invitae's ClinVar entry suggests it is likely loss of function in myh6, but not definitive proof for LOF of MYH6 to cause DCM (needed for PVS1), however there are true likely pathogenic loss of function mutations in this gene. Our RNA-seq analysis for control vs DCM shows MYH6 is not changed, nor is MYH6 reduced in patient 928. | clin, wgs | Clin_NotInPanel, WGS_Trunc |
| chr15_63062263_63062263_G_A | TPM1 | exonic | nonsynonymous SNV | rs199476317 | 938 | DCM | LIKELEY PATHOGENIC | CardioClassifier said VUS and only had PM2, PP2 | clin, wgs | Clin_P, WGS_P |
| chr2_178614138_178614138_T_- | TTN | exonic | frameshift deletion | . | 954 | DCM | LIKELEY PATHOGENIC | | wgs | WGS_Freq |
| chr1_109296792_109296792_G_A | MYBPHL | exonic | stopgain | rs777846742 | 954 | DCM | UNCERTAIN SIGNIFICANCE | VUS but keep in list. Not in ClinVar. Not enough data. No clinvar pathogenic / likely pathogenic mutations for this gene, however RNA shows dramatically reduced expression compared to healthy, and even other DCM lines. | wgs | WGS_Trunc |

| | | | | | | | | | | |
|---|---|---|---|---|---|---|---|---|---|---|
| chr2_178585291_178585291_G_A | TTN | exonic | stopgain | rs768345594 | 957 | DCM | LIKELEY PATHOGENIC | | wgs | WGS_P |
| chr7_150948442_150948442_A_C | KCNH2 | splicing | . | . | 959 | Other | LIKELEY PATHOGENIC | | wgs | WGS_healthyFreq |
| chr2_178587173_178587173_A_C | TTN | exonic | stopgain | . | 963 | DCM | LIKELEY PATHOGENIC | | wgs | WGS_Freq |
| chr2_178547445_178547445_G_CTGCTAGA | TTN | exonic | frameshift insertion | rs1219954334 is for g. 178547445_178547446insCTGCTAG -> our is slightly different: Original DNA:44-46:AGG, Original RNA: CCT=Leu. rsID DNA: 44-46: 44=A, 45=G, ins=CTGCTAG, 46=G. rsID RNA: 46=is part of previous codon, then the insertion starts the next codon: CTA = Leu. So there is a frameshift but, the original aa is Leu>Leu -> dbSNP says its Leu>Ser. Regardless, ours is also Leu>Ser. Our DNA: 44=A, ins=CTGCTAGA, 45=-. Our RNA=45:TCT=Ser. So our Mutation is same as rsID except the insertion of CTGCTAG is between 44 and 45 and the G at 46 is replaced with an A -> so ours is g. 178547445delinsCTGCTAGA | 965 | DCM | LIKELEY PATHOGENIC | | clin, wgs | Clin_P, WGS_merge, WGS_Freq |
| chr6_129464371_129464371_C_A | LAMA2 | exonic | stopgain | rs762806915 | 986 | DCM | LIKELEY PATHOGENIC (muscular dystrophy) | for muscular dystrophy | clin, wgs | Clin_P, WGS_P |
| chr6_129454186_129454186_G_T | LAMA2 | exonic | nonsynonymous SNV | . | 986 | DCM | UNCERTAIN SIGNIFICANCE | VUS but keep in list. Only 2 variants in all of gnomAD with a pathogenic / likely pathogenic annotation that are not frameshift, stopgain, startloss, splicing. One is from 2001 and provides no supporting explanation. The other is from 2020 for muscular dystrophy, however it also creates a cryptic splice site. Thus not enough evidence for a missense mutation to cause disease, so VUS. However, the electronic medical record lists the mutation as likely pathogenic. We checked ClinVar for all LAMA2 missense pathogenic / likely pathogenic mutations (mostly muscular). There are 16. 7 lack annotation. For some their call of pathogenic / likely pathogenic don't meet ACMG guidelines. Other ClinVar entries probably do meet the guidelines, but these are all recessive disorders , and they say the variant occurs in trans in patients. Our patient also has a trans pathogenic / likely pathogenic mutation of LAMA2 from the mom. | clin, wgs | clin_research, WGS_VUS |
| chr11_47339792_47339792_T_C | MYBPC3 | splicing | . | rs397515937 | 995 | DCM | PATHOGENIC | | clin, wgs | Clin_P, WGS_P |
| chr10_110812295_110812295_C_T | RBM20 | exonic | nonsynonymous SNV | rs747880281 | 1074 | DCM | LIKELEY PATHOGENIC | CardioClassifier said VUS and only had PM2, PP2. I added PS3 due to (PMID: 32905764) | wgs | WGS_Freq |
| chr2_178564611_178564611_G_- | TTN | exonic | frameshift deletion | . | 2012 | DCM | LIKELEY PATHOGENIC | | clin, wgs | Clin_P, WGS_VUS, WGS_Freq |
| chr6_7565521_7565521_G_T | DSP | splicing | . | (rs727504443 is for G>A; no rsID for G>T) | 2015 | DCM | LIKELEY PATHOGENIC | | clin, wgs | Clin_P, WGS_VUS, WGS_Freq |
| chr2_178587708_178587708_G_A | TTN | exonic | stopgain | rs764243269 | 2025 | DCM | LIKELEY PATHOGENIC | | clin, wgs | Clin_P, WGS_P |
| chr6_112114662_112114662_C_T | LAMA4 | splicing | . | rs368035482 | 2025 | DCM | UNCERTAIN SIGNIFICANCE | VUS but keep in list. If splice site affects expression, still not enough data to link LOF LAMA4 to DCM. We do not have cardiomyocyte RNA-seq data for this sample. | clin, wgs | clin_research, WGS_VUS |
| chr14_23386578_23386578_C_A | MYH6 | exonic | stopgain | . | 2035 | Other | UNCERTAIN SIGNIFICANCE | VUS but keep in list. Variant is in gnomAD but not ClinVar or dbSNP. Low/absent frequency data in gnomAD. Only 1 termination pathogenic / likely pathogenic variant downstream of this one in ClinVar. The entry is sparse so no way to see if claissfication met ACMG guidelines. | wgs | WGS_healthyTrunc |
| chrX_120442650_120442650_G_A | LAMP2 | exonic | stopgain | rs727503118 | 2064 | LVNC | LIKELEY PATHOGENIC | | wgs | WGS_P |
| chr2_178569563_178569563_-_CTT | TTN | exonic | nonframeshift insertion | rs772268958 | 2086 | Healthy Control | LIKELEY PATHOGENIC | | wgs | WGS_healthyFreq |
| chr11_47348424_47348424_C_T | MYBPC3 | | | rs397516074 | 693 (EMR) | HCM | PATHOGENIC (in dad) | The EMR data noted the dad had this mutation. We belive this mutation is pathogenic, however the patient was never tested for it. Our WGS reveals the patient does not actually carry this mutation. This variant is found in 7 patients in our cohort: 3 unrelated HCM patients (319, 431, 582) + 2 related HCM patients (711, 712) + 2 unrelated DCM patients (811, 814). This does not mean this variant wasn't pathogenic in the dad, as 693 could have gotten their pathogenic mutation from their mom. | clin_fam, wgs | clin_research, WGS_AbsentDoubleCheck |

**Table S6**

| sample | Line | CellType | Condition | BioRep | LibraryPreparationBatch | SequencingPool | GC content after filtering (fastp) | % Duplication (fastp) | NUM_UniquelyMappedFrags Millions (STAR) | MeanReads (avg F&R; Millions) (FastQC) | PCT_RIBOSOMAL_BASES (Picard) | PCT_CODEandUTR (Picard) | PCT_UniquelyMappedFrags (STAR) |
|---|---|---|---|---|---|---|---|---|---|---|---|---|---|
| 2003dmso_r1 | 2003 | cm | dmso | r1 | sw10a | sw11b | 50.30% | 47.20% | 62.539685 | 80.954059 | 0.004383 | 0.855332 | 83.64 |
| 2003ips_r1 | 2003 | ips | ips | r1 | sw09 | sw11b | 51.30% | 22.50% | 47.822322 | 66.74269 | 0.064762 | 0.762871 | 75.68 |
| 2003ips_r2 | 2003 | ips | ips | r2 | sw10b | sw11d | 49.10% | 35.40% | 53.858656 | 64.417335 | 0.00288 | 0.847528 | 87.51 |
| 2003myk0.25_r1 | 2003 | cm | myk025 | r1 | sw10b | sw11b | 47.60% | 26.00% | 40.059896 | 49.379751 | 0.009469 | 0.62332 | 84.28 |
| 2003omec0.4_r1 | 2003 | cm | omec04 | r1 | sw10a | sw11b | 49.20% | 59.10% | 44.343124 | 57.360816 | 0.003493 | 0.879318 | 85.2 |
| 2007ips_r1 | 2007 | ips | ips | r1 | sw10a | sw11c | 47.20% | 40.00% | 60.537711 | 69.517315 | 0.006372 | 0.75122 | 89.8 |
| 2007ips_r2 | 2007 | ips | ips | r2 | sw10b | sw11b | 50.30% | 25.20% | 67.45426 | 83.817058 | 0.018556 | 0.810332 | 83.09 |
| 2009dmso_r1 | 2009 | cm | dmso | r1 | sw10b | sw11a | 50.80% | 30.50% | 73.313799 | 92.156584 | 0.018549 | 0.853681 | 81.25 |
| 2009ips_r1 | 2009 | ips | ips | r1 | sw09 | sw11a | 52.90% | 22.40% | 36.351625 | 60.332424 | 0.045297 | 0.767597 | 66.47 |
| 2009myk0.25_r1 | 2009 | cm | myk025 | r1 | sw10b | sw11a | 50.30% | 23.60% | 24.814347 | 30.849938 | 0.016816 | 0.831578 | 82.38 |
| 2009omec0.4_r1 | 2009 | cm | omec04 | r1 | sw10b | sw11a | 50.50% | 27.20% | 64.880825 | 80.118233 | 0.017102 | 0.854572 | 82.4 |
| 2013dmso_r1 | 2013 | cm | dmso | r1 | sw10a | sw11c | 49.90% | 63.70% | 50.519146 | 64.663282 | 0.008304 | 0.859492 | 85.87 |
| 2013ips_r1 | 2013 | ips | ips | r1 | sw10b | sw11c | 49.30% | 21.30% | 34.84084 | 40.381469 | 0.004877 | 0.813107 | 88.53 |
| 2013myk0.25_r1 | 2013 | cm | myk025 | r1 | sw10b | sw11c | 48.40% | 42.50% | 45.594798 | 55.280593 | 0.007964 | 0.826949 | 87.15 |
| 2013omec0.4_r1 | 2013 | cm | omec04 | r1 | sw10a | sw11c | 50.00% | 61.30% | 50.529073 | 61.457531 | 0.002071 | 0.868491 | 87.68 |
| 2015dmso_r1 | 2015 | cm | dmso | r1 | sw09 | sw11b | 51.50% | 20.90% | 52.350078 | 73.270384 | 0.022529 | 0.842932 | 79.01 |
| 2015ips_r1 | 2015 | ips | ips | r1 | sw09 | sw11b | 51.20% | 23.10% | 39.427773 | 55.564992 | 0.029137 | 0.824275 | 77.9 |
| 2015myk0.25_r1 | 2015 | cm | myk025 | r1 | sw09 | sw11b | 49.90% | 23.00% | 43.468233 | 58.148526 | 0.020999 | 0.834551 | 82.31 |
| 2015omec0.4_r1 | 2015 | cm | omec04 | r1 | sw09 | sw11b | 50.40% | 35.80% | 89.048686 | 122.102495 | 0.025648 | 0.855753 | 81.6 |
| 2020Ddmso_r1 | 2020 | cm | dmso | r1 | sw12-BB | sw12-BB | 50.60% | 31.00% | 54.006279 | 64.025656 | 0.011046 | 0.854338 | 85.25 |
| 2020Domec1.0_r1 | 2020 | cm | omec10 | r1 | sw12-BB | sw12-BB | 51.00% | 32.70% | 54.779448 | 65.41854 | 0.009552 | 0.865048 | 84.95 |
| 2025ips_r1 | 2025 | ips | ips | r1 | sw12-AA | sw12-AA | 50.20% | 39.40% | 38.084511 | 72.710677 | 0.15974 | 0.685593 | 56.43 |
| 2036dmso_r1 | 2036 | cm | dmso | r1 | sw10b | sw11b | 49.20% | 25.90% | 39.517192 | 46.433689 | 0.010067 | 0.839099 | 87.17 |
| 2036dmso_r2 | 2036 | cm | dmso | r2 | sw10bmanual | sw11a | 48.30% | 21.80% | 62.137608 | 70.815811 | 0.00182 | 0.864781 | 88.43 |
| 2036ips_r1 | 2036 | ips | ips | r1 | sw09 | sw11b | 50.60% | 19.30% | 35.066249 | 47.832428 | 0.016378 | 0.848225 | 80.57 |
| 2036ips_r2 | 2036 | ips | ips | r2 | sw10b | sw11a | 49.70% | 28.10% | 73.197143 | 86.613604 | 0.007512 | 0.857317 | 86.28 |
| 2036myk0.25_r1 | 2036 | cm | myk025 | r1 | sw10b | sw11b | 49.70% | 29.00% | 42.179178 | 51.28433 | 0.011651 | 0.834578 | 84.82 |
| 2036myk0.25_r2 | 2036 | cm | myk025 | r2 | sw10b | sw11a | 50.00% | 27.40% | 46.615421 | 57.104626 | 0.016198 | 0.834835 | 83.93 |
| 2036omec0.4_r1 | 2036 | cm | omec04 | r1 | sw10b | sw11b | 49.90% | 35.90% | 62.286681 | 76.586923 | 0.01263 | 0.836841 | 84.02 |
| 2036omec0.4_r2 | 2036 | cm | omec04 | r2 | sw10b | sw11a | 49.10% | 40.60% | 74.305653 | 89.131378 | 0.01175 | 0.8627 | 85.4 |
| 2086ips_r1 | 2086 | ips | ips | r1 | sw12-AA | sw12-AA | 48.70% | 37.60% | 41.419996 | 78.297212 | 0.201685 | 0.661913 | 55.4 |
| 283dmso_r1 | 283 | cm | dmso | r1 | sw04 | sw11c | 52.40% | 28.20% | 39.624098 | 65.331468 | 0.088848 | 0.738456 | 61.63 |
| 283ips_r1 | 283 | ips | ips | r1 | sw04 | sw11c | 51.60% | 32.90% | 65.511867 | 103.254057 | 0.067462 | 0.738614 | 64.79 |
| 284dmso_r1 | 284 | cm | dmso | r1 | sw03 | sw11a | 51.30% | 36.20% | 61.460192 | 95.375744 | 0.05695 | 0.781501 | 69.05 |
| 284dmso_r2 | 284 | cm | dmso | r2 | sw04 | sw11b | 50.10% | 28.00% | 40.96328 | 54.059724 | 0.040327 | 0.806483 | 77.75 |
| 284ips_r1 | 284 | ips | ips | r1 | sw03 | sw11a | 53.70% | 44.90% | 59.44872 | 149.883103 | 0.122667 | 0.623248 | 42.54 |
| 287dmso_r1 | 287 | cm | dmso | r1 | sw03 | sw11a | 51.90% | 38.40% | 90.02388 | 133.359522 | 0.057786 | 0.800301 | 71.53 |
| 287dmso_r2 | 287 | cm | dmso | r2 | sw04 | sw11c | 50.60% | 38.00% | 46.940733 | 61.157297 | 0.030598 | 0.830709 | 78.41 |
| 287ips_r1 | 287 | ips | ips | r1 | sw03 | sw11a | 50.30% | 32.80% | 85.555972 | 116.255419 | 0.030259 | 0.823999 | 77.96 |
| 289dmso_r1 | 289 | cm | dmso | r1 | sw03 | sw11b | 50.90% | 38.30% | 93.636816 | 124.996919 | 0.044242 | 0.816507 | 78.68 |
| 289dmso_r2 | 289 | cm | dmso | r2 | sw04 | sw11c | 51.50% | 35.10% | 49.902509 | 72.561791 | 0.054785 | 0.799939 | 70.07 |
| 289ips_r1 | 289 | ips | ips | r1 | sw03 | sw11b | 52.10% | 29.30% | 59.487682 | 93.291105 | 0.05987 | 0.774857 | 66.81 |
| 295dmso_r1 | 295 | cm | dmso | r1 | sw04 | sw11a | 51.40% | 33.30% | 56.398087 | 79.802103 | 0.056169 | 0.784269 | 72.26 |
| 295ips_r1 | 295 | ips | ips | r1 | sw04 | sw11a | 52.00% | 26.30% | 43.045158 | 67.722671 | 0.058782 | 0.766432 | 64.86 |
| 297dmso_r1 | 297 | cm | dmso | r1 | sw03 | sw11a | 52.60% | 28.20% | 32.635166 | 55.095821 | 0.099667 | 0.742694 | 63.49 |
| 297dmso_r2 | 297 | cm | dmso | r2 | sw04 | sw11b | 51.60% | 29.70% | 38.698999 | 57.203194 | 0.055539 | 0.788542 | 69.48 |
| 297ips_r1 | 297 | ips | ips | r1 | sw03 | sw11a | 51.70% | 31.70% | 60.270551 | 91.095804 | 0.057723 | 0.789897 | 71.43 |
| 298dmso_r1 | 298 | cm | dmso | r1 | sw04 | sw11c | 53.00% | 29.80% | 31.55685 | 62.190031 | 0.093077 | 0.704291 | 51.86 |
| 298ips_r1 | 298 | ips | ips | r1 | sw04 | sw11c | 53.10% | 26.40% | 37.002649 | 75.799271 | 0.104211 | 0.671333 | 49.88 |
| 301dmso_r1 | 301 | cm | dmso | r1 | sw03 | sw11d | 52.20% | 41.50% | 67.540606 | 103.498162 | 0.069675 | 0.772407 | 68.89 |
| 301ips_r1 | 301 | ips | ips | r1 | sw03 | sw11d | 50.90% | 27.60% | 47.095755 | 63.014751 | 0.02184 | 0.83812 | 79.17 |
| 304ips_r1 | 304 | ips | ips | r1 | sw03 | sw11b | 54.10% | 31.90% | 59.246483 | 156.540622 | 0.119987 | 0.624168 | 40.38 |
| 310dmso_r1 | 310 | cm | dmso | r1 | sw03 | sw11a | 53.10% | 45.20% | 67.190036 | 138.696469 | 0.091669 | 0.689663 | 51.52 |
| 310ips_r1 | 310 | ips | ips | r1 | sw03 | sw11a | 51.50% | 28.30% | 41.911878 | 60.399023 | 0.036176 | 0.82359 | 73.84 |
| 319dmso_r1 | 319 | cm | dmso | r1 | sw04 | sw11b | 53.80% | 29.80% | 40.703943 | 90.02978 | 0.111876 | 0.658975 | 46.09 |
| 319ips_r1 | 319 | ips | ips | r1 | sw04 | sw11b | 51.20% | 25.20% | 42.700183 | 59.561314 | 0.038825 | 0.814026 | 72.79 |
| 320dmso_r1 | 320 | cm | dmso | r1 | sw03 | sw11c | 52.60% | 34.20% | 30.957048 | 49.701566 | 0.062735 | 0.799358 | 66.26 |
| 320dmso_r2 | 320 | cm | dmso | r2 | sw04 | sw11a | 52.60% | 34.40% | 39.508717 | 65.702912 | 0.067618 | 0.764093 | 61.15 |
| 320ips_r1 | 320 | ips | ips | r1 | sw03 | sw11c | 51.40% | 34.70% | 75.695147 | 108.199514 | 0.034507 | 0.818881 | 74.87 |
| 333dmso_r1 | 333 | cm | dmso | r1 | sw04 | sw11c | 52.20% | 32.50% | 54.70965 | 80.357938 | 0.059074 | 0.804908 | 69.15 |
| 333ips_r1 | 333 | ips | ips | r1 | sw04 | sw11c | 51.90% | 27.00% | 46.310498 | 65.18917 | 0.066488 | 0.780417 | 72.27 |
| 334dmso_r1 | 334 | cm | dmso | r1 | sw04 | sw11d | 51.20% | 39.10% | 61.500067 | 87.935801 | 0.050465 | 0.805965 | 72.46 |
| 334dmso_r2 | 334 | cm | dmso | r2 | sw04 | sw11c | 53.10% | 28.00% | 29.147052 | 48.069909 | 0.071336 | 0.765471 | 61.73 |
| 334ips_r1 | 334 | ips | ips | r1 | sw04 | sw11d | 50.60% | 31.40% | 53.497811 | 73.99456 | 0.037995 | 0.810966 | 73.92 |
| 338dmso_r1 | 338 | cm | dmso | r1 | sw04 | sw11b | 53.30% | 28.10% | 39.25627 | 72.235323 | 0.089826 | 0.719175 | 55.17 |
| 338ips_r1 | 338 | ips | ips | r1 | sw04 | sw11b | 54.10% | 23.00% | 40.593361 | 101.020816 | 0.121414 | 0.608461 | 40.73 |
| 352ips_r1 | 352 | ips | ips | r1 | sw03 | sw11a | 53.30% | 46.40% | 59.650446 | 128.993577 | 0.082502 | 0.706626 | 48.94 |
| 356dmso_r1 | 356 | cm | dmso | r1 | sw04 | sw11a | 52.40% | 33.90% | 44.522567 | 69.713542 | 0.071806 | 0.760535 | 65.1 |
| 356dmso_r2 | 356 | cm | dmso | r2 | sw04 | sw11b | 53.00% | 30.40% | 38.208166 | 66.32143 | 0.088122 | 0.732118 | 58.59 |
| 356ips_r1 | 356 | ips | ips | r1 | sw04 | sw11a | 50.20% | 27.50% | 35.83098 | 47.372639 | 0.030711 | 0.82848 | 77.56 |
| 358dmso_r1 | 358 | cm | dmso | r1 | sw04 | sw11b | 51.60% | 31.10% | 52.487452 | 70.439312 | 0.042917 | 0.830775 | 75.62 |
| 358ips_r1 | 358 | ips | ips | r1 | sw04 | sw11b | 53.20% | 28.20% | 43.67645 | 83.162462 | 0.09112 | 0.707646 | 53.26 |

| | | | | | | | | | | | | | |
|---|---|---|---|---|---|---|---|---|---|---|---|---|---|
| 367dmso_r1 | 367 | cm | dmso | r1 | sw03 | sw11d | 52.90% | 34.20% | 28.163355 | 58.887721 | 0.119321 | 0.670764 | 50.73 |
| 367ips_r1 | 367 | ips | ips | r1 | sw03 | sw11d | 54.60% | 42.40% | 53.838975 | 191.358279 | 0.139785 | 0.551802 | 29.79 |
| 371dmso_r1 | 371 | cm | dmso | r1 | sw04 | sw11c | 52.60% | 32.20% | 46.938589 | 80.665788 | 0.083321 | 0.730294 | 59.8 |
| 371ips_r1 | 371 | ips | ips | r1 | sw04 | sw11c | 51.70% | 31.60% | 66.289616 | 89.732929 | 0.047652 | 0.798245 | 75.2 |
| 372dmso_r1 | 372 | cm | dmso | r1 | sw03 | sw11a | 53.70% | 37.10% | 52.517192 | 128.149167 | 0.119639 | 0.650692 | 44.41 |
| 372ips_r1 | 372 | ips | ips | r1 | sw03 | sw11a | 53.60% | 34.90% | 47.497213 | 108.277695 | 0.09955 | 0.680587 | 46.83 |
| 373dmso_r1 | 373 | cm | dmso | r1 | sw06 | sw11d | 51.70% | 25.50% | 38.6952 | 61.705578 | 0.058152 | 0.799592 | 68.69 |
| 373ips_r1 | 373 | ips | ips | r1 | sw06 | sw11d | 49.60% | 30.90% | 58.602372 | 86.975544 | 0.020656 | 0.833941 | 78.51 |
| 374dmso_r1 | 374 | cm | dmso | r1 | sw04 | sw11b | 54.20% | 27.00% | 37.985368 | 98.999452 | 0.129237 | 0.603749 | 39.23 |
| 374ips_r1 | 374 | ips | ips | r1 | sw04 | sw11b | 53.40% | 28.20% | 38.383614 | 80.851846 | 0.10552 | 0.672139 | 48.18 |
| 375dmso_r1 | 375 | cm | dmso | r1 | sw04 | sw11c | 51.60% | 33.90% | 39.139844 | 54.160311 | 0.040856 | 0.819112 | 73.76 |
| 375ips_r1 | 375 | ips | ips | r1 | sw04 | sw11c | 51.90% | 28.90% | 50.556293 | 77.624717 | 0.051299 | 0.791981 | 66.72 |
| 376dmso_r1 | 376 | cm | dmso | r1 | sw04 | sw11d | 52.10% | 30.90% | 52.1962 | 82.946384 | 0.088619 | 0.737838 | 64.24 |
| 376ips_r1 | 376 | ips | ips | r1 | sw04 | sw11d | 52.10% | 30.60% | 32.834101 | 55.721056 | 0.071644 | 0.74772 | 60.21 |
| 378dmso_r1 | 378 | cm | dmso | r1 | sw04 | sw11b | 52.30% | 33.10% | 60.017067 | 90.047421 | 0.071891 | 0.773369 | 67.77 |
| 378ips_r1 | 378 | ips | ips | r1 | sw04 | sw11b | 51.60% | 27.30% | 41.038545 | 61.729363 | 0.053462 | 0.779867 | 67.87 |
| 380dmso_r1 | 380 | cm | dmso | r1 | sw06 | sw11b | 50.20% | 51.40% | 128.536329 | 195.402653 | 0.031436 | 0.817305 | 73.86 |
| 380ips_r1 | 380 | ips | ips | r1 | sw06 | sw11b | 49.00% | 37.30% | 96.312896 | 133.091575 | 0.017598 | 0.843316 | 81.11 |
| 388dmso_r1 | 388 | cm | dmso | r1 | sw06 | sw11a | 49.70% | 32.00% | 67.625963 | 92.226701 | 0.023153 | 0.849766 | 79.54 |
| 388dmso_r2 | 388 | cm | dmso | r2 | sw08 | sw11b | 52.60% | 21.90% | 26.842013 | 49.699616 | 0.080041 | 0.742375 | 57.4 |
| 388ips_r1 | 388 | ips | ips | r1 | sw06 | sw11a | 49.90% | 28.50% | 63.187967 | 91.149223 | 0.016353 | 0.835423 | 79.11 |
| 390dmso_r1 | 390 | cm | dmso | r1 | sw06 | sw11d | 50.00% | 33.20% | 87.159283 | 118.426798 | 0.035883 | 0.83279 | 79.32 |
| 390ips_r1 | 390 | ips | ips | r1 | sw06 | sw11d | 50.00% | 18.00% | 75.864141 | 100.656236 | 0.058775 | 0.788405 | 79.31 |
| 393ips_r1 | 393 | ips | ips | r1 | sw05 | sw11c | 50.30% | 31.70% | 81.571366 | 124.040227 | 0.054221 | 0.783393 | 70 |
| 394dmso_r1 | 394 | cm | dmso | r1 | sw05 | sw11c | 50.90% | 27.90% | 50.509828 | 72.051622 | 0.044133 | 0.823439 | 72.38 |
| 394ips_r1 | 394 | ips | ips | r1 | sw05 | sw11c | 51.30% | 27.80% | 60.426198 | 94.916308 | 0.066257 | 0.765233 | 65.7 |
| 395dmso_r1 | 395 | cm | dmso | r1 | sw07 | sw11d | 50.10% | 47.90% | 54.393613 | 80.594271 | 0.022622 | 0.850672 | 75.6 |
| 395ips_r1 | 395 | ips | ips | r1 | sw07 | sw11d | 49.60% | 27.50% | 54.404872 | 70.484744 | 0.020155 | 0.83477 | 81.22 |
| 397dmso_r1 | 397 | cm | dmso | r1 | sw03 | sw11b | 52.90% | 37.80% | 51.06809 | 79.503906 | 0.059192 | 0.798126 | 69.1 |
| 397dmso_r2 | 397 | cm | dmso | r2 | sw04 | sw11a | 52.50% | 35.50% | 45.114875 | 69.344002 | 0.058653 | 0.788031 | 66.74 |
| 397ips_r1 | 397 | ips | ips | r1 | sw03 | sw11b | 51.00% | 25.50% | 37.242936 | 56.880394 | 0.044544 | 0.789663 | 70.04 |
| 398dmso_r1 | 398 | cm | dmso | r1 | sw05 | sw11a | 53.10% | 26.70% | 43.536233 | 109.758084 | 0.126061 | 0.654522 | 42.06 |
| 398ips_r1 | 398 | ips | ips | r1 | sw05 | sw11a | 50.80% | 24.80% | 49.34187 | 85.450078 | 0.075067 | 0.754508 | 62.31 |
| 405dmso_r1 | 405 | cm | dmso | r1 | sw07 | sw11b | 50.90% | 33.40% | 67.071215 | 100.703214 | 0.055046 | 0.803622 | 70.48 |
| 411dmso_r1 | 411 | cm | dmso | r1 | sw07 | sw11c | 49.80% | 40.80% | 86.348934 | 122.293113 | 0.039756 | 0.836257 | 76.21 |
| 413dmso_r1 | 413 | cm | dmso | r1 | sw08 | sw11b | 51.00% | 22.70% | 20.770538 | 36.164441 | 0.076734 | 0.754531 | 61.82 |
| 419dmso_r2 | 419 | cm | dmso | r2 | sw08 | sw11d | 49.90% | 25.10% | 39.289612 | 50.061575 | 0.026898 | 0.86103 | 80.57 |
| 419ips_r1 | 419 | ips | ips | r1 | sw10a | sw11c | 49.50% | 23.60% | 48.207609 | 59.560232 | 0.002099 | 0.867365 | 85.06 |
| 420dmso_r1 | 420 | cm | dmso | r1 | sw10b | sw11a | 48.90% | 26.80% | 43.993202 | 53.272446 | 0.019154 | 0.82869 | 84.82 |
| 421dmso_r1 | 421 | cm | dmso | r1 | sw09 | sw11a | 51.60% | 37.70% | 78.940508 | 113.144988 | 0.014046 | 0.839711 | 77.63 |
| 428dmso_r1 | 428 | cm | dmso | r1 | sw08 | sw11d | 51.90% | 32.80% | 48.2061 | 95.066867 | 0.093754 | 0.716044 | 54.91 |
| 428ips_r1 | 428 | ips | ips | r1 | sw12-AA | sw12-AA | 51.30% | 17.00% | 25.434175 | 93.390521 | 0.26805 | 0.53012 | 28.05 |
| 431Homec0.4_r1 | 431 | cm | omec04 | r1 | sw12-AA | sw12-AA | 49.40% | 44.20% | 46.741396 | 103.399218 | 0.190298 | 0.642301 | 47.4 |
| 431ips_r1 | 431 | ips | ips | r1 | sw12-AA | sw12-AA | 50.30% | 29.40% | 45.572884 | 104.978658 | 0.212524 | 0.613006 | 44.8 |
| 437dmso_r1 | 437 | cm | dmso | r1 | sw08 | sw11a | 51.70% | 28.20% | 33.001629 | 54.621922 | 0.074231 | 0.762406 | 63.18 |
| 438dmso_r1 | 438 | cm | dmso | r1 | sw08 | sw11d | 49.60% | 38.10% | 78.572741 | 108.808108 | 0.033046 | 0.843786 | 77.26 |
| 440dmso_r1 | 440 | cm | dmso | r1 | sw10a | sw11b | 50.30% | 60.00% | 31.115053 | 41.25575 | 0.003434 | 0.871343 | 84.58 |
| 440ips_r1 | 440 | ips | ips | r1 | sw10a | sw11b | 50.10% | 27.80% | 79.317391 | 96.111169 | 0.003692 | 0.864944 | 85.14 |
| 440myk0.25_r1 | 440 | cm | myk025 | r1 | sw10a | sw11b | 49.80% | 40.70% | 50.931403 | 61.486891 | 0.005754 | 0.871124 | 86.26 |
| 440omec0.4_r1 | 440 | cm | omec04 | r1 | sw10a | sw11b | 49.00% | 63.00% | 34.135661 | 42.766511 | 0.002885 | 0.868785 | 87.42 |
| 494dmso_r1 | 494 | cm | dmso | r1 | sw08 | sw11a | 50.60% | 43.10% | 47.559835 | 68.120095 | 0.031797 | 0.79576 | 75.31 |
| 501dmso_r1 | 501 | cm | dmso | r1 | sw08 | sw11d | 51.00% | 22.50% | 26.40619 | 40.645982 | 0.070853 | 0.770939 | 67.37 |
| 502dmso_r1 | 502 | cm | dmso | r1 | sw10b | sw11b | 48.80% | 28.40% | 55.924287 | 67.84945 | 0.011875 | 0.855639 | 86.73 |
| 502myk0.25_r1 | 502 | cm | myk025 | r1 | sw09 | sw11b | 50.70% | 26.20% | 48.256469 | 65.284297 | 0.020799 | 0.812379 | 80.14 |
| 503dmso_r1 | 503 | cm | dmso | r1 | sw08 | sw11c | 51.30% | 25.20% | 50.988582 | 70.056707 | 0.047292 | 0.814188 | 74.18 |
| 512Cdmso_r1 | 512 | cm | dmso | r1 | sw12-BB | sw12-BB | 52.00% | 40.20% | 45.007579 | 62.241008 | 0.011831 | 0.839438 | 79.77 |
| 512Cmyk0.25_r1 | 512 | cm | myk025 | r1 | sw12-BB | sw12-BB | 51.20% | 36.20% | 37.770781 | 52.632229 | 0.011554 | 0.840044 | 79.66 |
| 512Comec0.4_r1 | 512 | cm | omec04 | r1 | sw12-BB | sw12-BB | 51.80% | 37.40% | 52.757757 | 72.914944 | 0.011136 | 0.835126 | 80.09 |
| 512Comec1.0_r1 | 512 | cm | omec10 | r1 | sw12-BB | sw12-BB | 51.90% | 37.40% | 46.740581 | 65.661936 | 0.010185 | 0.842742 | 79.2 |
| 512ips_r1 | 512 | ips | ips | r1 | sw12-BB | sw12-BB | 48.70% | 22.80% | 63.176205 | 79.918533 | 0.007798 | 0.833914 | 85.48 |
| 517dmso_r1 | 517 | cm | dmso | r1 | sw10bmanual | sw11c | 48.10% | 25.00% | 52.632335 | 62.163442 | 0.001135 | 0.867869 | 86.79 |
| 519dmso_r1 | 519 | cm | dmso | r1 | sw08 | sw11a | 51.20% | 26.30% | 35.627269 | 51.882418 | 0.049129 | 0.81552 | 70.71 |
| 520dmso_r1 | 520 | cm | dmso | r1 | sw09 | sw11b | 50.40% | 29.90% | 59.745383 | 82.528121 | 0.014366 | 0.845373 | 83.65 |
| 521dmso_r1 | 521 | cm | dmso | r1 | sw09 | sw11d | 50.50% | 23.00% | 42.111345 | 57.505047 | 0.030442 | 0.849502 | 81.88 |
| 526dmso_r1 | 526 | cm | dmso | r1 | sw09 | sw11d | 48.60% | 28.50% | 47.625225 | 61.491731 | 0.016692 | 0.857721 | 86.49 |
| 530dmso_r1 | 530 | cm | dmso | r1 | sw09 | sw11d | 50.40% | 27.00% | 43.669732 | 59.033266 | 0.013949 | 0.870309 | 84.16 |
| 532dmso_r1 | 532 | cm | dmso | r1 | sw09 | sw11a | 50.70% | 39.40% | 44.9205 | 63.991214 | 0.015061 | 0.857128 | 81.97 |
| 532ips_r1 | 532 | ips | ips | r1 | sw09 | sw11a | 51.60% | 21.60% | 45.018209 | 66.001389 | 0.028023 | 0.815989 | 74.35 |
| 534dmso_r1 | 534 | cm | dmso | r1 | sw10b | sw11a | 49.60% | 48.00% | 126.105336 | 153.222461 | 0.016404 | 0.836101 | 84.11 |
| 534ips_r1 | 534 | ips | ips | r1 | sw09 | sw11a | 50.20% | 18.60% | 30.281721 | 40.452215 | 0.012592 | 0.826827 | 84.01 |
| 534myk0.25_r1 | 534 | cm | myk025 | r1 | sw10a | sw11a | 50.20% | 32.30% | 27.120236 | 35.340002 | 0.006922 | 0.84984 | 83.94 |
| 534omec0.4_r1 | 534 | cm | omec04 | r1 | sw10a | sw11a | 46.50% | 62.10% | 34.167518 | 41.255763 | 0.002669 | 0.813565 | 87.41 |
| 535dmso_r1 | 535 | cm | dmso | r1 | sw10b | sw11a | 48.80% | 49.50% | 87.555292 | 101.972175 | 0.010362 | 0.861645 | 88 |
| 535ips_r1 | 535 | ips | ips | r1 | sw09 | sw11a | 50.90% | 18.90% | 36.708673 | 49.233802 | 0.025611 | 0.819317 | 81.81 |
| 541dmso_r1 | 541 | cm | dmso | r1 | sw10a | sw11c | 49.90% | 39.30% | 54.732014 | 65.648591 | 0.004326 | 0.880368 | 86.5 |

| | | | | | | | | | | | | | |
|---|---|---|---|---|---|---|---|---|---|---|---|---|---|
| 541ips_r1 | 541 | ips | ips | r1 | sw09 | sw11b | 51.10% | 18.60% | 32.72826 | 46.52303 | 0.062712 | 0.760553 | 76.97 |
| 541myk0.25_r1 | 541 | cm | myk025 | r1 | sw10a | sw11c | 48.70% | 41.30% | 53.999729 | 64.217599 | 0.00493 | 0.856871 | 87.33 |
| 541omec0.4_r1 | 541 | cm | omec04 | r1 | sw10a | sw11c | 50.10% | 42.90% | 61.613321 | 78.460586 | 0.00461 | 0.882528 | 84.15 |
| 543ips_r1 | 543 | ips | ips | r1 | sw10b | sw11d | 49.90% | 25.10% | 52.656225 | 63.127062 | 0.007527 | 0.82004 | 86.76 |
| 544dmso_r1 | 544 | cm | dmso | r1 | sw10b | sw11b | 49.90% | 21.90% | 25.176286 | 29.859023 | 0.009136 | 0.845949 | 86.28 |
| 544ips_r1 | 544 | ips | ips | r1 | sw09 | sw11b | 50.70% | 18.90% | 42.837937 | 60.11345 | 0.028529 | 0.82116 | 78.34 |
| 546dmso_r1 | 546 | cm | dmso | r1 | sw09 | sw11a | 50.60% | 21.40% | 25.516424 | 34.971036 | 0.024423 | 0.830308 | 80.78 |
| 546ips_r1 | 546 | ips | ips | r1 | sw10a | sw11a | 49.10% | 25.80% | 50.939129 | 62.223259 | 0.003435 | 0.781164 | 86.14 |
| 546myk0.25_r1 | 546 | cm | myk025 | r1 | sw09 | sw11a | 52.60% | 18.30% | 29.270184 | 41.485296 | 0.02445 | 0.834968 | 77.35 |
| 555dmso_r1 | 555 | cm | dmso | r1 | sw09 | sw11a | 50.90% | 23.80% | 27.63253 | 37.371959 | 0.020938 | 0.847212 | 81.97 |
| 555myk0.25_r1 | 555 | cm | myk025 | r1 | sw09 | sw11a | 50.40% | 23.00% | 35.737268 | 49.17088 | 0.031611 | 0.829773 | 80.72 |
| 557dmso_r1 | 557 | cm | dmso | r1 | sw08 | sw11b | 49.80% | 26.80% | 33.799592 | 45.995357 | 0.025907 | 0.855729 | 78.79 |
| 562Ddmso_r1 | 562 | cm | dmso | r1 | sw12-AA | sw12-AA | 49.00% | 43.10% | 50.767605 | 102.437999 | 0.190285 | 0.646949 | 51.18 |
| 562Dmyk0.25_r1 | 562 | cm | myk025 | r1 | sw12-AA | sw12-AA | 49.30% | 48.30% | 46.13347 | 97.764979 | 0.198917 | 0.636977 | 50.14 |
| 562ips_r1 | 562 | ips | ips | r1 | sw12-AA | sw12-AA | 47.80% | 21.70% | 31.0119 | 51.981234 | 0.156955 | 0.702394 | 60.88 |
| 565dmso_r1 | 565 | cm | dmso | r1 | sw10b | sw11a | 50.10% | 28.60% | 62.173914 | 73.383608 | 0.008692 | 0.858112 | 86.82 |
| 565dmso_r2 | 565 | cm | dmso | r2 | sw10a | sw11b | 50.50% | 49.70% | 85.555552 | 101.909845 | 0.004164 | 0.890591 | 87.8 |
| 565ips_r1 | 565 | ips | ips | r1 | sw10a | sw11a | 49.90% | 41.70% | 100.105707 | 117.387169 | 0.002802 | 0.874573 | 87.57 |
| 565myk0.25_r1 | 565 | cm | myk025 | r1 | sw10b | sw11a | 49.60% | 22.70% | 37.803552 | 44.579015 | 0.013259 | 0.845663 | 86.79 |
| 565myk0.25_r2 | 565 | cm | myk025 | r2 | sw10a | sw11b | 50.20% | 55.40% | 64.729978 | 85.178723 | 0.005122 | 0.883482 | 85.22 |
| 565omec0.4_r1 | 565 | cm | omec04 | r1 | sw10b | sw11a | 49.60% | 25.50% | 44.07123 | 52.237138 | 0.01029 | 0.859102 | 86.48 |
| 565omec0.4_r2 | 565 | cm | omec04 | r2 | sw10a | sw11b | 48.10% | 60.20% | 44.362215 | 55.024207 | 0.002481 | 0.87549 | 88.08 |
| 570Bomec1.0_r1 | 570 | cm | omec10 | r1 | sw12-BB | sw12-BB | 51.10% | 25.90% | 50.629704 | 65.690784 | 0.018585 | 0.853586 | 81.93 |
| 570ips_r1 | 570 | ips | ips | r1 | sw12-BB | sw12-BB | 50.10% | 43.60% | 44.163385 | 58.37553 | 0.008779 | 0.86177 | 83.06 |
| 571Fdmso_r1 | 571 | cm | dmso | r1 | sw12-BB | sw12-BB | 51.00% | 29.90% | 45.240831 | 60.115964 | 0.011558 | 0.851572 | 81.48 |
| 571Fmyk0.25_r1 | 571 | cm | myk025 | r1 | sw12-BB | sw12-BB | 50.80% | 30.30% | 37.371488 | 49.945089 | 0.012715 | 0.847622 | 81.62 |
| 571Fomec0.4_r1 | 571 | cm | omec04 | r1 | sw12-BB | sw12-BB | 51.80% | 36.40% | 47.978338 | 66.200262 | 0.011458 | 0.834941 | 79.46 |
| 571Fomec1.0_r1 | 571 | cm | omec10 | r1 | sw12-BB | sw12-BB | 52.90% | 34.10% | 39.225795 | 66.635126 | 0.048517 | 0.764721 | 64.31 |
| 571ips_r1 | 571 | ips | ips | r1 | sw12-BB | sw12-BB | 50.40% | 20.70% | 52.778428 | 68.348825 | 0.013743 | 0.857037 | 81.86 |
| 582ips_r1 | 582 | ips | ips | r1 | sw12-AA | sw12-AA | 49.50% | 36.40% | 34.916267 | 69.059223 | 0.214813 | 0.64314 | 53.38 |
| 592ips_r1 | 592 | ips | ips | r1 | sw10a | sw11c | 47.70% | 51.50% | 50.857681 | 63.584195 | 0.001719 | 0.867801 | 87.21 |
| 603dmso_r1 | 603 | cm | dmso | r1 | sw10a | sw11c | 50.60% | 42.80% | 67.382517 | 83.260351 | 0.005815 | 0.881403 | 85.04 |
| 603dmso_r2 | 603 | cm | dmso | r2 | sw10b | sw11d | 50.20% | 24.20% | 37.429957 | 44.231441 | 0.009775 | 0.868781 | 86.79 |
| 603ips_r1 | 603 | ips | ips | r1 | sw10a | sw11c | 49.40% | 54.40% | 48.233935 | 60.121835 | 0.001591 | 0.870173 | 87.76 |
| 603myk0.25_r1 | 603 | cm | myk025 | r1 | sw10a | sw11c | 50.30% | 41.40% | 59.845099 | 74.353345 | 0.004481 | 0.83437 | 84.46 |
| 603myk0.25_r2 | 603 | cm | myk025 | r2 | sw10b | sw11d | 49.00% | 24.40% | 22.507152 | 26.523945 | 0.009535 | 0.85641 | 87.26 |
| 603omec0.4_r1 | 603 | cm | omec04 | r1 | sw10a | sw11c | 51.00% | 49.80% | 74.275049 | 91.873661 | 0.004521 | 0.869075 | 84.22 |
| 603omec0.4_r2 | 603 | cm | omec04 | r2 | sw10b | sw11d | 49.80% | 37.80% | 75.02549 | 89.447677 | 0.012338 | 0.871286 | 85.57 |
| 612ips_r1 | 612 | ips | ips | r1 | sw12-AA | sw12-AA | 50.00% | 26.70% | 31.90106 | 76.291386 | 0.240166 | 0.596948 | 43.13 |
| 622ips_r1 | 622 | ips | ips | r1 | sw10a | sw11d | 49.10% | 41.80% | 50.080269 | 62.341055 | 0.002786 | 0.87297 | 85.81 |
| 653dmso_r1 | 653 | cm | dmso | r1 | sw10a | sw11d | 50.00% | 44.50% | 59.550346 | 73.636846 | 0.003869 | 0.891043 | 86.8 |
| 653ips_r1 | 653 | ips | ips | r1 | sw10a | sw11d | 48.70% | 47.70% | 73.587016 | 88.116465 | 0.001924 | 0.875705 | 87.92 |
| 653myk0.25_r1 | 653 | cm | myk025 | r1 | sw10b | sw11d | 50.60% | 57.50% | 144.648086 | 184.66818 | 0.023155 | 0.849798 | 80.41 |
| 653omec0.4_r1 | 653 | cm | omec04 | r1 | sw10a | sw11d | 51.90% | 71.20% | 35.693054 | 63.777759 | 0.016377 | 0.716322 | 62.65 |
| 662Eomec0.4_r1 | 662 | cm | omec04 | r1 | sw12-BB | sw12-BB | 53.00% | 26.60% | 37.123606 | 54.589637 | 0.063008 | 0.731516 | 73.29 |
| 662Eomec1.0_r1 | 662 | cm | omec10 | r1 | sw12-BB | sw12-BB | 52.20% | 31.70% | 48.890671 | 66.170329 | 0.015678 | 0.870128 | 80.05 |
| 662ips_r1 | 662 | ips | ips | r1 | sw12-BB | sw12-BB | 50.10% | 27.00% | 28.470425 | 72.439237 | 0.211392 | 0.586425 | 40.87 |
| 673ips_r1 | 673 | ips | ips | r1 | sw12-AA | sw12-AA | 49.70% | 30.00% | 39.490705 | 72.406605 | 0.184919 | 0.678696 | 58.33 |
| 697Ddmso_r1 | 697 | cm | dmso | r1 | sw12-BB | sw12-BB | 51.10% | 31.70% | 47.150725 | 61.639022 | 0.01032 | 0.862568 | 83.16 |
| 697Dmyk0.25_r1 | 697 | cm | myk025 | r1 | sw12-BB | sw12-BB | 51.10% | 36.40% | 54.441006 | 72.402251 | 0.013117 | 0.855645 | 80.34 |
| 697Domec0.4_r1 | 697 | cm | omec04 | r1 | sw12-BB | sw12-BB | 50.80% | 44.20% | 50.815347 | 69.067355 | 0.014405 | 0.848599 | 79.96 |
| 697ips_r1 | 697 | ips | ips | r1 | sw12-BB | sw12-BB | 48.90% | 19.20% | 65.390065 | 81.268318 | 0.008277 | 0.834507 | 85.13 |
| 738dmso_r1 | 738 | cm | dmso | r1 | sw10b | sw11c | 51.50% | 25.00% | 32.651873 | 39.671491 | 0.008494 | 0.868109 | 84.62 |
| 738dmso_r2 | 738 | cm | dmso | r2 | sw10a | sw11d | 49.60% | 67.20% | 51.26617 | 64.717636 | 0.007223 | 0.87646 | 86.6 |
| 738ips_r1 | 738 | ips | ips | r1 | sw10a | sw11c | 49.20% | 50.10% | 79.153512 | 95.016254 | 0.00141 | 0.879315 | 87.85 |
| 738myk0.25_r2 | 738 | cm | myk025 | r2 | sw10b | sw11d | 49.90% | 29.60% | 45.615053 | 58.758739 | 0.025149 | 0.846291 | 80.3 |
| 738omec0.4_r1 | 738 | cm | omec04 | r1 | sw10bmanual | sw11c | 47.50% | 19.80% | 59.329362 | 71.374542 | 0.000899 | 0.897016 | 84.42 |
| 738omec0.4_r2 | 738 | cm | omec04 | r2 | sw10a | sw11d | 50.20% | 43.70% | 50.517001 | 61.703228 | 0.004462 | 0.87506 | 85.25 |
| 811ips_r1 | 811 | ips | ips | r1 | sw12-AA | sw12-AA | 49.10% | 33.20% | 32.410619 | 61.606594 | 0.184381 | 0.67051 | 56.32 |
| 813ips_r1 | 813 | ips | ips | r1 | sw12-AA | sw12-AA | 49.70% | 28.90% | 43.025039 | 73.89741 | 0.143929 | 0.715423 | 61.19 |
| 822Cmyk0.25_r1 | 822 | cm | myk025 | r1 | sw12-BB | sw12-BB | 50.60% | 36.00% | 44.728252 | 61.063068 | 0.011214 | 0.866926 | 82.12 |
| 822Comec0.4_r1 | 822 | cm | omec04 | r1 | sw12-BB | sw12-BB | 50.60% | 46.50% | 104.684055 | 132.374485 | 0.007571 | 0.872123 | 84.28 |
| 822Comec1.0_r1 | 822 | cm | omec10 | r1 | sw12-BB | sw12-BB | 50.10% | 26.20% | 50.338246 | 57.801671 | 0.008045 | 0.868347 | 87.97 |
| 822ips_r1 | 822 | ips | ips | r1 | sw12-BB | sw12-BB | 50.00% | 40.80% | 24.755594 | 74.364587 | 0.258405 | 0.548832 | 35.71 |
| 839ips_r1 | 839 | ips | ips | r1 | sw10a | sw11a | 46.20% | 55.60% | 50.211612 | 58.2215 | 0.00171 | 0.850968 | 90 |
| 860Fdmso_r1 | 860 | cm | dmso | r1 | sw12-BB | sw12-BB | 51.50% | 38.70% | 37.924798 | 49.999342 | 0.010814 | 0.850318 | 82.65 |
| 860Fmyk0.25_r1 | 860 | cm | myk025 | r1 | sw12-BB | sw12-BB | 50.70% | 40.20% | 53.76336 | 68.509591 | 0.009013 | 0.847251 | 83.4 |
| 860ips_r1 | 860 | ips | ips | r1 | sw12-BB | sw12-BB | 50.30% | 32.20% | 170.18782 | 221.710547 | 0.008575 | 0.852347 | 80.78 |
| 861ips_r1 | 861 | ips | ips | r1 | sw12-AA | sw12-AA | 50.70% | 19.10% | 27.825498 | 70.103303 | 0.245267 | 0.591319 | 41.16 |
| 862Edmso_r1 | 862 | cm | dmso | r1 | sw12-BB | sw12-BB | 51.70% | 35.80% | 49.14476 | 69.426492 | 0.009327 | 0.83214 | 79.25 |
| 862Emyk0.25_r1 | 862 | cm | myk025 | r1 | sw12-BB | sw12-BB | 51.20% | 35.50% | 45.578683 | 60.211293 | 0.012466 | 0.830831 | 81.33 |
| 862Eomec0.4_r1 | 862 | cm | omec04 | r1 | sw12-BB | sw12-BB | 52.20% | 44.50% | 57.272998 | 79.598134 | 0.008543 | 0.837881 | 78.75 |
| 862Eomec1.0_r1 | 862 | cm | omec10 | r1 | sw12-BB | sw12-BB | 52.40% | 42.10% | 41.769031 | 59.520132 | 0.011956 | 0.834402 | 77.97 |
| 868ips_r1 | 868 | ips | ips | r1 | sw12-AA | sw12-AA | 49.90% | 25.40% | 31.63751 | 61.667255 | 0.214488 | 0.634313 | 52.55 |
| 869Edmso_r1 | 869 | cm | dmso | r1 | sw12-BB | sw12-BB | 50.50% | 28.80% | 53.243121 | 63.832447 | 0.013729 | 0.86275 | 84.5 |

| | | | | | | | | | | | | | |
|---|---|---|---|---|---|---|---|---|---|---|---|---|---|
| 869ips_r1 | 869 | ips | ips | r1 | sw12-BB | sw12-BB | 50.40% | 19.10% | 55.897001 | 73.185766 | 0.013641 | 0.821155 | 80.38 |
| 885ips_r1 | 885 | ips | ips | r1 | sw12-AA | sw12-AA | 50.80% | 19.90% | 21.241841 | 49.909702 | 0.221234 | 0.625899 | 44.25 |
| 910Cdmso_r1 | 910 | cm | dmso | r1 | sw12-BB | sw12-BB | 51.50% | 31.90% | 43.678077 | 56.974965 | 0.010971 | 0.855925 | 82.03 |
| 910Cmyk0.25_r1 | 910 | cm | myk025 | r1 | sw12-BB | sw12-BB | 50.70% | 34.60% | 41.586788 | 54.977407 | 0.010446 | 0.851081 | 81.98 |
| 910Comec0.4_r1 | 910 | cm | omec04 | r1 | sw12-BB | sw12-BB | 51.50% | 36.00% | 51.887818 | 66.943957 | 0.012362 | 0.855771 | 82.56 |
| 910Comec1.0_r1 | 910 | cm | omec10 | r1 | sw12-BB | sw12-BB | 51.70% | 33.80% | 46.276577 | 63.318577 | 0.014242 | 0.860214 | 80.37 |
| 910ips_r1 | 910 | ips | ips | r1 | sw12-BB | sw12-BB | 50.00% | 17.00% | 42.625376 | 57.560876 | 0.01261 | 0.841673 | 81.11 |
| 915ips_r1 | 915 | ips | ips | r1 | sw12-AA | sw12-AA | 49.20% | 25.00% | 38.216306 | 64.140273 | 0.175175 | 0.685573 | 61.17 |
| 920ips_r1 | 920 | ips | ips | r1 | sw12-AA | sw12-AA | 50.70% | 17.70% | 31.221033 | 65.080985 | 0.232974 | 0.60958 | 49.28 |
| 925Edmso_r1 | 925 | cm | dmso | r1 | sw12-BB | sw12-BB | 51.60% | 35.40% | 44.993495 | 60.762792 | 0.010032 | 0.795571 | 80.82 |
| 925ips_r1 | 925 | ips | ips | r1 | sw12-AA | sw12-AA | 49.20% | 28.60% | 42.26725 | 74.550692 | 0.200877 | 0.673319 | 58.1 |
| 928Admso_r1 | 928 | cm | dmso | r1 | sw12-BB | sw12-BB | 50.40% | 25.70% | 36.588715 | 47.720285 | 0.008961 | 0.813102 | 83.95 |
| 928Amyk0.25_r1 | 928 | cm | myk025 | r1 | sw12-BB | sw12-BB | 51.40% | 42.90% | 48.010101 | 64.1693 | 0.011153 | 0.835782 | 81.93 |
| 928Aomec0.4_r1 | 928 | cm | omec04 | r1 | sw12-BB | sw12-BB | 51.30% | 36.60% | 39.413656 | 54.280726 | 0.008885 | 0.83421 | 81.99 |
| 928Aomec1.0_r1 | 928 | cm | omec10 | r1 | sw12-BB | sw12-BB | 50.90% | 38.00% | 43.043805 | 54.93192 | 0.011333 | 0.813143 | 83.91 |
| 928ips_r1 | 928 | ips | ips | r1 | sw12-BB | sw12-BB | 49.50% | 22.70% | 41.773474 | 56.922112 | 0.010019 | 0.849762 | 81.66 |
| 933dmso_r1 | 933 | cm | dmso | r1 | sw10a | sw11d | 50.00% | 51.70% | 90.331204 | 107.471273 | 0.003977 | 0.875531 | 87.53 |
| 933dmso_r2 | 933 | cm | dmso | r2 | sw10b | sw11b | 49.70% | 23.70% | 32.461452 | 38.864347 | 0.013011 | 0.858443 | 85.76 |
| 933ips_r1 | 933 | ips | ips | r1 | sw09 | sw11d | 49.30% | 21.90% | 40.12219 | 53.855654 | 0.012135 | 0.835803 | 81.38 |
| 933ips_r2 | 933 | ips | ips | r2 | sw10b | sw11b | 50.40% | 25.70% | 52.653818 | 66.036729 | 0.023222 | 0.803677 | 81.53 |
| 933myk0.25_r1 | 933 | cm | myk025 | r1 | sw10a | sw11d | 49.20% | 65.20% | 34.29313 | 46.880921 | 0.002169 | 0.812693 | 84.08 |
| 933myk0.25_r2 | 933 | cm | myk025 | r2 | sw10b | sw11b | 49.10% | 27.70% | 46.030926 | 55.830131 | 0.013622 | 0.856508 | 85.44 |
| 933omec0.4_r1 | 933 | cm | omec04 | r1 | sw10a | sw11d | 47.50% | 68.30% | 60.760218 | 69.582066 | 0.002145 | 0.865542 | 89.87 |
| 933omec0.4_r2 | 933 | cm | omec04 | r2 | sw10b | sw11b | 49.00% | 29.50% | 53.896007 | 64.906411 | 0.013799 | 0.865634 | 85.62 |
| 934dmso_r1 | 934 | cm | dmso | r1 | sw09 | sw11d | 51.00% | 30.30% | 80.701729 | 124.420327 | 0.039828 | 0.806093 | 71.31 |
| 934ips_r1 | 934 | ips | ips | r1 | sw09 | sw11d | 50.10% | 24.30% | 56.048316 | 76.4074 | 0.014836 | 0.832472 | 80.16 |
| 934myk0.25_r1 | 934 | cm | myk025 | r1 | sw09 | sw11d | 50.00% | 33.70% | 65.795331 | 88.630753 | 0.00966 | 0.849173 | 82.92 |
| 934omec0.4_r1 | 934 | cm | omec04 | r1 | sw09 | sw11d | 52.10% | 34.60% | 81.513133 | 111.689115 | 0.030739 | 0.820937 | 79.95 |
| 936dmso_r1 | 936 | cm | dmso | r1 | sw09 | sw11b | 52.20% | 40.40% | 89.071204 | 121.116097 | 0.01371 | 0.855445 | 80.99 |
| 936myk0.25_r1 | 936 | cm | myk025 | r1 | sw10b | sw11b | 49.60% | 33.30% | 60.224503 | 72.852274 | 0.01015 | 0.862553 | 85.67 |
| 936omec0.4_r1 | 936 | cm | omec04 | r1 | sw09 | sw11b | 49.10% | 31.40% | 54.78249 | 71.402359 | 0.008336 | 0.856432 | 84.06 |
| 941dmso_r1 | 941 | cm | dmso | r1 | sw09 | sw11c | 51.60% | 20.00% | 34.874449 | 47.369961 | 0.053303 | 0.817813 | 79 |
| 941ips_r1 | 941 | ips | ips | r1 | sw09 | sw11c | 49.20% | 18.40% | 37.509454 | 47.745372 | 0.028966 | 0.836579 | 82.49 |
| 941myk0.25_r1 | 941 | cm | myk025 | r1 | sw09 | sw11c | 50.30% | 27.50% | 93.463333 | 123.636933 | 0.039274 | 0.808138 | 82.55 |
| 941omec0.4_r1 | 941 | cm | omec04 | r1 | sw09 | sw11c | 51.20% | 23.90% | 42.737024 | 56.789481 | 0.022587 | 0.85508 | 83.06 |
| 942dmso_r1 | 942 | cm | dmso | r1 | sw10b | sw11a | 49.20% | 22.50% | 24.61892 | 29.56304 | 0.012181 | 0.848696 | 86.87 |
| 942dmso_r2 | 942 | cm | dmso | r2 | sw10a | sw11d | 50.50% | 62.30% | 38.650471 | 50.802469 | 0.002111 | 0.849408 | 85.7 |
| 942ips_r1 | 942 | ips | ips | r1 | sw09 | sw11b | 50.70% | 19.60% | 56.382981 | 73.186254 | 0.031119 | 0.833515 | 81.96 |
| 942myk0.25_r1 | 942 | cm | myk025 | r1 | sw10b | sw11a | 49.60% | 20.50% | 20.653616 | 24.499216 | 0.01292 | 0.830298 | 86.39 |
| 942myk0.25_r2 | 942 | cm | myk025 | r2 | sw10a | sw11d | 49.30% | 71.20% | 25.526134 | 33.202759 | 0.002717 | 0.854353 | 86.17 |
| 942omec0.4_r1 | 942 | cm | omec04 | r1 | sw10b | sw11a | 49.60% | 23.20% | 39.697696 | 45.791509 | 0.00606 | 0.875322 | 88.58 |
| 942omec0.4_r2 | 942 | cm | omec04 | r2 | sw10a | sw11d | 50.70% | 61.90% | 43.183087 | 57.426746 | 0.003209 | 0.85916 | 84.63 |
| 944dmso_r1 | 944 | cm | dmso | r1 | sw09 | sw11b | 50.90% | 25.60% | 52.718014 | 68.047515 | 0.023084 | 0.851761 | 83.92 |
| 944ips_r1 | 944 | ips | ips | r1 | sw09 | sw11b | 51.00% | 18.50% | 46.380174 | 59.003471 | 0.024055 | 0.828933 | 82.84 |
| 944myk0.25_r1 | 944 | cm | myk025 | r1 | sw09 | sw11b | 51.00% | 20.70% | 31.706497 | 41.24733 | 0.017542 | 0.844968 | 83.04 |
| 954dmso_r1 | 954 | cm | dmso | r1 | sw09 | sw11a | 50.30% | 23.40% | 38.482989 | 49.794042 | 0.011187 | 0.866292 | 83.98 |
| 954ips_r1 | 954 | ips | ips | r1 | sw09 | sw11b | 49.50% | 26.60% | 60.562388 | 76.53879 | 0.008831 | 0.837715 | 85.07 |
| 954myk0.25_r1 | 954 | cm | myk025 | r1 | sw09 | sw11a | 49.40% | 24.60% | 38.28839 | 50.1982 | 0.016225 | 0.849285 | 84.26 |
| 954omec0.4_r1 | 954 | cm | omec04 | r1 | sw09 | sw11a | 50.50% | 27.60% | 43.712227 | 56.748144 | 0.01363 | 0.861321 | 83.21 |
| 955dmso_r1 | 955 | cm | dmso | r1 | sw10a | sw11c | 50.90% | 53.10% | 65.937913 | 80.262925 | 0.006857 | 0.85857 | 85.68 |
| 955dmso_r2 | 955 | cm | dmso | r2 | sw10b | sw11d | 50.00% | 23.20% | 26.288338 | 31.540819 | 0.011718 | 0.841826 | 85.95 |
| 955ips_r1 | 955 | ips | ips | r1 | sw09 | sw11c | 50.20% | 22.00% | 42.805 | 55.301604 | 0.015295 | 0.827678 | 82.17 |
| 955ips_r2 | 955 | ips | ips | r2 | sw10b | sw11d | 49.60% | 21.20% | 35.239993 | 41.352355 | 0.007071 | 0.858839 | 86.79 |
| 955myk0.25_r1 | 955 | cm | myk025 | r1 | sw10a | sw11c | 50.20% | 40.90% | 39.38658 | 51.303886 | 0.005591 | 0.825298 | 83.91 |
| 955myk0.25_r2 | 955 | cm | myk025 | r2 | sw10b | sw11d | 50.80% | 30.90% | 45.523971 | 54.825264 | 0.015465 | 0.844457 | 84.84 |
| 955omec0.4_r1 | 955 | cm | omec04 | r1 | sw10a | sw11c | 50.20% | 49.40% | 67.0139 | 80.848791 | 0.004533 | 0.860887 | 86.04 |
| 959dmso_r1 | 959 | cm | dmso | r1 | sw09 | sw11d | 50.90% | 28.10% | 43.175692 | 56.671542 | 0.010061 | 0.855995 | 83.35 |
| 959ips_r1 | 959 | ips | ips | r1 | sw09 | sw11d | 50.00% | 20.90% | 31.66314 | 41.545426 | 0.022139 | 0.785869 | 81.23 |
| 959myk0.25_r1 | 959 | cm | myk025 | r1 | sw09 | sw11d | 51.50% | 26.70% | 45.472763 | 61.075545 | 0.01974 | 0.838685 | 81.29 |
| 959omec0.4_r1 | 959 | cm | omec04 | r1 | sw09 | sw11d | 51.40% | 23.80% | 43.400474 | 57.229774 | 0.019192 | 0.84834 | 82.37 |
| 960dmso_r1 | 960 | cm | dmso | r1 | sw10b | sw11c | 49.60% | 28.00% | 57.123358 | 67.172857 | 0.009552 | 0.864758 | 86.82 |
| 960ips_r1 | 960 | ips | ips | r1 | sw09 | sw11c | 50.90% | 20.80% | 50.776399 | 66.964559 | 0.025706 | 0.822164 | 79.42 |
| 960myk0.25_r1 | 960 | cm | myk025 | r1 | sw09 | sw11c | 49.20% | 23.30% | 28.553729 | 39.600059 | 0.022072 | 0.829964 | 80.43 |
| 960omec0.4_r1 | 960 | cm | omec04 | r1 | sw09 | sw11c | 56.00% | 33.10% | 66.531364 | 98.498987 | 0.050242 | 0.772864 | 73.8 |
| 961dmso_r1 | 961 | cm | dmso | r1 | sw09 | sw11c | 51.00% | 25.10% | 55.480941 | 75.131766 | 0.01402 | 0.85308 | 81.65 |
| 961ips_r1 | 961 | ips | ips | r1 | sw09 | sw11c | 49.80% | 21.90% | 50.226278 | 66.670401 | 0.0045 | 0.867304 | 82.61 |
| 961myk0.25_r1 | 961 | cm | myk025 | r1 | sw09 | sw11c | 50.70% | 23.90% | 43.973116 | 60.260568 | 0.013017 | 0.836215 | 80.37 |
| 962dmso_r1 | 962 | cm | dmso | r1 | sw09 | sw11a | 51.40% | 27.10% | 43.137084 | 61.38376 | 0.038829 | 0.827076 | 78.06 |
| 962ips_r1 | 962 | ips | ips | r1 | sw09 | sw11a | 51.40% | 17.50% | 48.523349 | 66.435223 | 0.0433 | 0.804861 | 77 |
| 962myk0.25_r1 | 962 | cm | myk025 | r1 | sw09 | sw11a | 51.40% | 21.90% | 24.230398 | 33.832859 | 0.036009 | 0.830076 | 80.17 |
| 962omec0.4_r1 | 962 | cm | omec04 | r1 | sw09 | sw11a | 51.00% | 26.40% | 35.556494 | 46.605905 | 0.04619 | 0.816915 | 81.08 |
| 963dmso_r1 | 963 | cm | dmso | r1 | sw10a | sw11c | 50.90% | 55.10% | 60.73686 | 78.269383 | 0.003666 | 0.884298 | 85.2 |
| 963dmso_r2 | 963 | cm | dmso | r2 | sw10b | sw11d | 50.40% | 26.40% | 46.701675 | 56.790445 | 0.012412 | 0.8462 | 84.11 |
| 963ips_r1 | 963 | ips | ips | r1 | sw10a | sw11c | 50.50% | 42.20% | 57.137646 | 76.353866 | 0.001541 | 0.829061 | 82.28 |
| 963myk0.25_r1 | 963 | cm | myk025 | r1 | sw10a | sw11c | 47.70% | 69.60% | 35.692282 | 45.266026 | 0.002761 | 0.86082 | 87.71 |

| | | | | | | | | | | | | | |
|---|---|---|---|---|---|---|---|---|---|---|---|---|---|
| 963myk0.25_r2 | 963 | cm | myk025 | r2 | sw10b | sw11d | 49.70% | 22.00% | 22.111615 | 26.285255 | 0.01109 | 0.852244 | 85.7 |
| 963omec0.4_r1 | 963 | cm | omec04 | r1 | sw10a | sw11c | 50.40% | 31.20% | 34.940907 | 41.18409 | 0.003721 | 0.890664 | 88.2 |
| 963omec0.4_r2 | 963 | cm | omec04 | r2 | sw10b | sw11d | 50.40% | 24.70% | 33.288005 | 40.346557 | 0.011703 | 0.847246 | 84.52 |
| 964dmso_r1 | 964 | cm | dmso | r1 | sw09 | sw11b | 52.60% | 21.80% | 43.773898 | 65.349562 | 0.050489 | 0.809632 | 75.94 |
| 964ips_r1 | 964 | ips | ips | r1 | sw09 | sw11b | 50.50% | 22.60% | 47.333094 | 67.930262 | 0.05183 | 0.798486 | 78.37 |
| 965dmso_r2 | 965 | cm | dmso | r2 | sw10b | sw11b | 49.60% | 25.70% | 47.482416 | 55.860182 | 0.010142 | 0.867442 | 87 |
| 965ips_r1 | 965 | ips | ips | r1 | sw09 | sw11a | 50.60% | 22.40% | 55.16036 | 75.225362 | 0.026848 | 0.819562 | 80.64 |
| 965ips_r2 | 965 | ips | ips | r2 | sw10b | sw11b | 49.50% | 31.00% | 47.092242 | 55.981034 | 0.004132 | 0.83189 | 87.38 |
| 965myk0.25_r2 | 965 | cm | myk025 | r2 | sw10b | sw11b | 48.40% | 28.40% | 35.118295 | 41.137446 | 0.009996 | 0.852603 | 87.98 |
| 965omec0.4_r2 | 965 | cm | omec04 | r2 | sw10b | sw11b | 48.10% | 31.00% | 62.655682 | 73.375423 | 0.007873 | 0.876323 | 88.56 |
| 967Bdmso_r2 | 967 | cm | dmso | r2 | sw12-BB | sw12-BB | 49.90% | 51.80% | 50.83429 | 63.325203 | 0.009446 | 0.860592 | 86.57 |
| 967dmso_r1 | 967 | cm | dmso | r1 | sw09 | sw11c | 50.70% | 26.30% | 38.652378 | 55.004744 | 0.027177 | 0.846766 | 80.14 |
| 967ips_r1 | 967 | ips | ips | r1 | sw10a | sw11c | 49.90% | 33.10% | 48.975063 | 58.272786 | 0.002759 | 0.870268 | 87.03 |
| 969dmso_r1 | 969 | cm | dmso | r1 | sw10a | sw11d | 51.50% | 37.90% | 70.811421 | 87.934329 | 0.009219 | 0.860406 | 82.83 |
| 969ips_r1 | 969 | ips | ips | r1 | sw10a | sw11d | 49.90% | 26.90% | 44.196039 | 53.789977 | 0.004731 | 0.8493 | 85.39 |
| 969myk0.25_r1 | 969 | cm | myk025 | r1 | sw10a | sw11d | 49.80% | 44.50% | 61.653779 | 77.383702 | 0.004608 | 0.860484 | 84.72 |
| 969omec0.4_r1 | 969 | cm | omec04 | r1 | sw10a | sw11d | 49.90% | 35.40% | 49.502374 | 59.245531 | 0.004992 | 0.862872 | 84.94 |
| 979dmso_r1 | 979 | cm | dmso | r1 | sw09 | sw11c | 49.80% | 22.80% | 33.992469 | 46.191378 | 0.012388 | 0.840882 | 84.77 |
| 979ips_r1 | 979 | ips | ips | r1 | sw10b | sw11c | 49.10% | 30.90% | 59.699306 | 72.238395 | 0.013072 | 0.803605 | 85.28 |
| 986dmso_r1 | 986 | cm | dmso | r1 | sw10b | sw11b | 48.40% | 53.30% | 69.883995 | 84.947442 | 0.009347 | 0.848277 | 86.15 |
| 986dmso_r2 | 986 | cm | dmso | r2 | sw10b | sw11d | 50.90% | 32.90% | 61.216227 | 77.910799 | 0.013022 | 0.860556 | 81.98 |
| 986ips_r1 | 986 | ips | ips | r1 | sw09 | sw11c | 49.50% | 30.50% | 61.011119 | 86.389275 | 0.016138 | 0.826018 | 81.65 |
| 986myk0.25_r1 | 986 | cm | myk025 | r1 | sw10a | sw11b | 50.40% | 43.40% | 56.781781 | 70.183155 | 0.005943 | 0.858414 | 85.44 |
| 986omec0.4_r1 | 986 | cm | omec04 | r1 | sw10a | sw11b | 50.30% | 51.10% | 73.725251 | 95.481457 | 0.006099 | 0.87171 | 84.89 |
| 989dmso_r1 | 989 | cm | dmso | r1 | sw10a | sw11a | 50.60% | 47.40% | 70.263042 | 84.80663 | 0.004988 | 0.888873 | 86.44 |
| 989ips_r1 | 989 | ips | ips | r1 | sw10a | sw11a | 50.00% | 30.50% | 62.693511 | 77.07226 | 0.001975 | 0.864355 | 86.43 |
| 990dmso_r1 | 990 | cm | dmso | r1 | sw10a | sw11a | 49.50% | 58.80% | 51.047543 | 60.695379 | 0.003921 | 0.865972 | 88.14 |
| 990ips_r1 | 990 | ips | ips | r1 | sw09 | sw11a | 49.90% | 22.00% | 40.990684 | 58.794546 | 0.026723 | 0.807367 | 79.96 |
| 990ips_r2 | 990 | ips | ips | r2 | sw10b | sw11b | 49.00% | 26.70% | 51.991494 | 62.007001 | 0.009326 | 0.842178 | 86.45 |
| 990myk0.25_r1 | 990 | cm | myk025 | r1 | sw10a | sw11a | 50.20% | 51.90% | 68.781018 | 83.500926 | 0.025379 | 0.828753 | 84.99 |
| 990omec0.4_r1 | 990 | cm | omec04 | r1 | sw10a | sw11a | 49.90% | 59.70% | 45.494853 | 56.530121 | 0.004053 | 0.873537 | 87.33 |
| 991dmso_r1 | 991 | cm | dmso | r1 | sw10a | sw11d | 51.00% | 44.70% | 34.604534 | 44.361708 | 0.01854 | 0.857743 | 84.85 |
| 991dmso_r2 | 991 | cm | dmso | r2 | sw09 | sw11c | 49.90% | 34.40% | 46.232019 | 66.978689 | 0.01016 | 0.847927 | 81.28 |
| 991ips_r1 | 991 | ips | ips | r1 | sw09 | sw11c | 51.70% | 17.10% | 24.228882 | 34.048854 | 0.032678 | 0.815318 | 79.08 |
| 991myk0.25_r1 | 991 | cm | myk025 | r1 | sw10a | sw11d | 50.20% | 45.40% | 53.783544 | 64.664584 | 0.005193 | 0.875435 | 87.56 |
| 991omec0.4_r1 | 991 | cm | omec04 | r1 | sw10a | sw11d | 50.10% | 58.10% | 74.070874 | 92.693239 | 0.003579 | 0.881636 | 86.41 |
| 995Cdmso_r1 | 995 | cm | dmso | r1 | sw12-BB | sw12-BB | 51.40% | 34.70% | 36.489949 | 54.068112 | 0.010536 | 0.853912 | 77.54 |
| 995Cmyk0.25_r1 | 995 | cm | myk025 | r1 | sw12-BB | sw12-BB | 50.40% | 27.20% | 43.487809 | 51.886334 | 0.010185 | 0.811971 | 85.51 |
| 995Comec0.4_r1 | 995 | cm | omec04 | r1 | sw12-BB | sw12-BB | 51.30% | 33.90% | 38.587027 | 52.926634 | 0.009863 | 0.823739 | 80.91 |
| 995Comec1.0_r1 | 995 | cm | omec10 | r1 | sw12-BB | sw12-BB | 52.00% | 41.80% | 41.956251 | 57.797046 | 0.007665 | 0.852808 | 80.65 |
